# Systemic control and inter-organ communication of ovarian aging

**DOI:** 10.64898/2026.10.11.758357

**Authors:** Sharif Iqbal, Alexander Tyshkovskiy, Bohan Zhang, Sirui Zhang, Adrian Molière, Kejun Ying, Yunhao Bai, Harlan Stevens, Ludger Goeminne, Juan José Alba-Linares, Erik Jacques, Amin Hossain, Dmitrii Glubokov, Alibek Moldakozhayev, Jesse R. Poganik, Mahdi Moqri, Nathalie Agar, Fei Chen, Steve Horvath, Vadim N. Gladyshev

## Abstract

Whether ovarian aging is intrinsic and unidirectional or dynamically regulated by systemic context remains an open question for fertility and women’s healthspan. To address it, we developed a mouse ovarian transcriptomic aging clock (OvaAge) and applied it, together with published transcriptomic, epigenetic and pathway-resolved DNA methylation (PathAge) clocks, multi-omic analyses and reproductive outcomes, to new reciprocal heterochronic ovary transplantation and parabiosis experiments. Ovarian molecular age shifted in both directions: aged ovaries became younger in young hosts and young ovaries became older in aged hosts, and in each case the shift was partial across transcriptomic, epigenetic and pathway-resolved measures. Ovaries from 13-month-old C57BL/6J donors supported full-term pregnancies and live births at 17-18 months of age when transplanted into young hosts, but not when transplanted into old hosts, and young ovaries supported live births in 17-18-month-old recipients. Donor ovary age altered the transcriptomic age of the uterus in both directions, while young ovaries also lowered the transcriptomic age of aged liver and brain, identifying the ovary as an active node within an inter-organ aging network. Across unilateral and bilateral transplantation and parabiosis, an older environment increased inflammatory gene expression in the ovary, whereas a younger environment reduced it and increased mitochondrial and biosynthetic programs. Integration of methylation, expression and cell-type markers linked these changes to declining oocyte-associated programs and an expanding stromal compartment enriched for multinucleated giant cells (MNGCs). MNGC-rich regions formed a reversibly remodeled, focally aged niche with distinct spatial metabolic features, and a related macrophage expression signature was detected in human ovaries. Together, these findings identify ovarian aging as a modular, bidirectionally plastic process that is regulated by the systemic environment and point to tractable intervention points.

## Introduction

Ovarian reproductive function depends on a finite follicle reserve that progressively declines with age (Faddy *et al*, 1992; Lei & Spradling, 2013; Wallace & Kelsey, 2010). Across the reproductive lifespan, progressive follicle depletion is accompanied by stromal fibrosis, immune remodeling, endocrine dysregulation, and disrupted intercellular signaling that accompany declining oocyte competence and fertility (Amargant *et al*, 2020; Balough *et al*, 2024; Isola *et al*, 2024; Lan *et al*, 2026). Recent single-cell and spatial atlases have resolved these changes across ovarian cell types, revealing stress- and DNA damage-associated signatures in human oocytes (Wu *et al*, 2024), changes in cellular composition (Yan *et al*, 2023), and remodeling of follicular and stromal niches in primates (Lu *et al*, 2024). Emerging evidence suggests that ovarian aging is also spatially organized, with multinucleated giant cells (MNGCs) accumulating in stromal regions and displaying immune and degradation-associated molecular signatures (Converse *et al*, 2025). These observations raise the possibility that localized differences in damage processing and metabolism contribute to ovarian decline.

Follicle number alone, however, does not determine reproductive competence (Wilkosz *et al*, 2014). Fertility also depends on oocyte quality, support from follicular somatic cells, neuroendocrine coordination, and the capacity of the uterus to sustain pregnancy (Woods *et al*, 2017). Experiments placing aged mouse oocytes within young reconstructed follicles have improved maturation and subsequent developmental outcomes (Wang *et al*, 2024). Replenishing NAD+ or limiting its depletion through CD38 inhibition can also preserve aspects of ovarian function in aged mice (Bertoldo *et al*, 2020; Yang *et al*, 2024). These findings indicate that some features of reproductive decline remain responsive to the surrounding tissue and its metabolic state. Mice do not undergo menopause, but C57BL/6J females become irregular and then stop cycling during the second year of life (Felicio *et al*, 1986).

At the level of the whole organism, ovarian transplantation provides evidence for reciprocal influences between the ovary and its host. Young ovarian grafts can extend reproductive cycling in middle-aged mice, although recipient age still limits the response (Felicio *et al*, 1986). Other transplantation studies have reported longer lifespan and reduced cardiac pathology (Cargill *et al*, 2003; Mason *et al*, 2009; Mason *et al*, 2011). Benefits to longevity and inflammation after transplantation of germ cell-depleted ovaries further implicate ovarian somatic tissue in these effects (Habermehl *et al*, 2019). Human studies have also associated premenopausal oophorectomy with cognitive impairment and early bilateral oophorectomy with cardiovascular mortality (Rivera *et al*, 2009; Rocca *et al*, 2007). Although surgical removal differs from natural ovarian aging, these findings link loss of ovarian function to health beyond reproduction.

These cellular, spatial and transplantation studies have advanced our understanding of ovarian aging, yet they leave a central mechanistic gap. In other organs, heterochronic parabiosis has shown that tissue aging reflects both cell-intrinsic and systemic influences (Conboy *et al*, 2005; Zhang *et al*, 2023), but the contribution of systemic signals to ovarian aging is unknown. How do systemic aging signals accelerate or restrain ovarian decline? How do follicle reserve dynamics interact with the ovarian microenvironment and maternal physiology to determine reproductive competence? Are age-associated inflammatory and metabolic changes concentrated in specific somatic and follicular regions, and can their distribution change with host age? And does ovarian aging passively reflect organismal aging, or does it actively contribute to aging in other organs?

Addressing these questions requires quantitative tools that relate ovarian molecular state to reproductive function and distinguish the effects of ovarian and host age. Here, we develop OvaAge, an ovary-specific transcriptomic aging clock, and combine it with transcriptomic (Tyshkovskiy *et al*, 2026) and epigenetic (Lu *et al*, 2023) age estimators and with pathway-resolved DNA methylation models (PathAge and MetroAge) (Ying *et al*, 2024). We apply these tools to new reciprocal heterochronic ovary transplantation experiments, in which one or both ovaries were replaced, and to new heterochronic parabiosis experiments. Shared circulation provides a complementary test of systemic influences while retaining the ovary in its native environment (Conboy *et al*, 2005; Zhang *et al*, 2023). We combine these interventions with transcriptomic profiles of the brain, pituitary, liver and uterus, spatial and circulating metabolomics, and assays of mating, pregnancy and live birth. By linking molecular responses to reproductive outcomes, we test whether ovarian age responds to host age in both directions, whether late-life ovarian decline is partially reversible, and whether changing ovarian age affects aging in other organs.

## Results

### Ovarian aging combines loss of follicular programs with immune and stromal remodeling

Existing transcriptomic clocks (Tyshkovskiy *et al*, 2026) are trained on heterogeneous somatic tissues and lack resolution for the ovary, whose endocrine cyclicity, follicular dynamics, and stromal heterogeneity create transcriptional features absent from canonical aging models. We therefore first developed an ovary-specific transcriptomic clock and benchmarked it against epigenetic and multi-tissue aging metrics (**Fig. 1**). Ovaries were collected from a female mouse cohort spanning postnatal maturation through late reproductive aging (0.5-17 months) and profiled by bulk RNA-seq (**Fig. 1A**). The clock was trained by elastic net regression with chronological age as the outcome. In leave-one-out cross-validation, predicted age correlated with chronological age (r = 0.96; mean absolute error, 1.1 months) across development and adulthood (**Fig. 1B**).

**Figure 1.**
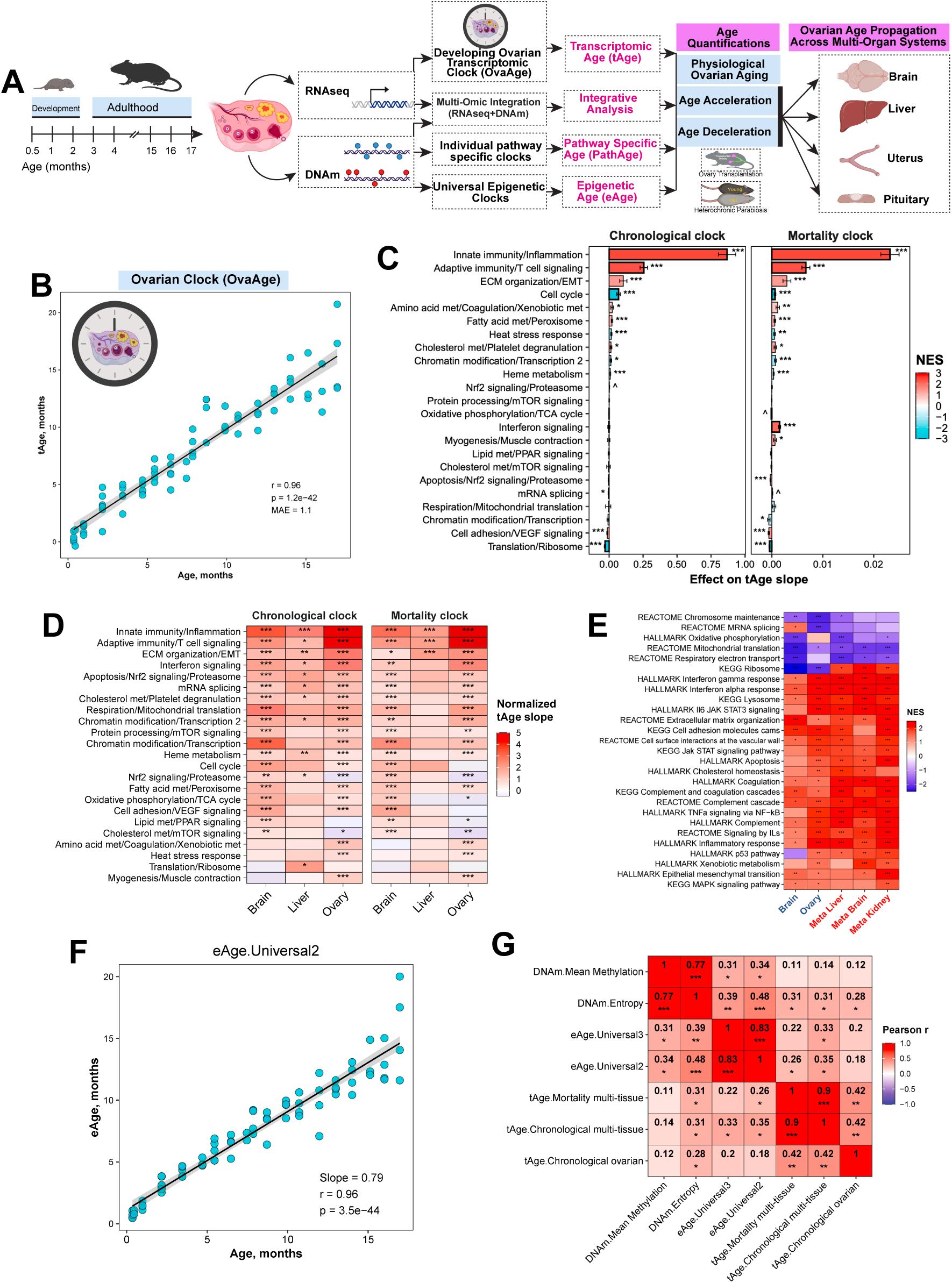
An ovary-specific transcriptomic clock and multi-omic measures of ovarian aging. (A) Study design. Ovaries collected between 0.5 and 17 months of age were profiled by bulk RNA sequencing and DNA methylation (DNAm). These data were used to build an ovary-specific transcriptomic clock (OvaAge) and to apply pathway-specific (PathAge) and universal (eAge) methylation clocks. The resulting age measures were then applied to heterochronic ovary transplantation and heterochronic parabiosis, and to brain, liver, uterus and pituitary of transplant recipients. (B) OvaAge, the ovary-specific elastic net clock developed in this study. Predicted transcriptomic age (tAge, months) is plotted against chronological age; r = 0.96, P = 1.2 × 10^−42^, mean absolute error (MAE) = 1.1 months. Values are leave-one-out cross-validated predictions. (C) Contribution of each co-regulated gene module defined by Tyshkovskiy et al (2026) to the ovarian tAge slope, calculated for the rodent multi-tissue clocks of chronological age (left) and expected mortality (right). Bar length shows the module’s effect on the tAge slope (± s.e.). Bar color shows the module-level enrichment score in the key, with positive values indicating that the module’s genes mainly increase with age. Asterisks show Benjamini-Hochberg (BH)-adjusted significance of each module effect; ^P < 0.1. Modules are ordered by their effect in the chronological clock. (D) Normalized aging slopes of module-specific transcriptomic clocks in brain, liver and ovary collected from the same mice. These are module-restricted versions of the YuGene-normalized rodent multi-tissue elastic net clocks of chronological age (left) and expected mortality (right) (Tyshkovskiy *et al*, 2026). Color shows the normalized tAge slope; asterisks show BH-adjusted P values. (E) Pre-ranked gene set enrichment analysis (GSEA; Hallmark, KEGG and Reactome) of age-associated expression in brain and ovary from this cohort, and in liver, brain and kidney aging meta-signatures (Meta Liver, Meta Brain, Meta Kidney) (Tyshkovskiy *et al*, 2026). Genes were ranked by their signed association with age. Color shows the normalized enrichment score (NES); asterisks show BH-adjusted significance. Pathways significant in at least one column are shown. (F) Epigenetic age (eAge, months) from the universal pan-mammalian clock 2 (Universal2) against chronological age (Lu *et al*, 2023); slope = 0.79, Pearson r = 0.96, P = 3.5 × 10^−44^. (G) Partial Pearson correlations, adjusted for chronological age, among ovarian aging measures: mean methylation, methylation entropy, eAge from the Universal3 and Universal2 clocks, tAge from the multi-tissue mortality and chronological clocks (Tyshkovskiy *et al*, 2026), and OvaAge (tAge.Chronological ovarian). Cells show Pearson r and BH-adjusted significance. In B and F, points are individual ovaries and lines show linear fits with 95% confidence bands. *P < 0.05, **P < 0.01, ***P < 0.001.

Gene-level analysis of chronological age slopes revealed genes associated with ovarian aging, with more upregulated than downregulated genes **(Ext. Data Fig. 1A**). The strongest increases were in immunoglobulin genes (*Ighm*, *Igkc*, *Jchain* and *Ighg2c*), the complement gene *C4b* and leukocyte genes such as *Cd53*, *Itgb2* and *Cxcr6*. These are products of B cells, plasma cells and other leukocytes, and their increase agrees with the doubling of immune cells and the shift toward lymphocytes and adaptive immunity found in single-cell studies of the aging mouse ovary (Ben Yaakov *et al*, 2023; Isola *et al*, 2024).

To relate the age predictions to biological processes, we used the co-regulated gene modules defined for the multi-tissue clocks of chronological age and expected mortality (Tyshkovskiy *et al*, 2026). In both clocks, innate immunity and inflammation made the largest positive contribution to the ovarian aging slope, followed by adaptive immune/T cell signaling and extracellular matrix (ECM) organization, whereas cell cycle and translation/ribosome modules contributed negatively (**Fig. 1C**). Module contributions to the mortality-clock slope also differed between development and adulthood (Ext. Data Fig. 1B), and the inflammatory module increased steadily from early postnatal stages through late life (Ext. Data Fig. 1C).

Brain and liver collected from the same animals allowed a direct comparison among organs. Immune, interferon and ECM modules increased with age in all three tissues, as expected from shared transcriptomic signatures of mammalian aging (Tyshkovskiy *et al*, 2026), and their slopes were steepest in the ovary (**Fig. 1D**). The ovary, however, diverged in several respects. The Nrf2 and proteasome module and the cholesterol metabolism and mTOR module declined with age in the ovary but increased in the brain. Interferon, IL6-JAK-STAT3, complement, lysosome and ECM gene sets increased with ovarian age, and ribosome, mRNA splicing and chromosome maintenance gene sets decreased (**Fig. 1E**), consistent with the inflammation, fibrosis and stiffening that build up in the ovarian stroma over repeated cycles of ovulation and repair (Amargant *et al*, 2020; Briley *et al*, 2016; Winkler *et al*, 2024). The multi-tissue chronological clock tracked age less closely in ovary (r = 0.87) than in brain (r = 0.97) or liver (r = 0.92) (**Ext. Data Fig. 1D**), and less closely than OvaAge (r = 0.96). After adjustment for chronological age, its residuals were only weakly correlated across organs of the same animal (Spearman’s ρ = 0.13 to 0.23; **Ext. Data Fig. 1E**). Ovarian aging therefore shares inflammatory features with other tissues but is captured less accurately by a multi-tissue model, which motivated an ovary-specific clock.

We next asked whether OvaAge detects conditions known to alter ovarian aging. In Ames dwarf mice (GSE84078), ovarian tAge was similar to that of wild-type mice in young animals but significantly lower in aged animals (**Ext. Data Fig. 2A**), consistent with the attenuated inflammatory and immune remodeling reported in this longevity model (Schneider *et al*, 2017). In heterochronic follicle reconstitution experiments (GSE270016) (Wang *et al*, 2024), aged oocytes within young follicular environments had lower tAge than those in aged follicles, while young oocytes in aged follicles had higher tAge (**Ext. Data Fig. 2B**). Pseudobulked granulosa cells from aged mice (GSE232309) also had higher tAge than those from young mice (**Ext. Data Fig. 2C**), as did ovarian somatic cells from aged *Macaca fascicularis* (GSE130664; **Ext. Data Fig. 2D**). Among clock modules, only adaptive immune/T cell signaling differed significantly between young and old macaques, contributing to the higher tAge of aged animals (**Ext. Data Fig. 2E**). OvaAge therefore discriminates age across independent datasets, cell compartments and species.

DNA methylation provided an independent measure in the same ovaries. Two universal pan-mammalian DNA methylation clocks (Lu *et al*, 2023) yielded age estimates strongly correlated with chronological age (clock 2, Universal2, r = 0.96, slope 0.79; clock 3, Universal3, r = 0.96, slope 0.86; **Fig. 1F and Ext. Data Fig. 3A**), and both methylation entropy (r = 0.80) and mean methylation (r = 0.74) increased with age (**Ext. Data Fig. 3B,C**). After adjustment for chronological age, age estimates remained strongly correlated between the two universal DNA methylation clocks (r = 0.83) and between the two multi-tissue transcriptomic clocks (r = 0.90). Partial correlations of OvaAge with the multi-tissue transcriptomic clocks were moderate (r = 0.42), and correlations between the methylation clocks and the transcriptomic clocks were weak (r = 0.18-0.35, **Fig. 1G**). Weak correlations among age-adjusted measures of biological aging have also been reported in human cohorts (Jansen *et al*, 2021; Li *et al*, 2020), suggesting that different clocks capture partly distinct aspects of aging. We therefore used transcriptomic and methylation measures side by side in the intervention experiments.

Because follicles are lost with age, part of the bulk signal may reflect a change in cell composition. Deconvolution with a single-cell reference of the mouse ovary (Isola *et al*, 2024) showed that the estimated fractions of oocytes (ρ = -0.82) and of two granulosa cell populations (ρ = -0.54 and -0.50) decreased with age, whereas those of phagocytes (ρ = 0.88), epithelial cells (ρ = 0.64) and luteal cells (ρ = 0.55) increased (**Ext. Data Fig. 4**). Loss of follicular cells and gain of phagocytic and other non-follicular cells therefore contribute to the bulk aging signal. Bulk data cannot resolve how much of the age signal arises within individual cell types, which will require cell type-resolved clocks. We next used transplantation and parabiosis to test whether these features of ovarian aging could be modified by the systemic environment.

### Systemic age context dynamically modulates ovarian biological age

Transplantation of young ovaries into aged mice extends lifespan and reduces cardiovascular and musculoskeletal pathology (Cargill *et al*, 2003; Mason *et al*, 2009; Mason *et al*, 2011; Peterson *et al*, 2016), even when the donor ovaries are depleted of germ cells (Habermehl *et al*, 2019), and ovarian loss in women is associated with accelerated cardiovascular, skeletal and neurological aging (Rocca *et al*, 2007; Rocca *et al*, 2018; Rocca *et al*, 2012). Whether the systemic environment, in turn, modulates the molecular age of the ovary has not been tested. If it does, an aged ovary transplanted into a young host, and vice versa, should shift in molecular age (Fig. 2). We tested this with unilateral orthotopic transplantation, in which each animal carries a transplanted ovary alongside a contralateral endogenous ovary, allowing comparison of two ovaries of different origin within the same systemic environment (**Fig. 2A**). Bilateral transplantation, in which both ovaries were replaced, removed the contralateral ovary and any ovary-ovary interaction (**Fig. 2E**).

**Figure 2.**
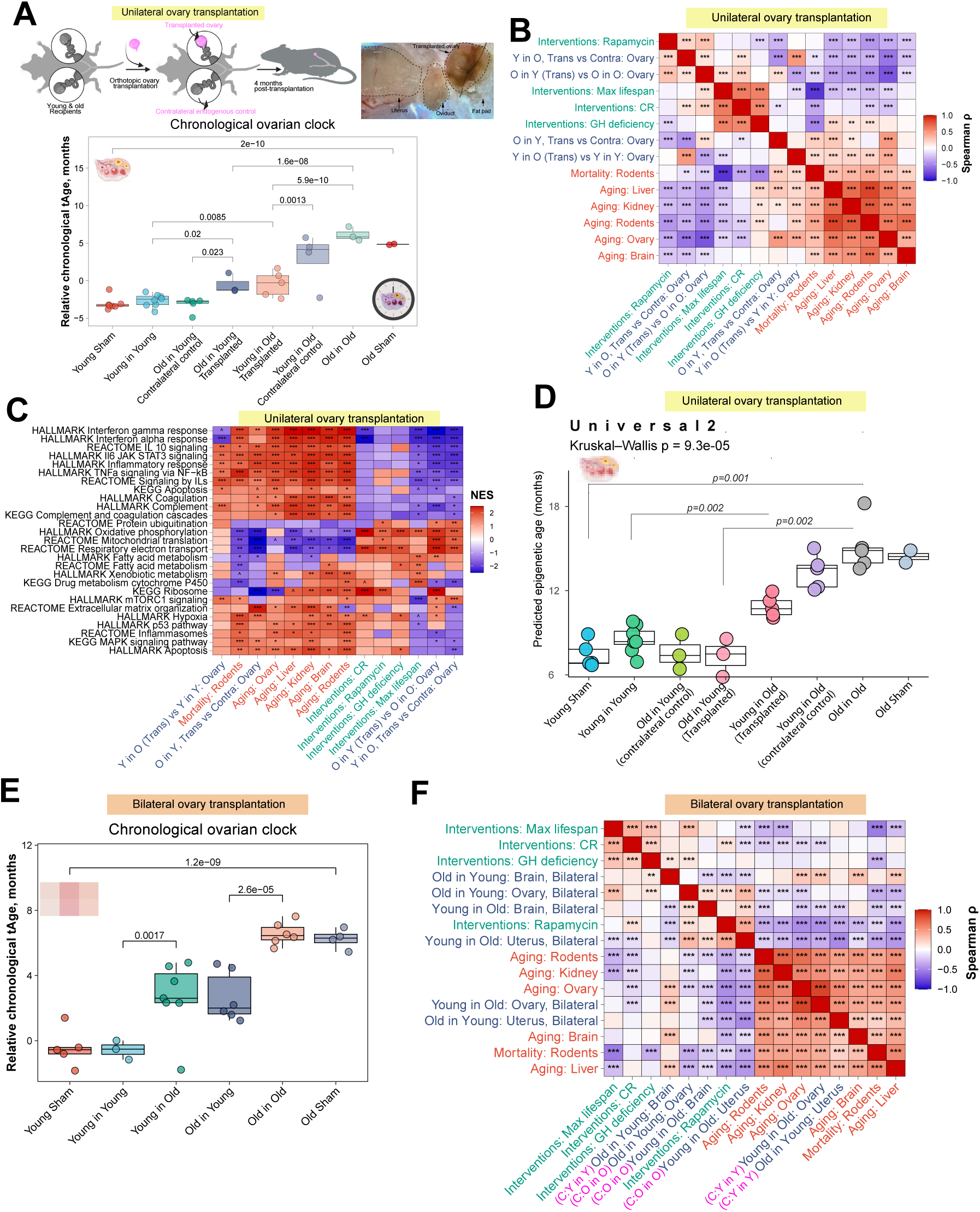
Recipient age shifts the molecular age of transplanted ovaries. (A) Unilateral orthotopic transplantation. A young or old ovary was transplanted into a young or old recipient, the contralateral endogenous ovary was retained, and tissues were collected 4 months later. The photograph shows a graft in situ, with uterus, oviduct, graft and fat pad outlined. Lower panel: OvaAge (Bayesian ridge model; relative chronological tAge, months) for Young Sham, Y in Y, O in Y (contralateral control), O in Y (transplanted), Y in O (transplanted), Y in O (contralateral control), O in O and Old Sham ovaries. In O in Y recipients the contralateral endogenous ovary is young; in Y in O recipients it is old. (B) Spearman correlations between unilateral transplantation signatures and reference signatures. The transplantation signatures are graft versus contralateral ovary within Y in O and O in Y recipients, O in Y graft versus O in O, and Y in O graft versus Y in Y. The references are ovarian aging (this study), liver, kidney, brain and rodent multi-tissue aging, rodent mortality, and longevity interventions (rapamycin, maximum lifespan, caloric restriction (CR) and growth hormone (GH) deficiency) (Tyshkovskiy *et al*, 2026; Tyshkovskiy *et al*, 2019). Correlations use log fold changes across the union of top-ranked genes in each pair of signatures (Methods). (C) GSEA (Hallmark, KEGG and Reactome) of the signatures in B. For transplantation contrasts, genes were ranked by the DESeq2 Wald statistic. Color shows NES. (D) Epigenetic age (eAge) from the universal pan-mammalian clock 2 (Universal2) across the unilateral transplantation groups in A. Kruskal-Wallis P = 9.3 × 10^−5^. (E) OvaAge (Bayesian ridge model) after bilateral transplantation, in which both ovaries were replaced. P = 0.0017 for Y in O versus Y in Y, P = 2.6 × 10^−5^ for O in Y versus O in O, and P = 1.2 × 10^−9^ for Young Sham versus Old Sham. (F) Spearman correlations between bilateral transplantation signatures from ovary, brain and uterus and the reference signatures listed in B. Pink labels give the reference group for each contrast (C: Y in Y or C: O in O). Group names give ovary (donor) age before recipient age: Y in O, young ovary in an old recipient; O in Y, old ovary in a young recipient; Y in Y and O in O, isochronic grafts (Y Iso, O Iso). Box plots show the median and interquartile range; points are individual samples. Pairwise P values are shown above brackets. In B, C and F, asterisks show BH-adjusted significance (*P < 0.05, **P < 0.01, ***P < 0.001).

In the unilateral design, principal component analysis (PCA) of ovarian transcriptomes separated young and old ovaries along PC1 (24.7% of variance; **Ext. Data Fig. 5A**). Aged ovaries transplanted into young hosts shifted toward young ovaries, whereas young ovaries transplanted into aged hosts shifted toward old ovaries. In both host ages, the contralateral endogenous ovary remained with untransplanted ovaries of its own age, so two ovaries in the same animal occupied different positions. In the methylome, samples separated mainly by host age (**Ext. Data Fig. 5B**). Within this structure, aged grafts in young hosts overlapped with young controls, whereas young grafts in aged hosts moved only partly toward the aged cluster.

OvaAge quantified these shifts (**Fig. 2A**). Aged ovaries transplanted into young hosts had lower tAge than old controls, and young ovaries transplanted into aged hosts had higher tAge than young isochronic grafts. In both directions the grafts moved toward the age of the host without reaching it, so that the transplanted and resident ovaries of the same animal occupied distinct molecular age states. Because both ovaries share the same circulation, host age shifted ovarian molecular age partially rather than resetting it.

Transcriptome-wide comparisons with aging and longevity signatures showed shifts in both directions (Fig. 2B). Relative to contralateral ovaries in the same host, aged grafts in young hosts correlated negatively with tissue aging and mortality signatures and positively with signatures of lifespan-extending interventions, including caloric restriction, growth hormone deficiency and rapamycin. Young grafts in aged hosts correlated only modestly and positively with aging signatures. At the pathway level, aged grafts in young hosts showed lower interferon-α/γ, IL6-JAK-STAT3, TNFα-NF-κB, complement, inflammasome, p53, apoptosis and ECM remodeling signatures and higher oxidative phosphorylation, mitochondrial translation, fatty acid metabolism, ribosome and mTORC1 signatures than their contralateral ovaries (**Fig. 2C**). Young grafts in aged hosts showed changes in the opposite direction.

These pathway shifts were not explained by changes in estimated follicular composition. Estimated epithelial and phagocyte fractions differed among transplantation groups (Kruskal-Wallis P = 0.0073 and 0.0070; Ext. Data Fig. 6A,B), whereas oocyte and granulosa cell fractions did not (Kruskal-Wallis P ≥ 0.17; Ext. Data Fig. 6C). Epithelial and phagocyte fractions were highest in old ovaries that remained in old hosts (old isochronic, old sham and the contralateral ovaries of young-into-old recipients), whereas old ovaries grafted into young hosts and young ovaries grafted into old hosts had fractions close to young controls. No single pairwise comparison remained significant after correction (two-sided Dunn test, BH-adjusted P ≥ 0.055). Other estimated cell types are shown in Ext. Data Fig. 6D. Epigenetic age also shifted (Fig. 2D). With the universal pan-mammalian DNA methylation clock 2 (Universal2), aged ovaries transplanted into young hosts had lower epigenetic age than aged controls, and young ovaries transplanted into aged hosts had epigenetic age intermediate between young and old ovaries. This was the only measure in which the two directions appeared to differ in magnitude, and the difference was partial.

Transplanted ovaries did not form a separate cluster but moved along the same axis that separates young and old ovaries. Because transplanted and contralateral ovaries were compared within the same animal, these differences cannot be attributed to endocrine variation between individuals. In the unilateral design, ovarian aging was therefore bidirectionally plastic. A young host shifted aged ovaries partially toward a young state, and an aged host shifted young ovaries partially toward an old state, across transcriptomic, pathway and epigenetic measures.

The contralateral ovary in the unilateral design could contribute to these responses through local or endocrine interactions. We therefore also performed bilateral heterochronic transplantation, replacing both ovaries (**Fig. 2E**). PCA of bilateral transcriptomes recapitulated the age axis observed in unilateral grafts, with heterochronic grafts in intermediate positions (**Ext. Data Fig. 5C**). In this design, both aged ovaries in young hosts and young ovaries in aged hosts underwent significant shifts in transcriptomic age (*P* = 2.6 × 10^−5^ and *P* = 0.0017, respectively; **Fig. 2E**), and both groups occupied intermediate states, consistent with partial shifts in both directions. Pathway enrichment (**Ext. Data Fig. 5D**) and correlation with external aging and intervention signatures (Tyshkovskiy *et al*, 2019; Tyshkovskiy *et al*, 2026) (**Fig. 2F**) showed shifts in the same inflammatory, mitochondrial and ECM programs identified in unilateral grafts. Multi-tissue transcriptomic clocks, applied without retraining, separated young and old ovarian controls weakly or not at all in both designs (**Ext. Data Fig. 7**), indicating that they are less sensitive to ovary-specific aging changes than OvaAge.

We next asked which metabolic programs accompanied these shifts, using gene set enrichment against Recon3D, KEGG, MitoCarta 3.0 and scFEA metabolic module collections (Alghamdi *et al*, 2021; Brunk *et al*, 2018; Kanehisa *et al*, 2021; Rath *et al*, 2021). In the ovarian aging reference signature, TCA cycle, oxidative phosphorylation, pyruvate metabolism, electron transport chain assembly, Fe-S cluster biogenesis and mitochondrial translation gene sets declined with age (**Ext. Data Figs 8** and **9**), consistent with impaired mitochondrial homeostasis in aging tissues (Ben-Meir *et al*, 2015; Bratic & Larsson, 2013; Lopez-Otin *et al*, 2023; May-Panloup *et al*, 2016; Smits *et al*, 2023; Tyshkovskiy *et al*, 2024). Young ovaries in aged hosts showed lower expression of cholesterol and steroid biosynthesis, fatty acid metabolism and mitochondrial fuel utilization genes, following the direction of ovarian aging and implicating the mevalonate-cholesterol axis, which supports steroidogenesis and oocyte quality (Caron *et al*, 1997; Liu *et al*, 2025) (**Ext. Data Fig. 9**). Aged ovaries in young hosts showed the opposite pattern, with higher expression of oxidative phosphorylation, mitochondrial ribosome and translation, Fe-S cluster, glycolysis and pyruvate metabolism genes and, more weakly, TCA cycle genes, together with one-carbon, amino acid and nucleotide biosynthesis programs (Ext. Data Fig. 8). Mitochondrial cholesterol, bile acid and steroid synthesis genes trended upward (FDR = 0.12), in line with studies linking cholesterol flux to oocyte and ovarian competence (Ben-Meir *et al*, 2015; Dunning *et al*, 2014; Liu *et al*, 2025). Together, these data indicate that ovarian aging involves a decline in mitochondrial and steroidogenic programs, and that the mitochondrial decline is at least partly reversible when an aged ovary is placed in a young host.

### Ovarian age propagates into recipient organs

Loss of ovarian function has consequences beyond fertility. In women, oophorectomy before menopause is associated with later cognitive impairment, while early bilateral oophorectomy is associated with increased cardiovascular mortality (Rivera *et al*, 2009; Rocca *et al*, 2007). In postreproductive mice, transplantation of young ovaries can extend lifespan and reduce cardiac pathology (Mason *et al*, 2009; Mason *et al*, 2011). These observations raise the question of how ovarian age influences molecular aging in other organs. We examined this relationship using reciprocal ovary transplantation in mice and a complementary analysis of ovariectomy and estradiol replacement in primates.

We first reanalyzed cortical transcriptomes of aged female rhesus macaques, a species that undergoes menopause (Walker & Herndon, 2008), comparing animals with intact ovaries, ovariectomized animals and ovariectomized animals treated with estradiol (Cervera-Juanes *et al*, 2024). In the occipital cortex, ovariectomy altered global gene expression (PERMANOVA *P* = 0.0049; pairwise *P* = 0.019 versus intact controls) and increased transcriptomic age (*P* = 0.004; **Fig. 3A,B**). Estradiol-treated animals did not differ detectably from untreated ovariectomized animals in global gene expression, and their transcriptomic age remained elevated relative to intact controls (*P* = 0.002), with module differences mainly in VEGF signaling and chromatin modification (**Fig. 3C**). In the prefrontal cortex, neither global gene expression nor transcriptomic age differed significantly among groups (**Fig. 3B and Ext. Data Fig. 10**), suggesting regional differences in the response to ovarian loss.

**Figure 3.**
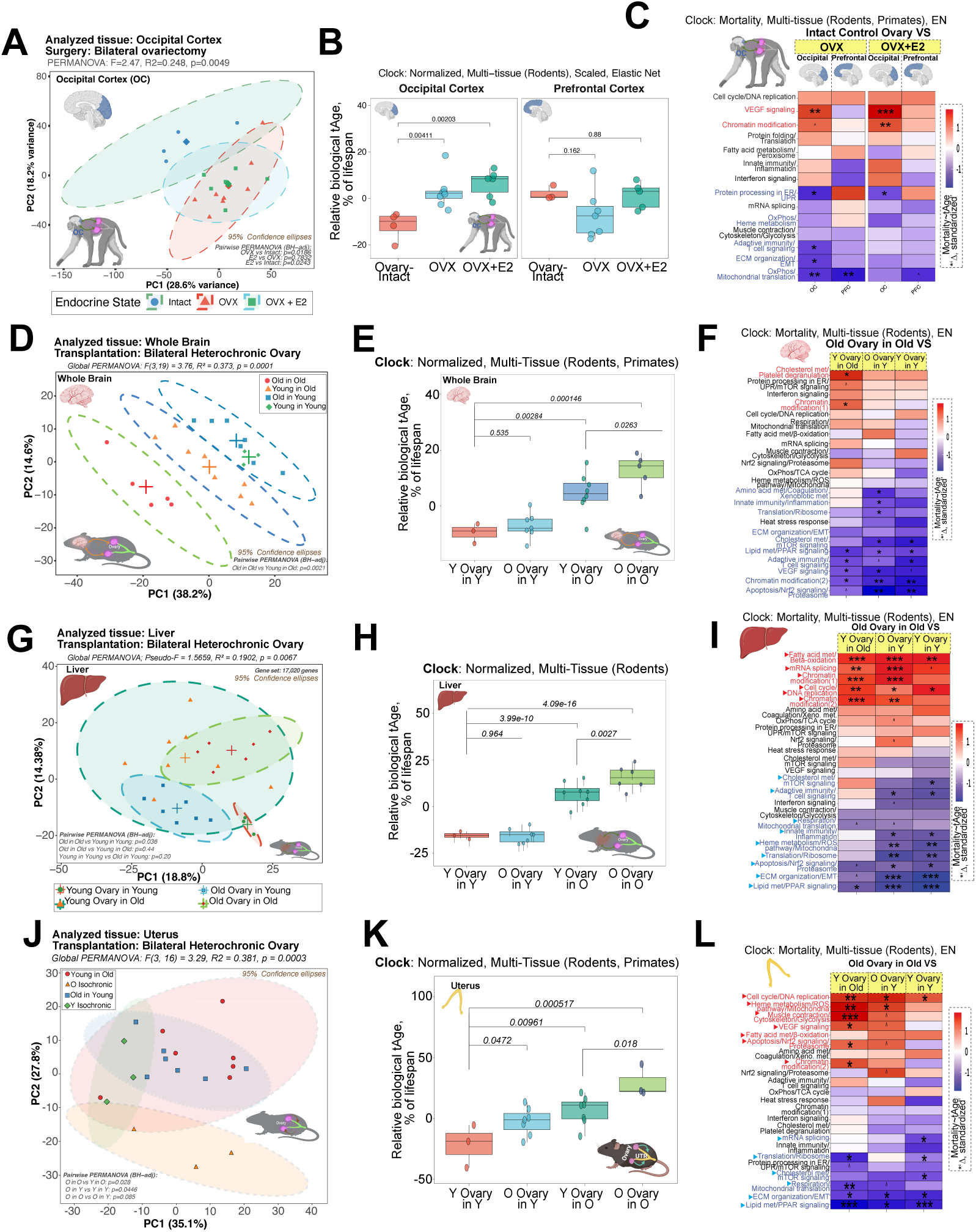
Ovarian status alters transcriptomic age in brain, liver and uterus. (A) Principal component analysis (PCA) of occipital cortex expression in old rhesus macaques that were ovary-intact, ovariectomized (OVX) or ovariectomized and treated with estradiol (OVX+E2). Data were reanalyzed from PRJNA1088413 (Cervera-Juanes *et al*, 2024). PERMANOVA F = 2.47, R^2^ = 0.248, P = 0.0049; BH-adjusted pairwise PERMANOVA P values are shown on the plot. (B) Relative biological tAge (% of lifespan) in occipital and prefrontal cortex from the scaled, normalized rodent multi-tissue elastic net clock (Tyshkovskiy *et al*, 2026). (C) Standardized differences in module-specific mortality tAge (multi-tissue elastic net clock trained on rodents and primates) for OVX and OVX+E2 versus intact animals, in each cortical region. (D, G, J) PCA of brain (D), liver (G) and uterine (J) expression after bilateral ovary transplantation in mice. Ellipses show 95% confidence regions. Global and BH-adjusted pairwise PERMANOVA statistics are shown on each plot. (E, H, K) Relative biological tAge (% of lifespan) in brain (E), liver (H) and uterus (K). Brain and uterus estimates come from the normalized multi-tissue clock trained on rodents and primates; liver estimates come from the normalized rodent multi-tissue clock (Tyshkovskiy *et al*, 2026). Y in O versus O in O gives P = 0.0263 (brain), 0.0027 (liver) and 0.018 (uterus). O in Y versus Y in Y gives P = 0.535, 0.964 and 0.0472. (F, I, L) Standardized differences in module-specific mortality tAge in brain (F), liver (I) and uterus (L) for Y in O, O in Y and Y in Y, each relative to O in O. In D-L, group names give ovary (donor) age before recipient age: Y in O, young ovary in an old recipient; O in Y, old ovary in a young recipient; Y in Y and O in O, isochronic grafts (Y Iso, O Iso). Box plots show the median and interquartile range; points are individual samples. In heatmaps, *P < 0.05, **P < 0.01, ***P < 0.001.

To test whether ovarian age itself influences recipient tissues, we compared mice receiving young or old ovaries within each recipient age group after bilateral transplantation. In old recipients, young ovaries altered brain gene expression (pairwise PERMANOVA *P* = 0.0021; **Fig. 3D**) and lowered normalized transcriptomic age relative to old ovaries (*P* = 0.026; **Fig. 3E**). In young recipients, old ovaries did not significantly increase brain transcriptomic age. The lower brain age in old recipients was accompanied by reduced cholesterol homeostasis signatures and inferred Srebf1 and Srebf2 activity, and by differences in the cholesterol metabolism, lipid and PPAR signaling, adaptive immunity and VEGF signaling modules of the mortality clock (**Fig. 3F** and **Ext. Data Fig. 11**), without a reduction in interferon signatures.

In the pituitary, where estradiol and inhibin regulate gonadotrope function (Gregg *et al*, 1991; Weiss *et al*, 1993), donor age did not significantly change transcriptomic age in either recipient age group (**Ext. Data Fig. 12A).** In old recipients, however, young ovaries reduced interferon and IL6-JAK-STAT3 signatures and increased estrogen response signatures, opposing the direction of pituitary aging, and in young recipients old ovaries increased TGFβ signaling and epithelial-mesenchymal transition signatures (**Ext. Data Fig. 12B-G).**

In the liver, where estrogen signaling contributes to the control of lipid and cholesterol content (Palmisano *et al*, 2017; Pedram *et al*, 2013), global gene expression did not differ significantly between donor ages within old recipients (**Fig. 3G**), but young ovaries lowered normalized transcriptomic age (*P* = 0.0027; **Fig. 3H**) and mortality-clock age (*P = 1.5 × 10*^−5^; **Ext. Data Fig. 13A**). Young ovaries also reduced cholesterol homeostasis signatures and inferred Srebf1 and Srebf2 activity (**Ext. Data Fig. 13B-F**), and mortality-clock modules for fatty acid metabolism and β-oxidation, mRNA splicing, chromatin modification and the cell cycle differed between donor ages (**Fig. 3I**). Old ovaries did not change hepatic transcriptomic age in young recipients but reduced oxidative phosphorylation, fatty acid, bile acid and xenobiotic metabolism signatures, resembling the reference pattern of liver aging (**Ext. Data Fig. 13D,E**).

Ovarian steroids drive cyclic uterine growth and prepare the endometrium for pregnancy, while reduced responsiveness to these hormones contributes to age-related defects in decidualization and placentation (Woods *et al*, 2017). The uterus was the only organ in which donor age altered transcriptomic age in both young and old recipients. Uterine gene expression differed between donor ages within old (*P = 0.028*) and young recipients (*P* = 0.045; **Fig. 3J**). Young ovaries lowered uterine transcriptomic age in old recipients (*P = 0.018*), while old ovaries increased it in young recipients (*P* = 0.047; **Fig. 3K**). In old recipients, young ovaries increased estrogen response, E2F target, G2M checkpoint and mTORC1 signatures and inferred estrogen, EGFR, MAPK and PI3K activity, consistent with greater hormonal stimulation of the uterus. In young recipients, old ovaries increased interferon response signatures and inferred JAK-STAT activity (**Fig. 3L** and **Ext. Data Fig. 14**).

Uterine gene expression changes induced by young ovaries in old recipients correlated negatively with aging metasignatures from several organs, whereas those induced by old ovaries in young recipients correlated positively (**Fig. 2F**). Across the organs examined, young ovaries lowered transcriptomic age in the old brain, liver and uterus. The reciprocal increase following transplantation of old ovaries was detected only in the young uterus, and pituitary age estimates did not change significantly in either recipient age group. Ovarian age can therefore influence molecular aging beyond the reproductive tract, with the extent and direction of the response depending on the recipient tissue.

### Transplantation and parabiosis reveal shared ovarian responses to an older environment

Heterochronic transplantation exposes an ovary to a host of a different age while also altering its vascular and neural connections. Heterochronic parabiosis provides a complementary test of systemic influences by leaving the ovary in place while the animal shares its circulation with a partner of a different age. Previous studies have shown that exposure to a young circulation improves aged progenitor cell function and lowers biological age in blood and liver (Conboy *et al*, 2005; Zhang *et al*, 2023). We joined mice aged 3.5 and 7.5 months for three months, separated them and collected ovaries one month later, at 7.5 and 11.5 months of age. Isochronic pairs served as controls ( **Fig. 4A** and **Ext. Data Fig. 15A**). The older cohort was selected near the onset of reproductive decline, giving a four-month age difference between partners.

**Figure 4.**
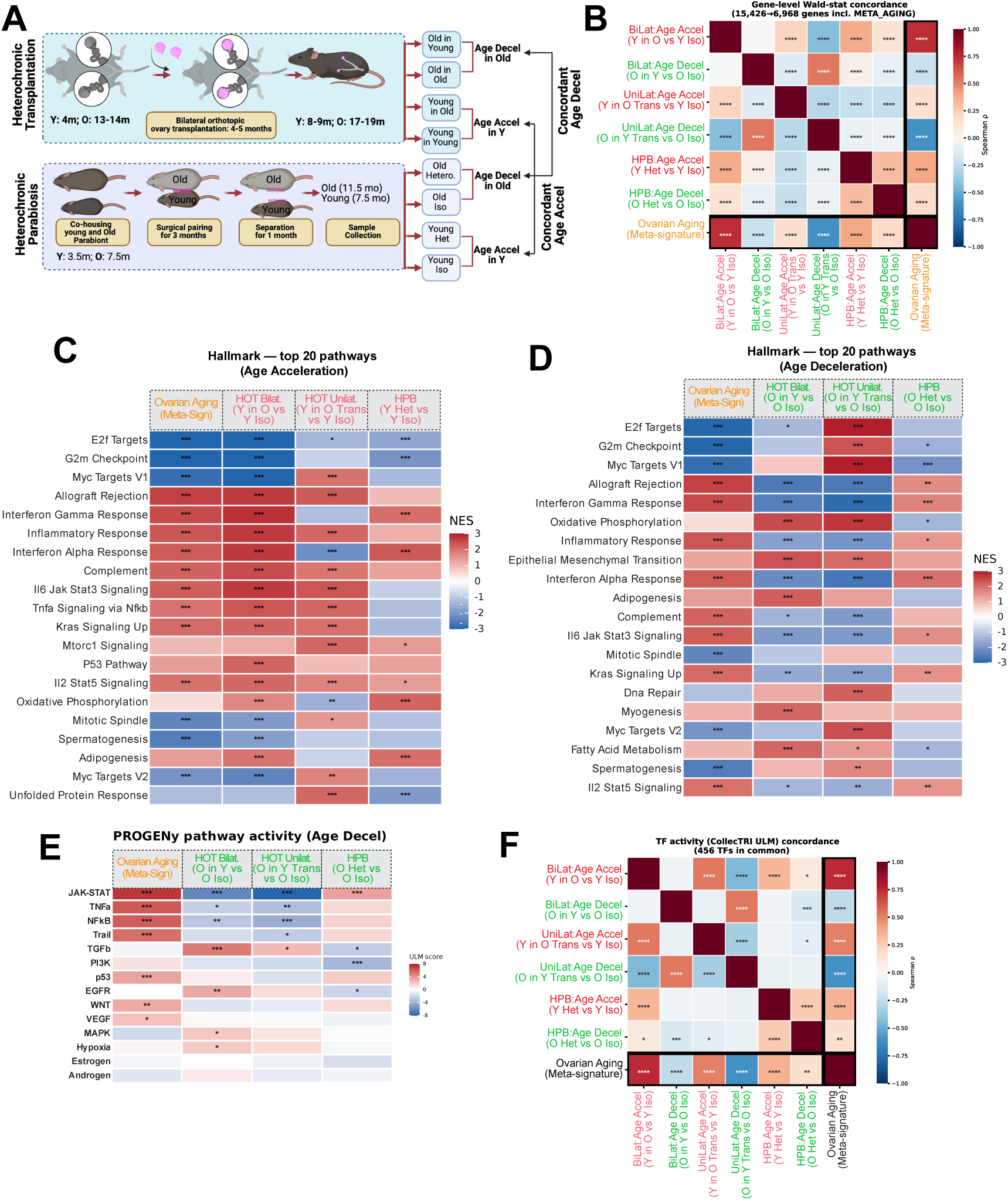
Ovarian expression responses shared by transplantation and parabiosis. (A) Experimental designs. Top, bilateral orthotopic ovary transplantation between young (4 months) and old (13-14 months) mice, analyzed 4-5 months later (8-9 and 17-19 months). Bottom, heterochronic parabiosis. Mice aged 3.5 and 7.5 months were co-housed, surgically joined for 3 months and separated for 1 month before collection (7.5 and 11.5 months). Contrasts with young ovaries in an old environment are labeled “age acceleration”; contrasts with old ovaries in a young environment are labeled “age deceleration”. (B) Spearman correlations of gene-level DESeq2 Wald statistics across six intervention contrasts and the ovarian aging reference signature. The intervention contrasts are bilateral transplantation (BiLat), unilateral transplantation (UniLat) and parabiosis (HPB), each in both directions. 6,968 genes are shared by all signatures. (C, D) Hallmark GSEA for the age-acceleration contrasts (C: Y in O vs Y Iso; Y in O Trans vs Y Iso; Y Het vs Y Iso) and the age-deceleration contrasts (D: O in Y vs O Iso; O in Y Trans vs O Iso; O Het vs O Iso), shown alongside the ovarian aging reference signature. Rows are the 20 top-ranked pathways. Color shows NES. (E) PROGENy signaling activity (univariate linear model (ULM) score) for the contrasts in D (Schubert *et al*, 2018; Badia-i-Mompel *et al*, 2022). (F) Spearman correlations of CollecTRI-based ULM transcription-factor activity scores across the contrasts in B, using 456 shared transcription factors (Müller-Dott *et al*, 2023). Het, heterochronic partner; Iso, isochronic control; Trans, graft. Unilateral controls pool isochronic and sham ovaries. The reference column is the ovarian lifespan signature of this study. Asterisks show BH-adjusted significance: *P < 0.05, **P < 0.01, ***P < 0.001, ****P < 0.0001. The labels “age acceleration” and “age deceleration” name the direction of exposure.

In younger heterochronic partners, OvaAge estimates were higher than in young isochronic controls with both OvaAge models (elastic net and Bayesian ridge) (*P = 0.0026 and 3.6 × 10*^−5^; **Ext. Data Fig. 15B,C**), as was the estimate of the innate immunity and inflammation module-specific clock (*P* = 0.002; **Ext. Data Fig. 15D**). Age estimates for older heterochronic partners were lower than those of older isochronic controls, but not significantly (*P = 0.12 and 0.13*). Exposure to an older circulation therefore induced an older ovarian expression profile in younger partners that persisted for at least one month after separation.

To identify processes shared across interventions, we compared ovarian differential expression across 6,968 genes common to all contrasts. Young ovaries exposed to an older environment, whether by bilateral or unilateral transplantation or by parabiosis, showed expression changes that correlated positively with the ovarian aging signature, most strongly after bilateral transplantation, whereas old grafts in young hosts correlated negatively in both transplantation designs (**Fig. 4B**). Inferred activities of 456 transcription factors showed the same grouping (**Fig. 4F**). Immune and inflammatory signatures increased in young ovaries exposed to older environments in all three designs, and E2F target and G2M checkpoint signatures decreased in bilateral grafts and parabionts (**Fig. 4C and Ext. Data Fig. 16A,C**). Old grafts in young hosts showed the reciprocal response in both transplantation designs, with lower interferon, inflammatory and IL6-JAK-STAT3 signatures, lower inferred JAK-STAT, TNFα and NF-κB activity, and higher oxidative phosphorylation, fatty acid metabolism, ribosome and translation signatures (**Fig. 4D,E and Ext. Data Fig. 16B,D**).

The ovaries of older heterochronic partners did not show the same response as old grafts placed in young hosts. Interferon and inflammatory signatures were not reduced and, in some cases, increased relative to isochronic controls. Their overall expression signature also failed to oppose the reference aging signature (**Fig. 4B,D**), consistent with the absence of a significant decrease in OvaAge. Thus, the molecular response to sharing circulation with a partner four months younger differed from that observed after transplantation into a young host. Since the experiments differed in animal age, surgery and timing of collection, this divergence cannot be attributed to the mode of systemic exposure alone.

Spi1 (PU.1), Irf1, Nfkb1 and Rela, among the transcription factors with the highest inferred activity during ovarian aging, were more active in young grafts in old hosts and less active in old grafts in young hosts (**Ext. Data Fig. 16E,F**). These regulators participate in myeloid differentiation, interferon signaling and NF-κB-dependent inflammation, consistent with the age-related changes in ovarian immune-cell composition (Ben Yaakov *et al*, 2023; Isola *et al*, 2024). An older environment thus increased inflammatory gene expression in young ovaries, whereas a younger environment reduced it in aged grafts and increased mitochondrial and biosynthetic programs, changes that resemble the expression signatures of lifespan-extending interventions such as caloric restriction and rapamycin (**Fig. 2B**).

### Ovarian DNA methylation responds to host age across multiple biological processes

Next, we examined how ovarian methylation changes across the lifespan and responds to transplantation. The first principal component of the 5,000 most variable CpGs explained 69% of the variance and ordered ovaries from development through peak reproductive activity to acyclicity (**Ext. Data Fig. 17A**). After unilateral transplantation, differentially methylated positions were detected almost exclusively in old grafts placed in young recipients. Few were detected in young ovaries in old recipients and none in contralateral endogenous ovaries (**Ext. Data Fig. 17B-D**). Among positions that lost methylation in old ovaries, a higher proportion mapped to CpG islands and shores and to active promoter and enhancer annotations than in the array background. The clearest changes at individual CpGs therefore occurred in aged ovaries exposed to a young host and included regions implicated in transcriptional regulation.

To link these methylation patterns to biological function, we applied pre-trained pathway-specific methylation clocks (PathAge) from the Ageome framework (Ying *et al*, 2024). Each PathAge model is an elastic net model based on CpGs assigned to the regulatory domains (McLean *et al*, 2010) of genes in a given pathway. We applied 1,808 PathAge models and their weighted composite, MetroAge, which correlated with chronological age across the ovarian series (r = 0.77, **Fig. 5A**). These models estimate age from methylation at CpGs assigned to pathway genes, without directly measuring pathway activity. MetroAge was lower in old ovaries in young recipients than in old isochronic ovaries ( *p = 0.028*) and higher in young ovaries in old recipients than in young isochronic ovaries (*p = 0.015*, **Fig. 5B**). Old endogenous ovaries sharing a host with a young ovary did not differ significantly from old isochronic ovaries. Thus, the combined methylation estimate detected shifts in both directions, even though individually significant CpG changes were concentrated in old ovaries exposed to a young host.

**Figure 5.**
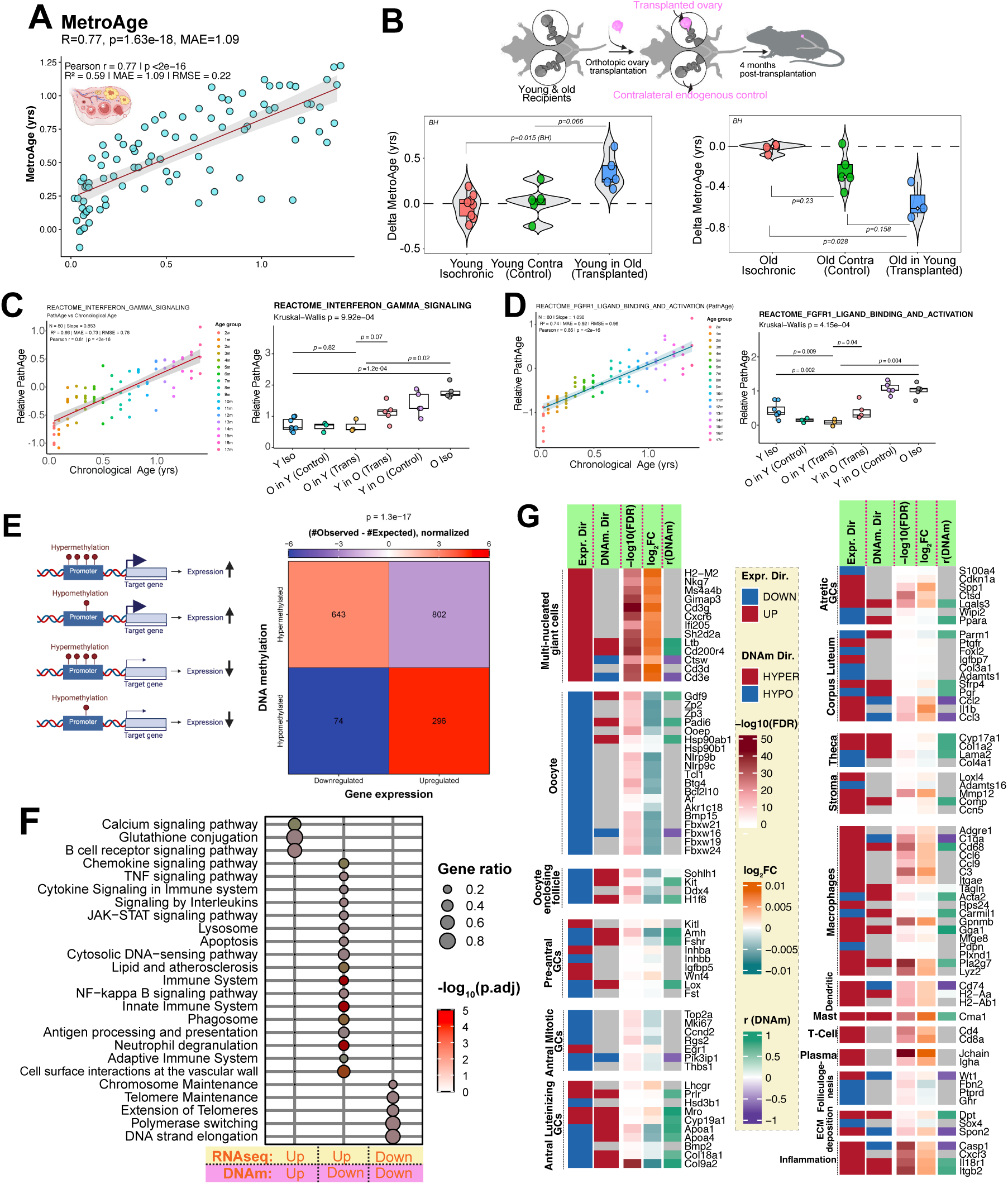
Pathway-specific methylation clocks and coupling of ovarian methylation and expression. (A) MetroAge against chronological age. MetroAge is a weighted average of all PathAge clock predictions, with each clock weighted by the inverse of its squared cross-validated root mean squared error (RMSE) (Ageome framework; Ying *et al*, 2024). Pearson r = 0.77, R^2^ = 0.59. (B) ΔMetroAge after unilateral transplantation, relative to the corresponding isochronic group (dashed line at 0). Left, young ovaries: Y Iso, young contralateral ovaries from O in Y recipients, and Y in O grafts. Y in O versus Y Iso gives BH-adjusted P = 0.015. Right, old ovaries: O Iso, old contralateral ovaries from Y in O recipients, and O in Y grafts. O in Y versus O Iso gives P = 0.028. (C, D) PathAge clocks for REACTOME-INTERFERON-GAMMA-SIGNALING (C) and REACTOME-FGFR1-LIGAND-BINDING-AND-ACTIVATION (D). Each pre-trained PathAge model (Ying et al, 2024) is an elastic net model of CpGs within the pathway genes’ GREAT regulatory domains (McLean *et al*, 2010). Left, the lifespan series (n = 80), colored by age; C: r = 0.81, D: r = 0.86. Right, unilateral transplantation groups. Kruskal-Wallis P = 9.9 × 10^−4^ (C) and 4.2 × 10^−4^ (D). Pairwise comparisons use Welch’s t-tests with BH correction. (E) Coupling of age-associated promoter methylation with age-associated gene expression in 90 ovarian samples. The schematic defines the four directional classes. The matrix gives gene counts (hypermethylated: 643 down, 802 up; hypomethylated: 74 down, 296 up); color shows (observed − expected)/expected. Overall P = 1.3 × 10^−17^. (F) Pathway over-representation in the three methylation-expression classes indicated below the plot (expression/methylation: up/up, up/down and down/down). Dot size shows the gene ratio; color shows −log10 adjusted P. (G) Selected coupled genes grouped by ovarian cell type or state using marker definitions from Lan et al (2026). Columns show expression direction, methylation direction, expression −log10 (FDR), log2 fold change and the correlation of promoter methylation with age (r(DNAm)). Cell-type labels annotate genes measured in bulk tissue. Group names give ovary (donor) age before recipient age: Y in O, young ovary in an old recipient; O in Y, old ovary in a young recipient; Y in Y and O in O, isochronic grafts (Y Iso, O Iso). Box plots show the median and interquartile range; points are individual samples.

The pathway models connected these responses to processes involved in follicle growth, metabolism, stromal organization and immune function. Models for fibroblast growth factor receptor signaling and its SHC and PI3K cascades closely tracked chronological age (r = 0.86 to 0.90), identifying age-associated methylation in genes involved in communication among follicular somatic cells (Buratini & Price, 2011). The PI3K-AKT-mTOR model also tracked age (r = 0.74), implicating this primordial follicle activation pathway in these age-related changes (Adhikari *et al*, 2010; Castrillon *et al*, 2003; Reddy *et al*, 2008). Other models covered fatty acid metabolism and mitochondrial β-oxidation (r = 0.82), collagen crosslinking (r = 0.66), which is relevant to the increasing stiffness of aged ovarian stroma (Amargant *et al*, 2020), and heparan sulfate synthesis (r = 0.86). Models for interferon-γ, interferon-α, interleukin-18 and JAK-STAT signaling tracked age (r = 0.61 to 0.81), as did those for SNARE-mediated vesicle fusion and the G2/M phase of the cell cycle (r = 0.75 to 0.84, **Fig. 5C,D** and **Ext. Data Figs 18-20**). PathAge estimates also differed among the transplant groups for every model shown (Kruskal-Wallis P ≤ 0.02), and old grafts in young recipients had lower PathAge estimates than old isochronic ovaries. For interferon-γ signaling, old isochronic ovaries had higher estimates than young isochronic ovaries (*p = 1.2 × 10^-^⁴*), while old grafts in young recipients were not statistically distinguishable from young isochronic ovaries. Young grafts in old recipients also remained below old isochronic ovaries (*p = 0.02*, **Fig. 5C**). The FGFR1 ligand-binding model likewise yielded lower estimates for old grafts in young recipients than for old isochronic ovaries ( *p = 0.004*, **Fig. 5D**). Together, the response in aged ovaries thus included methylation patterns associated with follicular signaling, metabolism, stromal structure and immunity.

We next related age-associated methylation changes to gene expression, following an approach used to identify aging genes in human blood (Moqri *et al*, 2026). Across 90 ovaries with both measurements, the direction of promoter methylation change was associated with the direction of expression change (*p = 1.3 × 10⁻¹⁷*, **Fig. 5E**). Promoter hypermethylation with reduced expression (643 genes) and hypomethylation with increased expression (296 genes) occurred more often than expected, consistent with an inverse association between promoter methylation and transcription. A further 802 genes increased in expression despite promoter hypermethylation, although this combination was less frequent than expected. These groups differed in biological function (**Fig. 5F**). Genes that gained both promoter methylation and expression were enriched for calcium signaling, glutathione conjugation and B cell receptor signaling. Genes that lost methylation and gained expression were enriched for cytokine, chemokine, TNF, NF-κB and JAK-STAT signaling, together with lysosome, phagosome and antigen presentation pathways. Genes that lost both methylation and expression were enriched for chromosome maintenance, telomere maintenance and DNA replication. The correspondence between immune gene expression and promoter hypomethylation links the two molecular measurements to the increasing immune component of the aging ovary. Because both measurements were made in whole ovaries, these associations may reflect changes in cell abundance as well as regulation within individual cell types.

We then used ovarian cell-type markers (Lan *et al*, 2026) to link these molecular changes to specific tissue compartments (**Fig. 5G and Ext. Data Fig. 21**). Expression of oocyte-associated genes, including *Gdf9*, *Zp2*, *Zp3*, *Padi6*, *Bmp15* and *Nlrp9b*, declined with age, consistent with follicle loss, and several gained promoter methylation. This group includes factors required for folliculogenesis and early embryonic development (Dong *et al*, 1996; Yurttas *et al*, 2008). Macrophage-associated genes, including *Adgre1*, *Cd68*, *C1qa* and *Lyz2*, increased in expression. Genes enriched in multinucleated giant cell regions (Converse *et al*, 2025), including *Ms4a4b*, *Gimap3*, *Cd3g*, *Ltb*, *Ctsw* and *Cd200r4*, showed a particularly consistent increase with age. These opposing oocyte and immune trajectories led us to examine multinucleated giant cell regions directly.

### Ovarian multi-nucleated giant cell regions have distinct molecular and metabolic profiles

Macrophage-derived multinucleated giant cells (MNGCs) accumulate in the aging ovarian stroma, where they are found alongside fibrosis (Briley *et al*, 2016). Their abundance increases with age in mice and nonhuman primates, and regions containing these cells express genes involved in immune responses and cellular degradation (Converse *et al*, 2025). We asked whether the transcriptional features of these regions change with host age and whether they differ in predicted age from other ovarian compartments. Genes enriched in MNGC regions decreased in expression in old grafts placed in young recipients and increased in young grafts placed in old recipients, whereas oocyte-associated genes changed in the opposite direction (**Fig. 6A**). Representative sections showed prominent pigmented giant cell aggregates in old ovaries and less conspicuous aggregates in old grafts recovered from young recipients (**Ext. Data Fig. 22A**), consistent with their lower estimated phagocyte fraction (**Ext. Data Fig. 6B**).

**Figure 6.**
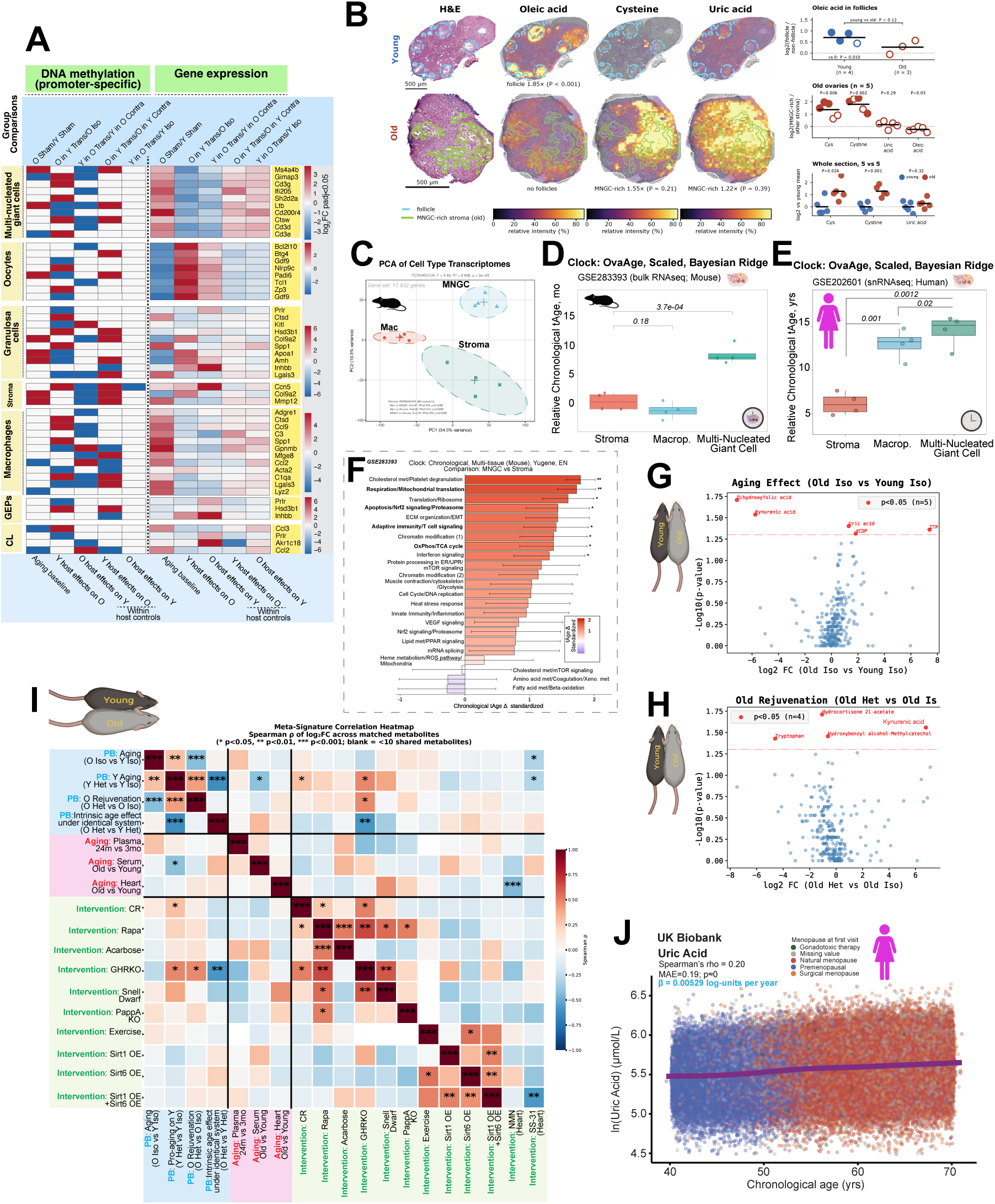
Molecular, transcriptomic and metabolomic features of multinucleated giant cell-rich ovarian regions. (A) Promoter methylation (left) and expression (right) log2 fold changes for selected marker genes of multinucleated giant cells (MNGCs), oocytes, granulosa cells, stroma, macrophages, germline epithelium progenitors (GEPs) and corpus luteum (CL). Only changes with adjusted P ≤ 0.05 are colored. Columns are the comparisons labeled above each column: the aging baseline (O Sham vs Y Sham), between-host comparisons of grafts with isochronic controls, and within-host comparisons of grafts with contralateral ovaries. (B) MALDI imaging of representative young and old ovaries. Registered adjacent H&E sections show follicles (cyan) and MNGC-rich stroma (green; old only). Oleic acid, cysteine and uric acid ion images ([M−H]−, 25 µm) are overlaid on the H&E on a shared relative scale; values below give per-ovary enrichment and spatial permutation P. Scale bars, 500 µm. Graphs show follicular oleic acid enrichment (young n = 4, old n = 3), MNGC-rich versus other stroma in old ovaries (n = 5), and whole-section intensities (n = 5 per group). Points are ovaries (filled, P < 0.05) and bars are means; P values are from t-tests. (C) PCA of mouse laser-capture profiles of stroma, macrophages (Mac) and MNGC regions (GSE283393; Converse *et al*, 2025; 17,932 genes). PERMANOVA statistics are shown on the plot. (D) OvaAge (scaled Bayesian ridge model) for the profiles in C; four captures per group. MNGC versus stroma P = 3.7 × 10−4 and macrophage versus stroma P = 0.18 (Welch’s t-tests on per-capture predictions). (E) OvaAge for human ovarian single-nucleus pseudobulk profiles (GSE202601; Jin *et al*, 2025): stroma, macrophages, and macrophage-associated nuclei selected by the mouse MNGC signature. (F) Standardized differences in module-specific chronological tAge (YuGene-normalized mouse multi-tissue elastic net clock) between mouse MNGC captures and stroma; error bars show s.e. (G, H) Volcano plots of plasma metabolites from parabiosis mice collected 1 month after separation: O Iso versus Y Iso (G) and O Het versus O Iso (H). Red points have nominal P < 0.05 (Mann-Whitney U test). (I) Spearman correlations of metabolite log2 fold changes between the four parabiosis (PB) comparisons and external metabolomic signatures of aging and longevity interventions. Correlations use matched metabolites; blank cells have fewer than 10 shared metabolites. (J) Natural-log plasma uric acid against age in 254,095 women from UK Biobank, colored by menopausal status at first visit. The curve is a generalized additive model fit (restricted maximum likelihood) with 95% confidence band; Spearman ρ = 0.20. The age coefficient from a linear model adjusted for body mass index, Townsend deprivation index, IPAQ physical activity and assessment center is 0.00529 log-units per year. Group names give ovary (donor) age before recipient age: Y in O, young ovary in an old recipient; O in Y, old ovary in a young recipient; Y in Y and O in O, isochronic grafts (Y Iso, O Iso). Het, heterochronic partner. Box plots show the median and interquartile range; points are individual samples. *P < 0.05, **P < 0.01, ***P < 0.001.

We next applied the transcriptomic clocks to profiles of laser-captured MNGC-rich and stromal regions from aged mouse ovaries and of ovarian macrophages isolated from young mice (Converse *et al*, 2025). The three sample groups had distinct transcriptomes (PERMANOVA *P = 5 × 10*^−4^; **Fig. 6C**). OvaAge predictions were higher in MNGC-rich regions than in stroma (P = 3.7 × 10−4, Welch’s t-test), whereas macrophage predictions did not differ significantly from stroma (**Fig. 6D**). Multi-tissue chronological, mortality and normalized clocks also gave the highest estimates for MNGC-rich regions (**Ext. Data Fig. 22B-E**). Cholesterol metabolism, mitochondrial respiration and translation, ribosome and adaptive immunity modules contributed most to the difference from stroma in the chronological clock (**Fig. 6F**), and innate and adaptive immunity, interferon signaling and oxidative phosphorylation modules in the mortality clock (**Ext. Data Fig. 22F-I**). MNGC-rich regions were enriched for interferon response, lysosome and antigen presentation gene sets relative to both macrophages and stroma, and their inferred TNFα, NF-κB and JAK-STAT activity and senescence-associated expression were intermediate between those of macrophages and stroma (**Ext. Data Fig. 23**).

To investigate whether a related macrophage phenotype occurs in humans, we applied the mouse MNGC-region signature to single-nucleus data from human ovaries (Jin *et al*, 2025). The signature identified a subset of ovarian macrophages (**Ext. Data Fig. 24**), designated MNGC-like on the basis of their expression profile. These cells had higher predicted transcriptomic age than other macrophages (*p = 0.02*) and stromal cells (*p = 0.0012*) from the same donors (**Fig. 6E**). The mouse signature therefore identifies a human macrophage population with stronger age-associated expression.

Relative to macrophages, MNGC-rich regions were enriched for gene sets involved in oxidative phosphorylation, respiratory complexes I and IV, mitochondrial ribosomes, glutathione metabolism and cytochrome P450 function, consistent with the original characterization of these regions (Converse *et al*, 2025), and genes of the glutathione-to-glutamate scFEA module showed the largest increase. Relative to stroma, MNGC-rich regions were enriched for lysosomal transport and glycan degradation gene sets and had lower expression of tricarboxylic acid cycle, branched-chain amino acid and fatty acid oxidation gene sets (**Ext. Data Fig. 25**). Lysosomal processing and altered mitochondrial and redox metabolism are therefore prominent features of the giant cell microenvironment.

To examine metabolism in its tissue context, we mapped metabolite signals across sections of five young and five old ovaries by matrix-assisted laser desorption/ionization (MALDI) mass spectrometry imaging, and registered each ion image to an adjacent H&E-stained section. In young ovaries, the signal assigned to oleic acid was concentrated in follicles (1.6-fold relative to non-follicular tissue; p = 0.01; **Fig. 6B**). Oleic acid can protect bovine oocytes from saturated fatty acid toxicity in culture (Aardema *et al*, 2011), providing a functional context for its follicular localization. Old ovaries contained few follicles (none in two of five sections), and where follicles were present, their oleic acid enrichment was smaller and not significant (1.2-fold, n = 3). In old ovaries, the signals assigned to cysteine and its oxidized dimer cystine were instead concentrated in MNGC-rich stroma, identified on H&E as pale-tan, vacuolated fields containing ceroid pigment (2.6-fold and 3.5-fold relative to other stroma; p = 0.006 and p = 0.002, respectively), in line with the glutathione-related expression of these regions described above. The uric acid signal was not consistently enriched in MNGC-rich stroma (1.1-fold, P = 0.29), and the oleic acid signal was lower there (0.9-fold). Across whole sections, cysteine and cystine signals were about 2.4-fold higher in old than in young ovaries (P = 0.026 and P = 0.001; **Fig. 6B**).

Although uric acid was not localized to MNGC-rich stroma, purine catabolism is linked to inflammation. Xanthine oxidoreductase produces uric acid, and its oxidase form also generates hydrogen peroxide (Kelley *et al*, 2010). Reactive oxygen species derived from this enzyme promote NLRP3-dependent interleukin-1β release in macrophages (Ives *et al*, 2015), urate crystals activate the NLRP3 inflammasome (Martinon *et al*, 2006), and NLRP3 inhibition delays ovarian aging in mice (Navarro-Pando *et al*, 2021). We therefore examined circulating metabolites in the parabiosis cohort one month after separation. Uric acid, dCDP and TTP were higher, while dihydroxyfolic acid and kynurenic acid were lower, in old compared with young isochronic mice (**Fig. 6G**). In old heterochronic mice, kynurenic acid was higher and tryptophan lower than in old isochronic mice (**Fig. 6H**). Across all metabolites, the response of old mice to a young partner was inversely correlated with the effect of age (ρ = -0.42; **Ext. Data Fig. 26E**). The responses of old and young heterochronic mice also correlated positively with the metabolite signature of long-lived growth hormone receptor knockout mice (ρ = 0.47 and 0.48, respectively; **Fig. 6I and Ext. Data Fig. 26A,B**). Further comparisons with external signatures are shown in Ext. Data Fig. 26C,D,F,G. These exploratory associations suggest that heterochronic exposure alters aspects of systemic metabolism that also vary with age and longevity.

Circulating uric acid was also associated with age in women. In 254,095 UK Biobank participants, uric acid concentrations were higher at older ages between 40 and 70 years (β = 0.00529 ln units per year after adjustment for body mass index, deprivation index, physical activity and assessment center, **Fig. 6J**), consistent with the higher serum urate reported after natural and surgical menopause (Hak & Choi, 2008). Together, these circulating measurements suggest uric acid among the metabolic features associated with reproductive and systemic aging.

### Reciprocal ovarian transplantation supports reproductive function late in life

To assess the functional significance of the ovarian responses to host age, we first examined their relationship to oocyte aging. Mitochondrial dysfunction accompanies the age-related decline in oocyte quality (May-Panloup *et al*, 2016), and studies in old mice show that oocyte quality can be improved by NAD+ precursors, metformin during *in vitro* maturation and exposure to a young follicular environment (Bertoldo *et al*, 2020; Cao *et al*, 2022; Miao *et al*, 2020; Wang *et al*, 2024). We compared whole-ovary expression changes after bilateral transplantation with reference signatures derived from oocyte aging metasignatures. Old ovaries transplanted into young recipients showed expression changes opposite to the oocyte aging signatures, resembling the responses to nicotinamide mononucleotide treatment and transfer of old oocytes into young follicles (**Fig. 7A**). Mitochondrial gene sets (Rath *et al*, 2021) that declined in both oocyte aging signatures and whole-ovary aging, including oxidative phosphorylation, mitochondrial ribosome and translation, increased in old ovaries transplanted into young recipients. These gene sets were also more highly expressed in young than in old grafts within old recipients (**Fig. 7B**). The agreement implicates mitochondrial processes associated with oocyte competence in the ovarian response to host age.

**Figure 7.**
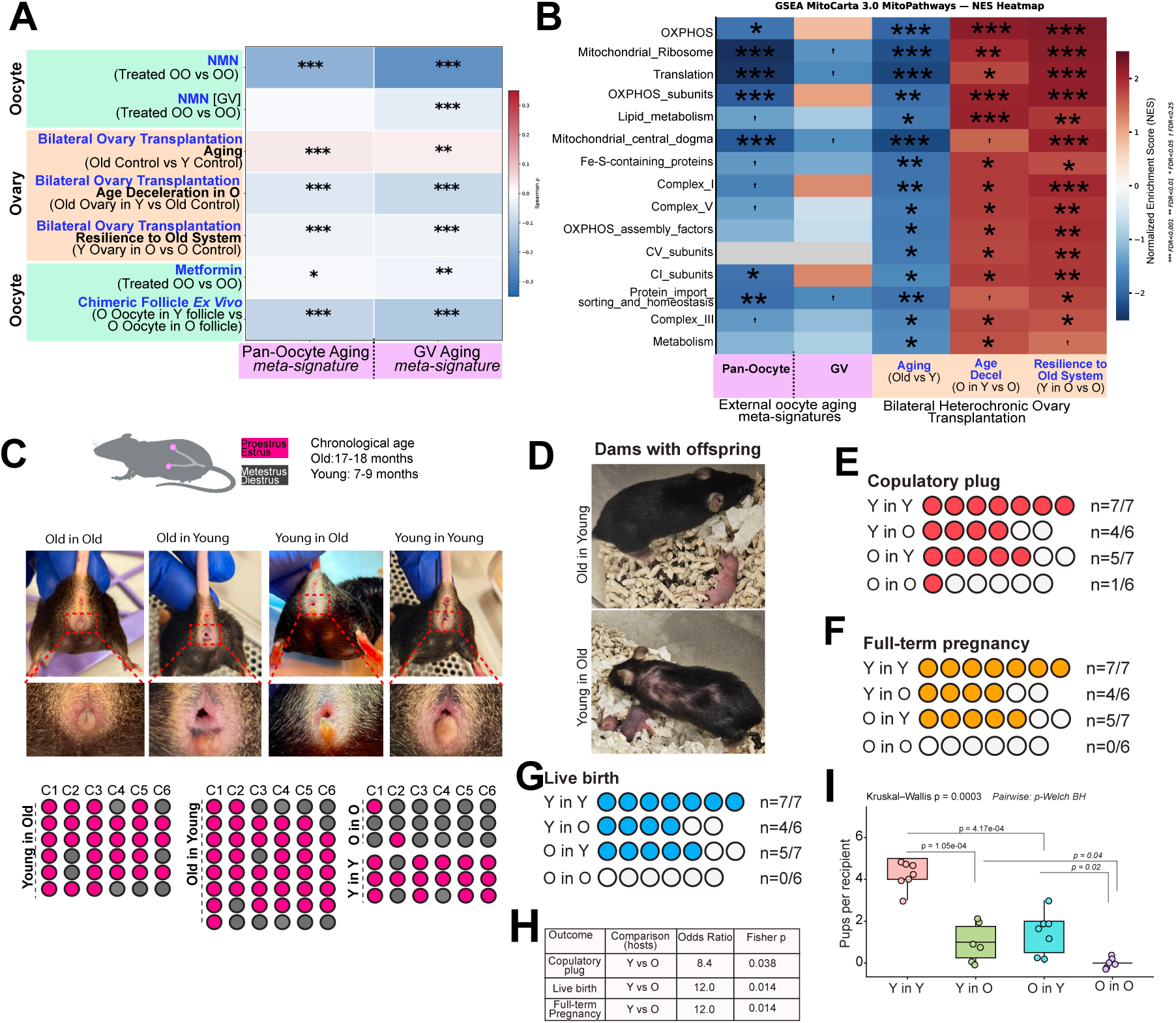
Reproductive outcomes after reciprocal ovary transplantation. (A) Spearman correlations between the pan-oocyte and germinal-vesicle (GV)-stage oocyte aging meta-signatures and expression responses to the indicated interventions. The interventions are nicotinamide mononucleotide (NMN; GSE153920, GSE183271; Miao *et al*, 2020), metformin (GSE201098; Cao *et al*, 2022), chimeric follicles reconstructed ex vivo (GSE270016; Wang *et al*, 2024) and bilateral ovary transplantation. The transplantation contrasts use bulk ovarian expression. Y in O versus O in O compares donor ages within old recipients. (B) GSEA of MitoCarta 3.0 MitoPathways (Rath *et al*, 2021) in the external oocyte aging meta-signatures and in the bilateral transplantation contrasts: aging (old vs young controls), O in Y vs O in O, and Y in O vs O in O. Color shows NES. (C) Representative external genital morphology 4-5 months after bilateral transplantation (young recipients 7-9 months and old recipients 17-18 months of age), and serial estrous-stage observations (columns C1-C6). Each row is one animal. Pink, proestrus or estrus; grey, metestrus or diestrus. (D) Representative O in Y and Y in O dams with offspring. (E-G) Copulatory plug (E), full-term pregnancy (F) and live birth (G) after mating following transplantation; all recipients had been through one round of mating before transplantation. Each circle is one recipient; filled circles indicate the outcome. Group sizes and counts: Y in Y 7/7, Y in O 4/6, O in Y 5/7 and O in O 1/6 for plugs; 7/7, 4/6, 5/7 and 0/6 for pregnancy and live birth. (H) Outcomes pooled by recipient age across both donor ages. Sample odds ratios (young vs old recipients, no continuity correction) are 8.4 for plugs and 12.0 for pregnancy and live birth. Two-sided Fisher’s exact P = 0.038 and 0.014. (I) Mean number of live pups per recipient across mating rounds; recipients without a litter are shown as zero. Kruskal-Wallis P = 0.0003. Pairwise comparisons use Welch’s t-tests with BH correction. Group names give ovary (donor) age before recipient age: Y in O, young ovary in an old recipient; O in Y, old ovary in a young recipient; Y in Y and O in O, isochronic grafts (Y Iso, O Iso). Box plots show the median and interquartile range; points are individual samples. In A and B, *P < 0.05, **P < 0.01, ***P < 0.001.

We next tested whether the molecular responses were accompanied by reproductive function (**Fig. 7C-I**). Groups are named by donor ovary age followed by recipient age. Bilateral transplantation was performed when recipients were 3-4 (young) or 13 (old) months old. All recipients had been through one round of mating before surgery; in old recipients, mating and pregnancy took place 1-2 months before transplantation. Four to five months later, when young recipients were 7-9 months and old recipients 17-18 months of age, we recorded estrous stages and mated the animals (**Fig. 7C**). Copulatory plugs were detected in 7 of 7 young-in-young, 5 of 7 old-in-young, 4 of 6 young-in-old and 1 of 6 old-in-old recipients (**Fig. 7E**). Because C57BL/6J females lose regular estrous cycles during the second year of life (Felicio *et al*, 1986) and age-related uterine defects can compromise reproduction independently of oocyte age (Woods *et al*, 2017), these outcomes reflect the combined capacity of graft and recipient.

Every plugged recipient except the single old-in-old animal subsequently carried a pregnancy to term and delivered live pups (**Fig. 7D,F**,G). Ovaries with a chronological age of 17-18 months at mating therefore supported live birth in young recipients, whereas ovaries of the same age did not do so in old recipients (5 of 7 versus 0 of 6; Fisher’s exact P = 0.021). Some oocytes in aged ovaries therefore retain developmental competence that can be expressed in a young host. Conversely, recipients aged 17-18 months delivered live pups after receiving young ovaries (4 of 6 versus 0 of 6 with old ovaries; Fisher’s exact P = 0.061). These births show that the neuroendocrine system and uterus of some aged females remained capable of supporting ovulation, implantation, gestation and delivery after ovarian replacement, consistent with the lower transcriptomic age and increased estrogen-responsive gene expression of the uterus in aged recipients of young ovaries (**Fig. 3K and Ext. Data Fig. 14**).

Reproductive output nevertheless remained below that of young controls. The number of live pups per recipient was lower in young-in-old and old-in-young recipients than in young-in-young controls (P = 1.1 × 10⁻⁴ and 4.2 × 10⁻⁴, respectively), but higher than in old-in-old recipients (P = 0.04 and 0.02, respectively, Kruskal-Wallis P = 0.0003, **Fig. 7I**). When donor ages were pooled, young recipients had higher odds of mating (odds ratio 8.4) and live birth (odds ratio 12.0) than old recipients (**Fig. 7H**). At the group level, both heterochronic combinations had intermediate reproductive output, broadly consistent with their intermediate ovarian transcriptomic ages (**Fig. 2E**). The reciprocal outcomes reveal reproductive capacity in both aged ovaries and aged recipients. A young host supported reproductive function in aged ovaries, while young ovaries supported pregnancy and live birth in aged females.

## Discussion

Reproductive aging combines exhaustion of a finite follicle reserve with deterioration of the somatic niche that supports the remaining follicles (Faddy *et al*, 1992; Wallace & Kelsey, 2010; Wang *et al*, 2024). Our findings show that latent reproductive capacity persists in both the aged ovary and the aged host. Aged ovaries supported live births in young recipients, and young ovaries supported pregnancy and live birth in aged recipients. Follicle depletion sets the limits of reproductive lifespan, but the function of the remaining tissue depends on the systemic environment in which it resides.

The capacity of aged ovaries to sustain reproduction agrees with classic transplantation studies (Parkening *et al*, 1985) and with evidence that a young somatic environment can restore the developmental competence of aged oocytes (Wang *et al*, 2024). Chronological age alone therefore does not abolish the competence of the surviving oocyte pool. In aged ovaries transplanted into young hosts, mitochondrial programs increased and canonical aging signatures reversed, paralleling metabolic interventions such as NAD+ repletion that improve oocyte quality (Bertoldo *et al*, 2020). Whole-ovary expression cannot isolate changes within oocytes, but the reproductive outcomes indicate that an improved systemic and follicular environment allowed pre-existing oocytes to mature and support term development.

Live births in 17-18-month-old recipients of young ovaries show that the aged neuroendocrine axis and uterus can still support ovulation, implantation and gestation. Embryo-transfer studies have identified impaired decidualization and placentation as major causes of age-related reproductive decline, defects that a young uterine environment can overcome (Woods *et al*, 2017). Our data raise a complementary possibility, that a young ovary partly reconditions the aged uterus. Young ovaries lowered uterine transcriptomic age in old recipients and increased estrogen-responsive programs, and these uterine changes opposed aging signatures from several organs. Prior parity may influence uterine capacity (Gosden, 1979), but old isochronic controls with a similar breeding history did not reproduce, arguing against parity alone explaining the rescue. Reproductive output nevertheless remained below young-control levels in both heterochronic groups, indicating incomplete functional restoration.

Host age shifted the expression and methylation profiles of grafted ovaries in both directions without changing their chronological age, and in both directions the grafts remained molecularly distinct from the endogenous ovaries of their hosts. The aging ovarian state therefore reflects both the tissue’s own history and its current systemic environment. The incomplete convergence of grafts toward the age of the host suggests constrained plasticity, in which some molecular programs remain responsive to systemic cues while others are fixed by prior decline.

OvaAge integrates age-associated changes in follicular, stromal and immune populations into a single whole-organ estimate, complementing clocks developed for individual ovarian cell populations (Jing *et al*, 2025; Kim *et al*, 2025). To our knowledge, this is the first application of a whole-organ transcriptomic clock to ovarian transplantation and heterochronic parabiosis. OvaAge assigned younger ages to ovaries of long-lived Ames dwarf mice and to follicles in which aged oocytes were combined with young somatic cells (Schneider *et al*, 2017; Wang *et al*, 2024), and it discriminated age in granulosa cells and primate ovarian somatic cells (Isola *et al*, 2024; Wang *et al*, 2020). The clock response also tracked reproductive function, as old ovaries that became transcriptionally younger in young recipients supported live birth, whereas old ovaries in old recipients did not. Whether the magnitude of a clock response predicts the extent or duration of fertility recovery remains to be tested.

Pathway-specific methylation clocks resolved the epigenetic response into biological processes. The lower transcriptomic age of aged ovaries in young recipients was paralleled by younger PathAge estimates for follicular signaling, metabolism, extracellular matrix organization and immunity (Ying *et al*, 2024). Lower interferon and JAK-STAT expression in these grafts coincided with younger PathAge estimates for immune pathways, and higher expression of fatty acid metabolism genes coincided with younger estimates for mitochondrial β-oxidation, a process required for oocyte developmental competence (Dunning *et al*, 2010). Because PathAge models estimate age from methylation at CpGs assigned to pathway genes, they indicate where epigenetic age changed rather than measuring pathway activity.

These pathway responses also raise a question about the follicle reserve. PI3K signaling controls primordial follicle activation (Reddy *et al*, 2008), and collagen remodeling alters the structural constraints on growing follicles (Amargant *et al*, 2020). A younger methylation profile in these pathways cannot by itself distinguish accelerated follicle recruitment, which could deplete the reserve, from improved somatic support of existing follicles. Combining PathAge with gene expression, follicle counts and reproductive outcomes could separate these possibilities.

Paired methylation and expression profiles from the same ovaries, spanning development to reproductive senescence and extending to changes in host age, connected promoter methylation to transcription. Earlier studies combined the two layers in granulosa cells, porcine ovaries and individual mouse oocytes (Castillo-Fernandez *et al*, 2020; Xi *et al*, 2019; Yu *et al*, 2015a). Our design adds a developmental series and a test of responsiveness to host age, following the use of cross-assay agreement to identify aging genes in human blood (Moqri *et al*, 2026). Applied to pharmacological, dietary or genetic interventions, OvaAge, PathAge and paired methylation-expression analysis could distinguish interventions that produce similar overall age shifts but differ in their effects on follicular support, inflammation or tissue remodeling.

Our data point to inflammatory and stromal remodeling as plausible mediators between host age and the environment of residual follicles. Ovarian aging alters immune-cell abundance and polarization (Isola *et al*, 2024), and stromal collagen deposition increases tissue stiffness (Amargant *et al*, 2020). Inflammatory and matrix-associated expression in aged grafts responded to host age, suggesting that part of this environment remains modifiable. CD38 depletion preserves ovarian NAD+ and reproductive function (Yang *et al*, 2024), and inhibition of IL-11 signaling reduces matrix stiffness and improves fertility in aged rodents (Wu *et al*, 2026). The graft responses also shared features with expression signatures of caloric restriction, rapamycin and reduced growth hormone signaling (Tyshkovskiy *et al*, 2026), suggesting that ovarian plasticity engages pathways targeted by established longevity interventions.

MNGC-rich stroma had elevated transcriptomic age, and MNGC-associated genes responded to host age. Together with the characterization of these cells in aged ovarian stroma (Converse *et al*, 2025), these findings identify an inflammatory compartment that can coexist with follicles that retain developmental competence. A related signature in human ovarian macrophages (Jin *et al*, 2025) suggests that some features of this compartment are shared across species. Whether MNGC-rich regions mainly clear debris accumulated over repeated ovulation and repair, or create an environment detrimental to neighboring follicles, will require models that separate their degradative capacity from local inflammatory signaling.

Spatial metabolite mapping added a metabolic dimension to this compartment. Signals assigned to cysteine and cystine were enriched in MNGC-rich stroma of every old ovary examined, whereas oleic acid marked follicles in young ovaries. Together with the glutathione-related expression of MNGC-rich regions, this points to altered thiol and redox handling in the giant cell niche. Uric acid was not localized to these regions. Because urate crystals activate the NLRP3 inflammasome (Martinon *et al*, 2006) and modulating purine metabolism and NLRP3 pathways alters macrophage responses and ovarian aging (Ives *et al*, 2015; Navarro-Pando *et al*, 2021), purine metabolism remains a candidate contributor to ovarian inflammation. Circulating metabolite shifts after parabiosis and the age-associated rise in circulating urate in women instead suggest that purine metabolism is a systemic feature of reproductive aging.

The responses of recipient organs place the aging ovary within inter-organ communication. Donor ovary age changed molecular features of aging in a tissue-specific manner across the brain, liver, uterus and pituitary. This is consistent with clinical associations of premenopausal oophorectomy with later cognitive impairment (Rocca *et al*, 2007) and cardiovascular mortality (Rivera *et al*, 2009), and with the extended lifespan and reduced inflammation of mice receiving young or germ cell-depleted ovarian grafts (Habermehl *et al*, 2019; Mason *et al*, 2009). Ovarian replacement could influence recipient tissues through direct ovarian signals, including steroids and peptide hormones, and through secondary changes in pituitary feedback, because estradiol and inhibin regulate gonadotropin secretion (Gregg *et al*, 1991). FSH, for example, promotes amyloid and tau pathology in mouse models of Alzheimer’s disease (Xiong *et al*, 2022). In aged ovariectomized macaques, estradiol replacement provides behavioral and molecular benefits (Cervera-Juanes *et al*, 2024; Kohama *et al*, 2016) but did not restore occipital cortical transcriptomic age, suggesting that preserving some tissues requires ovarian factors beyond estradiol. Comparisons of ovarian grafts with multi-hormone replacement could identify which signals support individual recipient tissues.

Heterochronic parabiosis separated systemic signals from surgical engraftment. Young intact ovaries acquired age-associated inflammatory signatures when exposed to an older circulation, indicating that systemic factors can shift ovarian aging signatures without transplantation. Older parabiont ovaries, however, did not show the transcriptomic rejuvenation of aged ovaries transplanted into young hosts. Differences in animal age, the age gap between partners and the timing of collection could contribute (Zhang *et al*, 2023), but this result suggests that shared circulation alone is insufficient for robust ovarian rejuvenation and that new vascular and tissue connections within a young host may contribute.

This study has limitations. All molecular measurements were made in bulk tissue, so shifts in estimated age combine regulation within resident cells with changes in cell composition, and cell type-resolved clocks will be needed to separate the two. These changes may include host-derived immune, vascular and stromal cells that enter a graft after revascularization. Donors and recipients were of the same inbred strain, so the graft origin of offspring was inferred from removal of both host ovaries rather than confirmed genetically. Finally, the parabiosis partners differed by four months of age, which may limit the rejuvenation detectable in the older partners.

Collectively, these findings support a model of constrained plasticity in which the aging ovary remains responsive to the physiological state of its host. The aged ovary retains latent developmental potential that may be partly obscured by local inflammation and stromal remodeling, and changes in ovarian age extend beyond reproduction to other organs. Identifying the signals that mediate this communication may inform non-surgical strategies to emulate a youthful systemic environment.

## Methods

### Animals and ethics

Female mice were housed under specific pathogen-free conditions on a 12 h light-dark cycle with *ad libitum* access to food and water. All procedures were approved by the Institutional Animal Care and Use Committee of Brigham and Women’s Hospital. For transplantation, young C57BL/6 females were obtained from Charles River Laboratories and aged C57BL/6JNia females from the National Institute on Aging (NIA) aged rodent colony. The NIA supplies the birth month of each animal, and exact ages were counted from receipt and confirmed with tail-tattoo identifiers.

### Ovarian lifespan series

Ovaries were collected from 80 female mice at 18 ages between 0.5 and 17 months (3-7 mice per age; Fig. 1A), and brain and liver were collected from the same animals for the cross-tissue analyses (Fig. 1D and Extended Data Fig. 1D,E). Every ovary was profiled by bulk RNA sequencing and by DNA methylation arrays.

### Orthotopic ovary transplantation

Ovaries were transplanted between young and old females to test the effect of systemic host age on ovarian aging. Heterochronic groups received an ovary from a donor of the other age class, and isochronic groups received an ovary from a donor of the same age class and genetic background. Group names give donor age before recipient age, so that old-in-young denotes an old ovary in a young recipient and young-in-old denotes a young ovary in an old recipient.

Animals were anaesthetized with isoflurane delivered through a SomnoSuite low-flow anaesthesia system (Kent Scientific), with induction in a chamber and maintenance through a nose cone. They were kept on a homeothermic warming pad with a rectal temperature probe, and body temperature was kept above 34 °C. Ophthalmic lubricant was applied, and the dorsal surgical site was shaved, treated with depilatory cream and disinfected with povidone-iodine, which was dried before incision. Animals were draped and positioned laterally to avoid respiratory compression. The ovary, located caudal to the kidney within the ovarian fat pad and enclosed by the bursa, was reached through a small dorsolateral incision of the skin and muscle layer, exteriorized with ring forceps and kept moist with warm sterile phosphate-buffered saline. Under a stereomicroscope, a small slit was made in the bursa with spring scissors next to the fat pad on its caudal side, the recipient ovary was removed, and the donor ovary was placed into the bursa in its anatomical orientation without sutures (Parkening *et al*, 1985). Donor ovaries were harvested immediately before grafting, from euthanized donors of the required age class or from the reciprocal recipient of a heterochronic exchange, and kept in sterile saline. In unilateral procedures one ovary was replaced and the contralateral ovary remained *in situ* as a within-animal endogenous control. In bilateral procedures both ovaries were replaced. The reproductive organs were returned to the peritoneal cavity, the muscle layer was closed with a continuous absorbable suture and the skin with non-absorbable sutures or wound clips. Sham-operated animals underwent the same incision, exteriorization and handling, after which the ovaries were returned to their original position without transplantation.

Surgeries and tissue collections were performed between 10:00 and 16:00 on consecutive days, in mixed batches of young and old animals, to limit differences in age at surgery and to standardize collection across groups. Under anaesthesia, animals received extended-release buprenorphine (Ethiqa XR, 3.25 mg kg⁻¹ subcutaneously), which provides analgesia for approximately 72 h, and meloxicam (5 mg kg⁻¹ subcutaneously), given at least 30 min apart. Meloxicam was repeated 24 h later, with a third dose after a further 24 h when needed, at doses based on the body weight measured that day. After surgery each animal received 0.5-1 ml of warm sterile saline subcutaneously and recovered on a warming pad with hydration gel and moistened food on the cage floor. Body weight and condition were recorded daily for 7 days. In the reproductive cohort, young recipients were 3-4 months old and old recipients 13 months old at surgery, and estrous stages were recorded and mating began 4-5 months later, at 7-9 and 17-18 months of age (Fig. 7C and Table 1). In the molecular cohort, recipients were 4 (young) or 13-14 (old) months old at surgery, and tissues for molecular and histological analyses were collected 4-5 months later, at 8-9 and 17-19 months of age (Fig. 4A).

**Table 1.** Ages and timeline of the reproductive cohort.

|  | Young recipients | Old recipients |
| --- | --- | --- |
| **Mating before transplantation (one round; HyperOva and hCG)** | Before surgery (age not recorded) | 10-12 months, 1-2 months before surgery |
| **Age at bilateral transplantation** | 3-4 months | About 13 months |
| **Age of donor ovaries at grafting** | Young ovaries (Y in Y) 3-4 months; old ovaries (O in Y) about 13 months | Young ovaries (Y in O) 3-4 months; old ovaries (O in O) about 13 months |
| **Interval from transplantation to estrous recording and mating** | 4-5 months | 4-5 months |
| **Age at estrous recording and mating after transplantation** | 7-9 months | 17-18 months |
| **Chronological age of grafted ovaries at mating** | Y in Y 7-9 months; O in Y 17-18 months | Y in O 7-9 months; O in O 17-18 months |
| **Mating period after transplantation** | Several rounds over 4-8 weeks | Several rounds over 4-8 weeks |
| **Recipients (Fig. 7E-G)** | Y in Y n = 7; O in Y n = 7 | Y in O n = 6; O in O n = 6 |

Group names give donor ovary age before recipient age. Donor ovaries came from animals of the same age classes as the recipients.

### Heterochronic parabiosis

Young (3.5 months) and old (7.5 months) females were joined as heterochronic pairs, and age-matched young-young and old-old pairs served as isochronic controls (six young isochronic, six heterochronic and four old isochronic pairs; Fig. 4A and Extended Data Fig. 15A). Pairs were matched for body size and co-housed for 1-2 weeks before surgery (Kamran *et al*, 2013). Joining and separation followed published protocols (Kamran *et al*, 2013; Zhang *et al*, 2023). Both partners were anaesthetized with isoflurane and placed side by side in lateral recumbency rather than on their backs, which avoids respiratory compromise, particularly in old animals. After shaving and povidone-iodine disinfection, matching longitudinal skin incisions were made along the opposing flanks from the elbow to the knee, the olecranon and knee joints of the two partners were joined with 2-0 non-absorbable sutures, and the skin of the two animals was sutured together with the inner surfaces apposed. Analgesia and post-operative care were as for transplantation. In addition, each partner received 5% glucose in lactated Ringer’s solution subcutaneously (at least 0.6 ml per day in two doses) for 7 days, and pairs were weighed daily. Pairs with continued weight loss, hunched posture, hypothermia or poor coordination were warmed and given additional fluids and food on the cage floor. Pairs remained joined for 3 months and were then separated under isoflurane anaesthesia by reversing the attachment, and the skin of each animal was closed. Ovaries and blood were collected 1 month after separation, when young and old partners were 7.5 and 11.5 months old. Both partners of each isochronic pair were analysed.

### Reproductive assessment

Reproductive function after bilateral transplantation was assessed in young-in-young (n = 7), young-in-old (n = 6), old-in-young (n = 7) and old-in-old (n = 6) recipients (Table 1). All recipients had been through one round of mating before transplantation, after superovulation as described below. In old recipients, mating and pregnancy took place 1-2 months before surgery. Breeding history before surgery was recorded for the cohort rather than for each animal. Estrous stage was assigned from the appearance of the vaginal opening (Byers *et al*, 2012), distinguishing proestrus or estrus from metestrus or diestrus, and recorded at six serial observations (C1-C6, Fig. 7C). For mating, females were superovulated with CARD HyperOva (inhibin antiserum and equine chorionic gonadotropin; Kyudo) followed 48 h later by human chorionic gonadotropin, according to the manufacturer’s protocol (Takeo & Nakagata, 2015), and were paired overnight with sexually mature males aged 4-8 months. Copulatory plugs were scored the next morning. Several rounds of superovulation and mating were performed over 4-8 weeks. Females with plugs were followed for pregnancy, full-term pregnancy was recorded when pregnancy proceeded to parturition, and litter size was recorded for each recipient as the mean number of live pups across mating rounds, with zero for recipients without litters. Each recipient was one experimental unit (Fig. 7C-I).

### RNA sequencing and DNA methylation profiling

For the lifespan series, total RNA was isolated from whole ovaries, brain and liver with the RNAqueous kit (Thermo Fisher Scientific). Libraries were prepared and sequenced by Novogene. Messenger RNA was enriched with poly(T) oligo-attached magnetic beads and fragmented, first-strand cDNA was synthesized with random hexamer primers, and libraries were completed by second-strand synthesis, end repair, A-tailing, adapter ligation, size selection and amplification. Libraries were quantified by Qubit and real-time PCR, checked on a Bioanalyzer and sequenced on an Illumina NovaSeq X Plus as 150-bp paired-end reads (target 6 Gb per sample). Ovarian libraries yielded 40-85 million raw reads per sample with at least 91.6% of bases at Q30. RNA integrity was measured by Novogene on an Agilent 5400 system, and samples that failed quality control (low RNA integrity or genomic DNA contamination) were resubmitted. Novogene returned raw reads, which were aligned to the GRCm39 mouse genome and summarized as gene-level counts with Ensembl annotation. Two annotation releases were used, an earlier release (57,010 genes) for the lifespan, transplantation ovary, uterus and pituitary samples and a recent release (78,317 genes) for recipient brain and liver and for parabiosis ovaries. Cross-model ovarian analyses (Fig. 4 and Extended Data Figs 8, 9 and 16) used the updated combined transplantation count matrix of 28,266 Entrez genes, mapped to MGI symbols with org.Mm.eg.db, with unmapped and duplicated symbols removed.

DNA was extracted with the QIAamp DNA Micro Kit (Qiagen), and methylation was profiled with the HorvathMammalMethylChip320 array (Arneson *et al*, 2022) by the Epigenetic Clock Development Foundation (lifespan series, project ET0522; unilateral transplantation, a separate plate). Raw IDAT files were processed with SeSAMe (Zhou *et al*, 2018) using default background correction, dye-bias normalization and detection filtering. Probes with a detection P value above 0.01 in any sample, and control and SNP-associated probes, were removed, leaving 270,319 probes whose β values were used for all analyses. Probes were annotated to genes with ChIPseeker v1.24.0 (Yu *et al*, 2015b) (annotatePeak, default priorities) and TxDb.Mmusculus.UCSC.mm10.knownGene v3.10.0, and were assigned to a promoter when located between 1,500 bp upstream and 100 bp downstream of a transcription start site. CpG-island context was assigned with annotatr v1.14.0 (Cavalcante & Sartor, 2017) using the priority island, shore, shelf, open sea. Probes were overlapped with the universal chromatin-state annotation of the mouse genome (Vu & Ernst, 2023) using findOverlaps in GenomicRanges v1.40.0 (Lawrence *et al*, 2013).

### Public datasets

The following public data were reanalysed. Ovarian clock applications used GSE84078 (Ames dwarf and wild-type ovaries) (Schneider *et al*, 2017), GSE270016 (reconstituted follicles) (Wang *et al*, 2024), GSE232309 (mouse ovarian single-cell atlas) (Isola *et al*, 2024) and GSE130664 (cynomolgus macaque ovarian single-cell atlas) (Wang *et al*, 2020). Laser-capture profiles of multinucleated giant cell (MNGC) regions, macrophages and stroma came from GSE283393 (Converse *et al*, 2025), and human ovarian single-nucleus profiles from GSE202601 (Jin *et al*, 2025). Rhesus macaque cortex from intact, ovariectomized and estradiol-treated animals came from PRJNA1088413 (Cervera-Juanes *et al*, 2024), with human reference signatures from GSE33000 (Narayanan *et al*, 2014), GSE71620 (Chen *et al*, 2016), GSE68719 (Dumitriu *et al*, 2016), GSE64810 (Labadorf *et al*, 2015) and GSE173955. The external ovarian aging meta-signature used GSE310764, GSE253194, GSE283764 and GSE273724. Organ aging references used GSE15129, GSE20411, GDS1311, GSE41018, GSE48911 and GDS2654 (brain), GSE15129, GSE20425, GSE20426 and GSE11097 (liver), GSE296745, GSE98901, GSE236128 and GTEx v8 (uterus), and GSE314817, GSE215110 and GTEx v8 (pituitary). Oocyte aging signatures used GSE85019, GSE128305, GSE140090, GSE159281, GSE159789, GSE163724, GSE173598 and GSE268029, and oocyte intervention responses used GSE153920 (Miao *et al*, 2020), GSE183271, GSE201098 (Cao *et al*, 2022) and GSE270016 (Wang *et al*, 2024). Metabolomic signatures came from Metabolomics Workbench studies ST003188, ST001700, ST001329, ST000090, ST000705, ST002931 and ST001523 (Sud *et al*, 2016). Tissue aging and longevity-intervention transcriptomic signatures were taken from published analyses (Tyshkovskiy *et al*, 2019; Tyshkovskiy *et al*, 2026). Plasma urate data came from UK Biobank (Sudlow *et al*, 2015).

### Ovarian transcriptomic clock (OvaAge)

OvaAge was trained on whole-ovary RNA-seq profiles of the lifespan series (11-516 days of age; the saved models were fitted to 79 ovaries). Genes with at least 10 counts in at least 20% of samples were retained and mapped to Entrez identifiers, counts were normalized with relative log expression factors, log10(x + 1)-transformed and standardized within each sample, and the profiles were restricted to the 18,696 genes of the multi-tissue clock gene list, with absent genes imputed by the training median. Each gene was expressed relative to the median of the youngest (0.5-month) group, and the outcome was chronological age relative to the mean age of that group, in months. An elastic net model with fixed hyperparameters (scikit-learn ElasticNet; α = 0.01, l1 ratio = 0.2) retained 423 genes, and a Bayesian ridge model used all 15,587 genes with data. Fig. 1B shows the elastic net model. Leave-one-out cross-validation with these settings in the 80 lifespan ovaries reproduced the panel (r = 0.96, mean absolute error 1.05 months). The Bayesian ridge model was used in Fig. 2A,E, Fig. 6D,E and Extended Data Figs 2A and 15C, and the elastic net model in Extended Data Figs 15B and 22B.

OvaAge was applied without retraining to bulk ovarian RNA-seq of young and old Ames dwarf and wild-type mice (GSE84078) (Schneider *et al*, 2017) and of reconstituted follicles (GSE270016) (Wang *et al*, 2024), to granulosa-cell pseudobulk profiles of young and old mice (GSE232309) (Isola *et al*, 2024), to somatic ovarian-cortex pseudobulk profiles of young and old cynomolgus macaques (GSE130664) (Wang *et al*, 2020) after ortholog mapping (Extended Data Fig. 2A-D), to transplanted and parabiosed ovaries, and to laser-capture and human pseudobulk profiles (see below). For GSE130664, UMI counts were summed within each combination of animal, age group and annotated cell type in consecutive chunks of at least 10⁶ UMIs, outlying profiles were removed in principal-component space (union of an isolation forest at 10% contamination and a Mahalanobis-distance criterion at the 0.95 quantile), genes with at least 2 counts in at least 5% of profiles were kept, and macaque genes were mapped to mouse orthologs before normalization as above. Several pseudobulk profiles were therefore derived from each of the four young and four old macaques. Predictions in isolated cell compartments or in another species are relative values, because the model was trained on whole mouse ovaries.

### Published transcriptomic clocks, module-specific clocks and clock decomposition

Transcriptomic age (tAge) was estimated with the multi-tissue clocks of Tyshkovskiy *et al* (2026) using the tAge package of the Gladyshev laboratory through its TACO interface. Raw counts were restricted to genes with at least 10 counts in at least 20% of samples, normalized with relative log expression factors (edgeR), log10(x + 1)-transformed and standardized within each sample. For YuGene models the standardized profiles were further transformed with YuGene (Lê Cao *et al*, 2014). Profiles were aligned to the gene list of each model, with absent genes set to zero, and the median profile of the reference group was subtracted from every sample, so that predictions are relative to that group. The models predict relative chronological age (months), expected mortality (log10 hazard ratio) or normalized age (percentage of lifespan, labelled lifespan-adjusted in the package). They were trained on mouse, rodent (mouse and rat; 3,575 control transcriptomes across 26 tissues) or rodent and human compendia by elastic net or Bayesian ridge regression and evaluated by nested cross-validation (Tyshkovskiy *et al*, 2026). Bayesian ridge models also return a predictive standard deviation for each sample, shown as error bars. Models were applied without retraining.

The models used were as follows. Normalized age, rodent compendium, scaled expression, elastic net for macaque cortex (Fig. 3B) and liver (Fig. 3H). Normalized age, rodent and human compendium for brain and uterus (Fig. 3E,K). Bayesian ridge multi-tissue chronological and mortality models for ovarian grafts (Extended Data Fig. 7) and the chronological Bayesian ridge model for pituitary (Extended Data Fig. 12A). Mortality, rodent compendium, YuGene, Bayesian ridge for liver (Extended Data Fig. 13A). Mortality, rodent and human compendium, YuGene, elastic net for uterus (Extended Data Fig. 14A). Chronological, mortality and normalized multi-tissue mouse models with YuGene normalization and elastic net for MNGC captures (Extended Data Fig. 22C-E). For recipient brain and liver, the batch-corrected counts described below were used as input.

Module-specific clocks are models in the same framework that use only the genes of one co-regulated gene module, defined by co-expression network analysis of the rodent compendium (Tyshkovskiy *et al*, 2026). Module clocks of the multi-tissue models were imported from that framework. The innate immunity and inflammation module-specific clock (YuGene-normalized, elastic net) was also applied to the parabiosis ovaries (Extended Data Fig. 15D). For each module clock, predictions of the two compared groups were standardized to zero mean and unit standard deviation and regressed on a group indicator, and the slope (standardized difference, tAge Δ), its standard error and P value were recorded, with Benjamini-Hochberg adjustment across modules. Three modules of poor predictability were excluded by the package, leaving 23 modules in the mouse and rodent sets and 14 in the rodent and human set. Normalized slopes of module-clock predictions against chronological age in brain, liver and ovary are shown in Fig. 1D, the additional slope of mortality module clocks during the postnatal window (0.5-2 months) relative to the adult slope in Extended Data Fig. 1B (bars coloured by slope), and predictions of the innate immunity and inflammation module across age in Extended Data Fig. 1C. Standardized group differences are shown for the macaque cortex (mortality, rodent and human compendium, scaled, elastic net; 14 modules; Fig. 3C), brain (mortality, rodent compendium, scaled, elastic net; Fig. 3F), liver and uterus (mortality, rodent compendium, elastic net; Fig. 3I,L), the macaque ovary (Extended Data Fig. 2E) and MNGC captures (Extended Data Fig. 22F).

Enrichment of module genes and of Hallmark and KEGG gene sets was estimated in TACO with random-effects meta-analysis (Viechtbauer, 2010). Gene-specific slopes (or group differences) and their standard errors within a gene set were combined with a restricted maximum likelihood estimator, giving a summary effect and P value for each set. P values were adjusted across gene sets with the Benjamini-Hochberg method (Benjamini & Hochberg, 1995), and the enrichment score was the signed −log10 adjusted P value. This score is distinct from a GSEA normalized enrichment score.

The contribution of a gene to a clock prediction was the clock coefficient multiplied by the gene’s relative expression, on the expression scale used to fit the clock. The contribution of a gene set was the sum of the contributions of its member genes. For lifespan slopes, the contribution of a gene was its coefficient multiplied by its expression slope with age, and the contribution of a gene set was the slope with age of the summed, coefficient-weighted expression of its genes. For group comparisons, the difference in contribution between groups was standardized by its standard error. Gene-set contributions were computed for co-regulated modules and for Hallmark and KEGG gene sets, and Gene Ontology terms were not used. TACO computes contributions for elastic net models only. Module contributions to the age slope of the composite clocks in the ovarian lifespan series are shown in Fig. 1C (23 modules). Gene-set contributions to group differences are shown for brain (normalized age, mouse compendium, YuGene; Extended Data Fig. 11A), liver (normalized age, rodent compendium, scaled; Extended Data Fig. 13B), uterus (normalized age, rodent and human compendium, YuGene and scaled; Extended Data Fig. 14B,C) and MNGC captures (mortality models; Extended Data Fig. 22G-I). Bars show standardized group differences in contribution with error bars, bar colour shows the enrichment score of the gene set, and the displayed gene sets were selected from the TACO output for each comparison.

### Gene expression changes with age and functional enrichment

Age-associated expression in the ovarian lifespan series was estimated with edgeR (Robinson *et al*, 2010) by fitting a negative binomial generalized linear model with age in days as a continuous covariate and testing with a likelihood-ratio test, giving 7,377 of 16,011 genes at a false discovery rate (FDR) below 0.05 (Extended Data Fig. 1A).

Pre-ranked gene set enrichment analysis (GSEA) (Subramanian *et al*, 2005) of age-associated expression was performed with fgsea (Korotkevich *et al*, 2021) against MSigDB Hallmark (Liberzon *et al*, 2015), KEGG (Kanehisa *et al*, 2021) and Reactome (Gillespie *et al*, 2022) collections. Columns represent the ovarian and brain lifespan signatures of this study and published liver, brain and kidney aging meta-signatures (Tyshkovskiy *et al*, 2026). Genes were ranked by their Spearman correlation with age. P values were Benjamini-Hochberg-adjusted within each signature and collection, pathways with adjusted P < 0.05 in at least one column are shown and NES values are clamped to [−3, 3] (Fig. 1E). Spearman correlations between the gene-level age-slope signatures of ovary, brain and liver are shown in Extended Data Fig. 1E. Multi-tissue chronological clock predictions in ovary, brain and liver are plotted against age in Extended Data Fig. 1D.

### Comparison of transplantation signatures with aging and longevity signatures

Differential expression between transplantation groups was estimated with DESeq2 (Love *et al*, 2014) (Wald test). Each transplantation signature was compared with published tissue aging, mortality and longevity-intervention signatures (Tyshkovskiy *et al*, 2019; Tyshkovskiy *et al*, 2026) (Fig. 2B,F). For each pair of signatures, genes were ranked by P value within each signature, the top N genes of each were combined, and Spearman correlation of log2 fold changes was computed across the union. Candidate values of N from 100 to 1,000 in steps of 50 were evaluated, and the value giving the largest number of correlations with Benjamini-Hochberg-adjusted P < 0.05 and |ρ| > 0.1 was used (N = 400). Because N was selected using the displayed correlations, these comparisons are exploratory. P values were adjusted jointly across all pairs, and rows and columns were clustered by complete linkage with distance (1 − ρ)/2.

Functional enrichment of transplantation contrasts was assessed by pre-ranked GSEA with genes ranked by the DESeq2 Wald statistic, using Hallmark, KEGG and Reactome collections, alongside the published aging and intervention signatures (Fig. 2C and Extended Data Fig. 5D). NES values are clamped to [−3, 3].

### Principal component analysis and PERMANOVA

PERMANOVA (Anderson, 2001) on Euclidean distances with 9,999 permutations was used throughout, with Benjamini-Hochberg adjustment of pairwise tests. Analysis-specific settings were as follows.

Ovarian transplantation (Extended Data Fig. 5A,C). Principal components were computed from normalized, log-transformed ovarian expression profiles, separately for the unilateral and bilateral experiments.

#### Recipient brain (Fig. 3D)

One old-in-old sample was excluded because its mean pairwise correlation with other samples was 0.65, compared with about 0.93 for all others, leaving 23 samples (young-in-young 3, old-in-young 7, young-in-old 8, old-in-old 5). Genes with at least 10 counts in at least 3 samples were retained (22,516 genes), counts were transformed with the DESeq2 variance-stabilizing transformation (VST; blind = FALSE), and processing date and recipient cohort were removed with limma removeBatchEffect (Ritchie *et al*, 2015) with the group design protected. Principal components were computed on all genes with centring and without scaling. PERMANOVA used all genes and a custom R implementation (seed 42). Ellipses show 95% regions (multivariate t).

#### Recipient liver (Fig. 3G)

All 24 samples were retained (young-in-young 3, old-in-young 7, young-in-old 8, old-in-old 6). Genes with at least 10 counts in at least 3 samples were retained (17,686 genes), transformed by VST (blind = TRUE) and corrected for processing date with removeBatchEffect. Principal components were computed with centring and without scaling. Ellipses are 95% normal-theory regions.

#### Recipient uterus (Fig. 3J)

One old-in-young sample (M23) was excluded as an outlier, leaving 20 samples (3, 7, 7 and 3). Genes with at least 10 counts in at least 3 samples were transformed by VST (blind = FALSE), and recipient cohort and dissection date were removed with removeBatchEffect with the group design protected. The 500 genes with the highest variance were standardized and analysed with scikit-learn (Pedregosa *et al*, 2011). PERMANOVA used the same 500 standardized genes (seed 42). Ellipses are 95% normal-theory regions.

Macaque cortex, MNGC laser-capture profiles and DNA methylation are described in the corresponding sections below.

### Differential expression and pathway activity in recipient organs

Whole brain (n = 23), liver (n = 24), uterus (n = 20) and pituitary (n = 21; young-in-young 3, old-in-young 8, young-in-old 7, old-in-old 3) of bilateral transplant recipients were analysed with DESeq2 (Love *et al*, 2014) using design group. Wald contrasts were old-in-young versus young-in-young (effect of an old ovary in a young recipient), young-in-old versus old-in-old (effect of a young ovary in an old recipient) and young-in-old versus young-in-young (recipient age under a young ovary). The Wald statistic was the gene-level statistic for all downstream analyses. Ensembl identifiers were mapped to MGI symbols with Ensembl BioMart and org.Mm.eg.db, and for duplicated symbols the entry with the largest absolute statistic was kept. For brain and liver, DESeq2 was run on batch-corrected counts, obtained by back-transforming the limma-corrected VST values to the count scale and rounding. For uterus, it was run on rounded back-transformed batch-corrected VST values, and for pituitary on rounded size-factor-normalized counts. In old recipients, recipient cohort coincided with group, so cohort effects cannot be separated from the young-in-old versus old-in-old contrast in brain, liver and uterus.

External aging references formed the first column of each organ heatmap (Extended Data Figs 11-14). For brain and liver, public microarray datasets were analysed with limma (old versus young) after log2 transformation where needed, probes were annotated with platform tables and the probe with the largest |t| was kept per gene, and a meta t statistic was computed as the mean of per-contrast t statistics weighted by the square root of sample size, retaining genes measured in at least two contrasts. The brain reference combined eight contrasts from GSE15129, GSE20411, GDS1311, GSE41018, GSE48911 and GDS2654 (21,541 genes), and the liver reference combined nine contrasts from GSE15129, GSE20425, GSE20426 and GSE11097 (23,539 genes; rat symbols matched to mouse by identity). For uterus and pituitary, each dataset was compared old versus young as the log2 fold change of log2(CPM + 1) or log2(TPM + 1), and the reference statistic was the mean log2 fold change across datasets for genes present in at least two (uterus GSE296745, GSE98901, GSE236128 and GTEx v8; pituitary GSE314817, GSE215110 and GTEx v8).

Pre-ranked GSEA used fgsea (Korotkevich *et al*, 2021) (nPermSimple = 10,000, seed 42, gene sets of 15-500 genes) with MSigDB v2024.1 mouse Hallmark (Liberzon *et al*, 2015) and human KEGG legacy (Kanehisa *et al*, 2021) collections, with KEGG symbols converted to mouse by case conversion (brain and liver) or an ortholog table (uterus and pituitary). P values were Benjamini-Hochberg-adjusted within each contrast and collection, and heatmaps show gene sets with adjusted P < 0.05 in at least one column, NES clamped to [−3, 3] and rows ordered by Ward.D2 clustering (Extended Data Figs 11C,D, 12B,C, 13D,E and 14E,F).

Signaling pathway activities were inferred from mouse PROGENy footprints (Holland *et al*, 2020; Schubert *et al*, 2018) with the multivariate linear model in decoupleR (Badia-i-Mompel *et al*, 2022) (minimum 5 targets), using the top 100 responsive genes per pathway for brain and liver and the top 500 for uterus and pituitary (Extended Data Figs 11B, 12E, 13C and 14D). Transcription factor activities were inferred with the univariate linear model in decoupleR from mouse DoRothEA regulons of confidence levels A-C (Garcia-Alonso *et al*, 2019; Holland *et al*, 2020) (12,895 interactions, 273 TFs), excluding factors with fewer than 5 targets. TFs with nominal P < 0.05 in at least two columns are displayed (Extended Data Figs 11E, 12D and 13F). Significance marks on PROGENy and TF heatmaps are nominal P values.

For the pituitary metabolic panels (Extended Data Fig. 12F,G), transplantation contrasts were ranked by limma moderated t statistics computed on log2(normalized counts + 1), and the reference was a mouse-only pituitary signature formed by averaging per-gene statistics from GSE314817 (6 versus 27 months, DESeq2 with sex as covariate) and GSE215110 (limma). GSEA used fgsea with gene sets of 10-500 genes against Recon3D subsystems mapped to mouse orthologs (94 sets) (Brunk *et al*, 2018) and a metabolic subset of KEGG legacy (71 sets) (Kanehisa *et al*, 2021).

### Rhesus macaque ovariectomy and estradiol replacement

Bulk RNA-seq of occipital cortex (OC; intact n = 4, ovariectomized (OVX) n = 8, OVX with 17β-estradiol (OVX+E2) n = 6) and dorsolateral prefrontal cortex (PFC; n = 3, 7 and 5) from old female rhesus macaques was reanalysed from PRJNA1088413 (Cervera-Juanes *et al*, 2024). Macaque Ensembl genes (35,432) were mapped to mouse orthologs through Ensembl BioMart (Martin *et al*, 2023). Counts of one-to-many orthologs were assigned to each ortholog and counts of many-to-one orthologs were summed, giving 18,941 mouse genes for all downstream analyses.

Each region was analysed separately with DESeq2 (Love *et al*, 2014) using design group, with the intact group as reference, for OVX versus intact, OVX+E2 versus intact and OVX+E2 versus OVX. Genes were ranked by the Wald statistic and analysed with fgsea _(Korotkevich_ *_et al_*, _2021)_ (gene sets of 15-500 genes, nPermSimple = 10,000, seed 42) against mouse Hallmark, KEGG and Reactome collections from msigdbr 7.5.1, with Benjamini-Hochberg adjustment within each contrast and collection. Using the OVX-versus-intact results, a pathway was classified as concordant when significant (adjusted P < 0.05) in at least one region with NES of the same sign in both regions, and as discordant when the signs differed. A pathway was classified as rescued in a region when significant for OVX versus intact and either not significant or of opposite sign for OVX+E2 versus intact. Heatmaps show pathways significant in at least one region-contrast combination, with NES clamped to [−3, 3] (Extended Data Fig. 10D,E).

TF activities were inferred with the univariate linear model in decoupleR (Badia-i-Mompel *et al*, 2022) (minimum 5 targets) from mouse DoRothEA regulons of confidence levels A and B (Garcia-Alonso *et al*, 2019; Holland *et al*, 2020) (123 TFs), using the DESeq2 Wald statistics of each contrast. P values were Benjamini-Hochberg-adjusted across all TF, contrast and region tests, and TFs with adjusted P < 0.05 in at least one contrast are shown (Extended Data Fig. 10F).

For comparison with human brain aging and neurodegeneration (Extended Data Fig. 10C), reference signatures were derived for Alzheimer’s and Huntington’s disease versus controls (GSE33000, dorsolateral prefrontal cortex) (Narayanan *et al*, 2014), aging (donors over 60 versus under 40 years; GSE71620, Brodmann area 47) (Chen *et al*, 2016), Parkinson’s disease (GSE68719) (Dumitriu *et al*, 2016), Huntington’s disease (GSE64810) (Labadorf *et al*, 2015) and hippocampal Alzheimer’s disease (GSE173955). Array datasets were analysed with two-sample t tests per probe, and author-supplied DESeq2 or likelihood-ratio statistics were used for the sequencing datasets. Macaque signatures were recomputed after retaining genes with at least 10 counts in at least 3 samples, and mouse symbols were converted to human symbols. All signatures were analysed by fgsea (gene sets of 10-500 genes, nPermSimple = 10,000) against human KEGG pathways obtained through the KEGG REST API (352 pathways), and Spearman correlations of NES were computed across shared pathways (344-346). P values are nominal.

For principal component analysis (Fig. 3A and Extended Data Fig. 10A), counts were transformed by VST, each gene was regressed on chronological age and the residuals plus the gene mean formed the age-adjusted matrix, PCA was performed for each region, and groups were compared by PERMANOVA on Euclidean distances with Benjamini-Hochberg-adjusted pairwise tests. Transcriptomic age (Fig. 3B) and module differences (Fig. 3C) were estimated from the mouse-ortholog counts with the models listed above. For Extended Data Fig. 10B, the mean DESeq2 log2 fold change of genes in six curated groups (inflammation and complement, synaptic function, mitochondrial function, glial markers, hormone response, aging and senescence) was tested against zero by one-sample t test across genes in each region and contrast.

### Ovarian signatures across transplantation and parabiosis

Ovarian differential expression for the cross-model comparison (Fig. 4B-F and Extended Data Fig. 16) used DESeq2 (Love *et al*, 2014) fitted separately to each two-group subset, retaining genes with at least 5 counts in at least 3 samples. Bilateral transplantation contrasts compared young-in-old (n = 7) with young-in-young (n = 3) grafts (age acceleration) and old-in-young (n = 6) with old-in-old (n = 6) grafts (age deceleration). Unilateral contrasts compared the transplanted ovary of young-in-old recipients (n = 5) with pooled young isochronic and sham ovaries (n = 15), and the transplanted ovary of old-in-young recipients (n = 3) with pooled old isochronic and sham ovaries (n = 5). Parabiosis contrasts used design plate + group and compared young heterochronic (n = 5) with young isochronic (n = 10) partners and old heterochronic (n = 4) with old isochronic (n = 6) partners. The reference column was the ovarian lifespan signature of this study (7,377 genes at FDR < 0.05, scored as the signed −log10 FDR).

Spearman correlations among the six intervention contrasts and the lifespan signature were computed across the 6,968 genes present in all seven (Fig. 4B) and across the activity scores of the 456 transcription factors scored in all seven (Fig. 4F), with Benjamini-Hochberg adjustment across the 21 pairs of each matrix. Pre-ranked GSEA used fgsea (Korotkevich *et al*, 2021) (gene sets of 15-500 genes, seed 42) with MSigDB 2024.1 mouse Hallmark and Reactome collections and the human KEGG legacy collection matched to mouse symbols. Each heatmap shows the 20 gene sets with the smallest adjusted P value across its four columns, with NES clamped to [−3, 3] (Fig. 4C,D and Extended Data Fig. 16A-D). Pathway and TF activities were inferred with the univariate linear model in decoupleR (Badia-i-Mompel *et al*, 2022) (minimum 5 targets) from mouse PROGENy footprints (100 genes per pathway) (Schubert *et al*, 2018) and mouse CollecTRI regulons (39,961 interactions) (Müller-Dott *et al*, 2023) obtained through OmnipathR (Türei *et al*, 2021). TF heatmaps show the 20 regulators with the smallest nominal P value across the four columns. Significance marks on PROGENy and TF heatmaps are nominal P values (Fig. 4E and Extended Data Fig. 16E,F).

### Ovarian aging meta-signature and metabolic gene-set enrichment

An external ovarian aging reference was derived from four public mouse datasets (Extended Data Figs 8 and 9). GSE310764 (old 18-24 months, n = 7, versus young 2 months, n = 4) and GSE253194 (old 18-25 months, n = 6, versus young) were analysed with PyDESeq2 (Muzellec *et al*, 2023), and GSE283764 (12 versus 6 months, n = 5 each) and GSE273724 (17 versus 7 months, n = 4 each) with Welch t tests on the deposited normalized values. Two-sided P values were converted to Z scores signed by the direction of change and combined by Stouffer’s method with weights equal to the square root of sample size (Whitlock, 2005), over the 24,963 genes present in all four datasets, followed by Benjamini-Hochberg adjustment.

Unilateral transplantation contrasts were recomputed with PyDESeq2 (design condition, Cook’s refitting, at least 10 total counts per gene). Young-in-old grafts (n = 5) were compared with pooled young isochronic and sham ovaries (n = 15; Extended Data Fig. 9), and old-in-young grafts (n = 3) with pooled old isochronic and sham ovaries (n = 5; Extended Data Fig. 8). Entrez gene identifiers were converted to gene symbols before ranking. Pre-ranked GSEA of Wald statistics, or of the meta Z score for the reference, used GSEApy _(Fang_ *_et al_*, _2023)_ (1,000 permutations, seed 42, gene sets of 5-500 genes) against four mouse collections. These were Recon3D subsystems (95 sets) from BiGG reaction annotations mapped to mouse orthologs _(Brunk_ *_et al_*, _2018)_, KEGG metabolic pathways (34 sets) from the KEGG REST API _(Kanehisa_ *_et al_*, _2021)_, MitoCarta3.0 MitoPathways (149 sets) _(Rath_ *_et al_*, _2021)_ and the gene lists of scFEA metabolic modules (168 modules) _(Alghamdi_ *_et al_*, _2021)_. Gene sets with fewer than 3 genes were excluded. Heatmaps show gene sets with FDR < 0.25 or |NES| > 1.5 in either column, up to 25 per collection, with rows ordered by Ward clustering. Significance follows the GSEA convention (FDR < 0.25) _(Subramanian *et al*, 2005)_. These analyses report enrichment of pathway-annotated genes. scFEA flux estimation was not performed, and no metabolic flux was measured.

### Cell-type deconvolution

Cell-type proportions in bulk ovarian RNA-seq were estimated with MuSiC (Wang *et al*, 2019) using the GSE232309 single-cell reference (Isola *et al*, 2024), which contains cells from 3-month (n = 4) and 9-month-old (n = 4) C57BL/6J mice. Published annotations were used, except that phagocyte subpopulations and epithelial subpopulations were each merged to increase population size. Cells from both ages were included to capture age-related variation in cell-type expression. The reference comprised 13 annotated populations (phagocytes, three stromal populations, theca, two granulosa populations, epithelium, T and B lymphocytes, luteal cells, endothelium and oocytes), and the estimated proportions of each sample summed to one. In the lifespan series (0.5-17 months, at least three ovaries per age), the fitted proportions explained the bulk profiles with a mean RZ of 0.377 (s.d. 0.055), and change with age was tested by Spearman correlation with Benjamini-Hochberg correction across cell types (Extended Data Fig. 4). Estimated proportions of each cell type in the unilateral transplantation experiment (n = 38 ovaries, eight groups) were compared by Kruskal-Wallis test, followed by two-sided Dunn’s pairwise tests with Benjamini-Hochberg adjustment over all 28 pairs (Extended Data Fig. 6).

### DNA methylation analyses

Epigenetic age was estimated with the universal pan-mammalian clocks 2 and 3 (Universal2 and Universal3) (Lu *et al*, 2023), using the *Mus musculus* species parameters given by Lu *et al* (2023). Arrays were normalized with SeSAMe, and clock estimates (in years) were computed by the Epigenetic Clock Development Foundation. For these clocks, the Foundation used the probes shared with the 40k mammalian array and assigned missing probes the median of a mouse reference set. Estimates were multiplied by 12 to give months (Fig. 1F, Fig. 2D and Extended Data Fig. 3A). Clock estimates, mean methylation and methylation entropy (Extended Data Fig. 3B,C) were regressed on chronological age by ordinary least squares in the lifespan ovaries, with Pearson correlation. Fig. 1G shows partial Pearson correlations, adjusted linearly for chronological age, among mean methylation, entropy, the two universal clocks, the multi-tissue chronological and mortality transcriptomic clocks and OvaAge in the lifespan ovaries. In the unilateral transplantation experiment, Universal2 estimates were compared across eight groups (n = 36; young sham 5, young isochronic 8, old-in-young graft 3 and contralateral 3, young-in-old graft 5 and contralateral 5, old isochronic 5, old sham 2) by Kruskal-Wallis test, with pairwise Welch t tests Benjamini-Hochberg-adjusted across all 28 pairs and selected pairs displayed (Fig. 2D).

Principal component analysis of β values used the 5,000 most variable CpGs across the lifespan series, with samples grouped into development (0.5-2 months), peak reproductive (3-8 months), early decline (9-12 months), cyclic cessation (13-14 months) and acyclic (15-17 months) stages (Extended Data Fig. 17A), and the 50,000 most variable CpGs across unilateral transplantation samples (Extended Data Fig. 5B).

For differential methylation after transplantation (Extended Data Fig. 17B-D), β values were converted to M values with the beta2m function of lumi v2.40.0 (Du *et al*, 2010) to improve homoscedasticity. Linear mixed models were fitted with limma v3.44.3 (Ritchie *et al*, 2015), modelling M values as a function of transplantation group with array plate position as a fixed effect and mouse as a random effect. Differentially methylated positions (DMPs) were defined by empirical Bayes-moderated t tests at FDR < 0.05. DMPs were called for four unilateral contrasts (young-in-old grafts versus young isochronic ovaries, old-in-young grafts versus old isochronic ovaries, and the contralateral endogenous ovaries of old and of young recipients versus old and young isochronic ovaries; 36 arrays from 24 mice). Hypermethylated and hypomethylated DMPs were summarized as fractions in each CpG-island context, gene feature and chromatin state, alongside all 270,319 analysed probes, without a formal enrichment test.

Pathway-specific methylation age (PathAge) followed the Ageome framework (Ying *et al*, 2024). CpGs were assigned to genes with GREAT regulatory-domain rules (McLean *et al*, 2010), with a basal domain from 5 kb upstream to 1 kb downstream of each transcription start site extended up to 1 Mb to the nearest basal domains, and were grouped into 1,808 Hallmark, KEGG and Reactome pathways. Pre-trained pathway clocks from that framework, elastic net models trained with cross-validation on mouse profiles from the Mammalian Methylation Consortium, were applied without retraining to SeSAMe-normalized β values after mapping probes to the mammalian 40k array manifest, averaging probes at the same genomic position and imputing missing values with the probe mean. MetroAge was the average of all PathAge predictions weighted by the inverse of the squared cross-validated root mean squared error of each model. In the lifespan series (80 ovaries, 2 weeks to 17 months), PathAge was regressed on chronological age by ordinary least squares, and the slope, Pearson r, RZ, mean absolute error and root mean squared error are reported (Fig. 5A,C,D and Extended Data Figs 18-20). In the unilateral transplantation experiment, groups (young isochronic n = 8, old isochronic n = 5, old-in-young grafts n = 3, young-in-old grafts n = 5, and contralateral endogenous ovaries of young and old recipients, n = 5 each) were compared by Kruskal-Wallis test, and five pairs (young isochronic versus old isochronic, young isochronic versus young-in-old grafts, old-in-young grafts versus old isochronic, young-in-old grafts versus old isochronic, and old-in-young versus young-in-old grafts) by two-sided Welch t tests with Benjamini-Hochberg adjustment across the five comparisons within each pathway. For Fig. 5B, MetroAge was calibrated to chronological age by a linear fit across the transplantation samples, and each sample was expressed relative to the median of its matched isochronic group (ΔMetroAge). On each axis, the isochronic, contralateral and graft groups were compared by Kruskal-Wallis test followed by Dunn’s tests with Benjamini-Hochberg adjustment. Chronological age in the lifespan panels is in years. Sham-operated ovaries were not included, and replicate measurements of a sample were averaged.

Promoter methylation and expression were integrated in the 90 ovaries with both measurements (the 80 lifespan ovaries and 10 ovaries from 1-month-old mice of a rapamycin or vehicle treatment cohort), following a multi-omic aging-gene framework (Moqri *et al*, 2026). Age-associated promoter CpGs were autosomal probes annotated to promoters (10 kb upstream to 1 kb downstream of a transcription start site) whose β values changed with age in days in a linear regression (P < 10⁻¹⁰; 8,721 probes in 5,146 genes). Genes with an age-associated change in expression (FDR < 0.05; see above) and at least one age-associated promoter CpG were assigned to four classes by the direction of expression change and the sign of the mean correlation of their promoter CpGs with age. The association between the two directions was tested by χZ test with Yates’ continuity correction, and normalized differences between observed and expected counts are displayed (Fig. 5E). Each class was tested for over-representation of KEGG and Reactome pathways (Fig. 5F). Genes were annotated to ovarian cell types and states with curated marker genes of ovarian cell types and states (Fig. 5G). For Extended Data Fig. 21, every CpG was correlated with age in months (Pearson) across the ovaries with methylation data, and for each listed gene the CpG with the smallest P value among all CpGs annotated to that gene was plotted with a least-squares line, together with the direction of expression change from the lifespan model. Promoter methylation and expression changes across transplantation contrasts for genes annotated to ovarian cell types (Fig. 6A) show log2 fold changes with adjusted P ≤ 0.05. Cell-type annotations refer to genes measured in bulk tissue, not to measurements in purified cells.

### Multinucleated giant cell analyses

Unnormalized counts of laser-capture MNGC regions, macrophages and non-MNGC stroma (four captures each) were obtained from GSE283393 (Converse *et al*, 2025). For principal component analysis (Fig. 6C), genes with at least 10 counts in at least 3 samples were retained (17,932 of 54,232 features), counts were transformed by VST (blind = TRUE) and compartments were compared by PERMANOVA on Euclidean distances (9,999 permutations) with Benjamini-Hochberg-adjusted pairwise tests. Clocks were applied in TACO to the unnormalized counts of the 12 captures, namely OvaAge (Bayesian ridge, Fig. 6D; elastic net, Extended Data Fig. 22B) and the multi-tissue models listed above (Extended Data Fig. 22C-E). The difference between MNGC and stromal captures was decomposed with the chronological multi-tissue mouse clock (YuGene, elastic net) into standardized module differences (Fig. 6F), and with mortality clocks into module (rodent and human compendium, scaled; Extended Data Fig. 22F), KEGG and Hallmark (rodent compendium, YuGene; Extended Data Fig. 22G,H) and gene (rodent compendium, scaled; log10 hazard ratio; Extended Data Fig. 22I) contributions. Group statistics are described under Statistics.

For pathway analyses, Ensembl identifiers were mapped to symbols with org.Mm.eg.db, retaining for duplicated symbols the identifier with the highest total count, and genes with at least 10 counts in at least 3 samples were analysed with DESeq2 (Love *et al*, 2014) (15,740 genes). Genes ranked by the Wald statistic for MNGC versus macrophages and MNGC versus stroma were analysed with fgsea (Korotkevich *et al*, 2021) (gene sets of 15-500 genes, nPermSimple = 10,000, seed 42) against mouse Hallmark and KEGG collections from msigdbr 7.5.1, with Benjamini-Hochberg adjustment within each comparison (Extended Data Fig. 23A,B). PROGENy scores were computed from the VST matrix (top 500 genes per pathway, scaled, 10,000 permutations) (Schubert *et al*, 2018) and compared among compartments by one-way ANOVA with Tukey’s *post hoc* test (Extended Data Fig. 23C). For senescence scores, log2(normalized counts + 1) were z-scored per gene across the 12 samples and averaged over the genes of each set (minimum 5 genes), for SenMayo (Saul *et al*, 2022), a core SASP list, FRIDMAN-SENESCENCE-UP (Fridman & Tainsky, 2008), the Reactome SASP set (Gillespie *et al*, 2022), a WikiPathways senescence set and CellAge senescence inducers mapped to mouse orthologs (Avelar *et al*, 2020), and compared by one-way ANOVA with Tukey’s test (Extended Data Fig. 23D).

For metabolic enrichment (Extended Data Fig. 25), DESeq2 was run on genes with at least 10 total counts (24,271 genes), Wald statistics were mapped to symbols (18,795 genes) and analysed with fgsea (gene sets of 5-500 genes, nPermSimple = 10,000) against the Recon3D, KEGG metabolic, MitoCarta3.0 and scFEA module collections described above, with Benjamini-Hochberg FDR within each collection and comparison. Heatmaps show gene sets with FDR < 0.25 in at least one comparison, up to 50 per collection, with rows ordered by Ward.D2 clustering.

#### Deriving an MNGC signature

To derive an MNGC-specific transcriptional signature, we analysed the GSE283393 bulk RNA-seq counts with PyDESeq2 (Muzellec *et al*, 2023) (design compartment, macrophages as reference, Cook’s distance refitting). PCA of VST counts confirmed separation of MNGC, macrophage and stromal samples (Extended Data Fig. 24A). MNGC-up genes were defined as those significantly upregulated in both the MNGC-versus-macrophage and MNGC-versus-stroma contrasts (adjusted P < 0.05, log2 fold change ≥ 2), with normalized expression in every MNGC replicate above the median of both comparison groups and mean normalized expression ≥ 20 in MNGCs, yielding 216 genes. MNGC-down genes were defined similarly (adjusted P < 0.05, log2 fold change ≤ −2), requiring expression in every MNGC replicate below the macrophage and stromal medians and mean normalized expression ≥ 100 across macrophage and stromal samples, yielding 167 genes (Extended Data Fig. 24B).

Applying the MNGC signature to human snRNA-seq data. Human ovary snRNA-seq data (GSE202601) (Jin *et al*, 2025) were processed in Scanpy (Wolf *et al*, 2018) with standard quality control (mitochondrial content < 10%, at least 100 genes per nucleus and genes detected in at least 5 nuclei), with doublets removed by Scrublet (Wolock *et al*, 2019) at a doublet score threshold of 0.3, normalized to 10,000 counts per nucleus and log-transformed. Donor batch effects were corrected with Harmony (Korsunsky *et al*, 2019) (Extended Data Fig. 24C). The mouse MNGC signature was mapped to human orthologs through MyGene.info HomoloGene records (Xin *et al*, 2016) and used to compute per-nucleus module scores with the Scanpy score-genes function (Tirosh *et al*, 2016), with MNGC score = MNGC-up − MNGC-down (Extended Data Fig. 24G). Immune nuclei were reclustered with the Leiden algorithm (Traag *et al*, 2019) (resolution 0.5) on the Harmony embedding, and macrophage clusters were identified by canonical markers (*LYZ*, *CD68*, *CSF1R*, *C1QA* and *TYROBP*; Extended Data Fig. 24D-F). Within each donor, macrophage nuclei with MNGC scores at or above the donor median were classified as MNGC-like. Raw counts were summed by donor and MNGC-like status, and stromal nuclei were summed by donor, to generate pseudobulk count matrices, which were analysed with the OvaAge Bayesian ridge model in TACO and expressed on TACO’s human-years scale (Fig. 6E). Fig. 6E shows the four donors of the older group (ovary47, ovary51, ovary52 and ovary54), and compartments were compared by two-sided paired t tests across donors with Benjamini-Hochberg adjustment over the three comparisons. Immune clusters were characterized by Wilcoxon rank-sum marker ranking in Scanpy and displayed as variable-scaled mean expression of *MRC1*, *CD163*, *LYZ*, *CD68*, *CSF1R*, *CSF3R*, *CALD1*, *MS4A1*, *PAX5*, *CD247*, *ITK*, *NKG7* and *GNLY* (Extended Data Fig. 24D). The signature-defined group identifies an expression state and does not establish multinucleation.

### Histology and MALDI mass spectrometry imaging

Haematoxylin and eosin (H&E) staining was performed on ovarian cryosections adjacent to those used for MALDI imaging (Fig. 6B) and on sections of transplanted ovaries (Extended Data Fig. 22A), and slides were scanned on a Leica slide scanner.

MALDI Fourier-transform ion cyclotron resonance mass spectrometry imaging was performed by A. Hossain in N. Agar’s laboratory on ovarian cryosections adjacent to sections stained with haematoxylin and eosin, which were scanned on a Leica slide scanner. Ten ovaries, five from young (7.5 months) and five from old (11.5 months) female mice, were mounted on one slide. They were imaged in negative ion mode at 25 µm pixel size. Ions were assigned by accurate mass as deprotonated [M−H]− species of uric acid (m/z 167.02106, ±5.3 ppm), cysteine (m/z 120.01246, ±3.5 ppm), cystine (m/z 239.01658) and oleic acid (m/z 281.24867, ±9.2 ppm). The young and old labels of two ovaries were corrected against the sample inventory before analysis; the section originally labelled young showed the histology of an aged ovary.

#### Image processing and co-registration

Ion images were provided as whole-slide renders on a shared intensity scale (viridis colour map; 100% = slide-wide reference; higher values clipped). Each ovary was cropped from the slide image and the colour map was inverted to scalar values by nearest-colour lookup. Per-ovary means obtained this way agreed with the laboratory’s per-ovary averages from the raw spectra (Spearman ρ = 1.00 across ovaries), so intensities are relative but comparable across ovaries on the slide. The adjacent H&E section was registered to each MALDI section in three steps: a search over rotation and mirror orientation, with area-matched scale, that maximized overlap of tissue outlines; affine refinement maximizing Mattes mutual information between haematoxylin density and a structural MALDI image built only from ions not tested here (glutamate, ATP, hexose and lactate); and outline refinement. Mirrored solutions were rejected because adjacent cryosections are not mirrored. Among the remaining candidates within 0.03 of the best outline overlap (intersection over union, IoU), the candidate with the highest mutual information was kept; three ovaries with near-tied candidates were resolved by follicle or cyst positions visible in both modalities. Final IoU was 0.77-0.94. One young ovary (IoU 0.80, cysts misaligned) was excluded from regional analyses but retained for whole-section comparisons. Oleic acid, cysteine, cystine and uric acid were never used for alignment.

Histological annotation. Annotation used the registered H&E images, resampled to 1.25 µm per pixel, without reference to ion images. Follicles were outlined manually as circles. MNGC-rich stroma was defined by the colour of the pale-tan to olive-brown, vacuolated giant cell fields and their golden-brown ceroid pigment. Non-white tissue pixels (CIELAB L* < 92, chroma > 6) were classed as MNGC-like when their b* value (yellow-blue axis; Gaussian smoothing σ = 2 µm) exceeded the lower of −3 and the ovary’s median tissue b* plus 8 and their CIELAB hue angle was at least −30°, which excludes eosin-pink and haematoxylin-purple tissue, or when they met a pigment rule (R > G > B + 12, R − B > 45, hue 12-45°, saturation ≤ 0.6). The MNGC-like fraction was smoothed (σ = 8 µm); regions in which it reached 0.3, with holes filled and an area of at least 600 µmZ, were retained after excluding a 40 µm halo around follicles and were mapped to the MALDI grid (at least half of each 10 µm pixel). The same rule labelled 0-0.7% of young sections and 2.5-25% of old sections, so MNGC-rich comparisons were restricted to old ovaries. Sensitivity analyses used b* offsets of 6, 8 and 10 and hue limits of −30°, −25° and −20°.

Quantification and statistics. For each ovary, enrichment was summarized as the log2 ratio of mean ion intensity in follicle cores (circles shrunk by 15 µm) to non-follicular tissue, or in MNGC-rich stroma to other stroma (excluding a 40 µm halo around MNGC-rich regions and follicles); a 30 µm rim at the tissue edge was excluded throughout. Because ion images were rendered with a ceiling (values above 100% clipped; up to 11% of pixels in one ovary), enrichment was also expressed as the probability that a randomly chosen MNGC-rich pixel exceeded a randomly chosen pixel of other stroma (area under the curve, AUC), which is insensitive to clipping. Within-ovary P values were obtained from spatial null distributions (1,000 draws) generated by placing the same follicle circles at random positions within the tissue, or by randomly rotating and mirroring the MNGC-rich region about the tissue centroid. As threshold-free checks, the Spearman correlation between local MNGC-rich area fraction and ion intensity, and between pairs of ion maps (co-localization), were compared with rotation nulls. Follicle-level oleic acid enrichment was compared between young and old ovaries with a linear mixed model with ovary as a random effect. Group-level inference used one value per ovary: two-sided one-sample t-tests of the log2 ratios, with sign counts and Fisher’s combination of the within-ovary P values. Whole-section comparisons used log2 per-ovary mean intensities from the raw spectral averages, compared by Welch’s t-test (five young versus five old ovaries). Cysteine, cystine, uric acid and oleic acid were specified in advance; with Benjamini-Hochberg adjustment across all 77 annotated ions, cystine had q = 0.09. Cysteine and cystine enrichment in MNGC-rich stroma kept the same direction under all alternative annotation thresholds and registrations (not shown). Analyses used Python with scikit-image, SimpleITK, SciPy and statsmodels.

### Parabiosis plasma metabolomics

Blood was collected one month after separation from young isochronic (YI, n = 11), young heterochronic (YH, n = 6), old heterochronic (OH, n = 6) and old isochronic (OI, n = 7) parabionts. Samples were spiked with 1 µM chlorpropamide as internal standard and analysed in triplicate by liquid chromatography-mass spectrometry in positive ion mode on an HSS T3 column (300 µm × 150 mm) with a 30 min water-acetonitrile-formic acid gradient, across seven batches. Features were annotated against a library of 465 serum metabolites and 22 targeted compounds.

Each intensity was divided by the chlorpropamide intensity of the same sample, replicate and batch, log2-transformed after adding half the smallest positive normalized value, and technical triplicates were collapsed to their median. Within each batch one feature was retained per annotated metabolite, prioritizing MS/MS confirmation, then annotation scores, detection rate and mean intensity, giving 341 metabolites (Tier 1, MS/MS-supported, 248; Tier 2, high annotation scores, 80; Tier 3, 13), of which 340 were detected in at least 50% of the samples of at least one group. For each contrast (OI versus YI, OH versus OI, YH versus YI and OH versus YH), groups were compared by two-sided Mann-Whitney U tests with at least 2 values per group, fold changes were differences of group medians of log2 intensities, and P values were Benjamini-Hochberg-adjusted within each contrast. Volcano plots mark nominal P < 0.05 (Fig. 6G,H).

Fifteen external signatures were derived from Metabolomics Workbench studies (Sud *et al*, 2016) as log2 ratios of group means. These covered aging (ST003188 plasma, 24 versus 3 months; ST001700 serum; ST001329 heart) and interventions (ST000090 caloric restriction, rapamycin and acarbose; ST000705 GHRKO, Snell dwarf and PAPP-A knockout; ST002931 exercise; ST001523 *Sirt1*, *Sirt6* and *Sirt1* plus *Sirt6* overexpression; ST001329 NMN and SS-31). Metabolite names were harmonized by lower-casing and removing adduct annotations, bracketed text, numeric suffixes and asterisks, and duplicate entries were averaged, giving 133 metabolites shared with at least one external dataset. Spearman correlations among all 19 signatures were computed on pairwise-complete metabolites and are displayed where at least 10 metabolites were shared (Fig. 6I). Scatter plots show matched log2 fold changes with a least-squares line, Spearman ρ and Pearson r (Extended Data Fig. 26A-F), and the heatmap shows the 50 metabolites shared with at least five external signatures (Extended Data Fig. 26G). Correlation P values are nominal. Benjamini-Hochberg adjustment across the 60 parabiosis-versus-external tests gave a minimum FDR of 0.21. Some external comparisons share control groups.

### UK Biobank urate analysis

UK Biobank (Sudlow *et al*, 2015) participants recorded as female were included if a baseline urate measurement (field 30880, µmol/L) was available (n = 254,095). Urate was natural-log-transformed. Its relationship with age at the first assessment visit was modelled with an unadjusted generalized additive model with a thin-plate regression spline (mgcv default basis dimension) fitted by restricted maximum likelihood (Wood, 2011), with 95% confidence bands from predictions that include smoothing-parameter uncertainty. A linear model estimated the age coefficient with adjustment for body mass index (field 23104), Townsend deprivation index (field 22189), IPAQ physical activity group (field 22032) and assessment centre (field 54) in participants with complete data (Fig. 6J).

### Oocyte aging meta-signatures

Oocyte aging signatures were derived from eight mouse oocyte RNA-seq datasets (257 samples), namely GSE85019, GSE128305, GSE140090, GSE159281, GSE159789, GSE163724, GSE173598 and GSE268029. Genes with at least 10 counts in at least 5 samples were retained, duplicate symbols were collapsed to the most highly expressed entry and expression was transformed to log2(CPM + 1). Within each dataset, the Spearman correlation of each gene with age was computed, two-sided P values (floored at 10⁻¹⁵) were converted to signed Z scores capped at |Z| = 8, and Z scores were combined by Stouffer’s method weighted by the square root of sample size (Whitlock, 2005), with the weighted mean ρ retained as the effect size and Benjamini-Hochberg adjustment. The pan-oocyte signature combined all eight datasets (1,889 genes at FDR < 0.05) and the germinal-vesicle (GV) signature combined the five datasets with GV oocytes of more than one age (GSE85019, GSE159281, GSE159789, GSE173598 and GSE268029; 218 samples; 2,756 genes at FDR < 0.05).

Intervention responses were computed as log2 ratios of group-mean counts per million (pseudocount 1) for oocytes from aged mice after NMN supplementation versus aged controls (GSE153920, n = 3 per group) (Miao *et al*, 2020), aged GV oocytes after NMN versus aged GV oocytes (GSE183271, n = 3 per group), aged oocytes matured *in vitro* with metformin versus controls (GSE201098, n = 3 per group) (Cao *et al*, 2022) and aged oocytes in young versus aged reconstituted follicles (GSE270016, n = 6 and 12) (Wang *et al*, 2024). Bilateral transplantation ovaries were taken from the updated combined count matrix, and three contrasts were computed in the same way, namely old versus young controls (pooled isochronic and sham ovaries, n = 10 and 8), old-in-young grafts versus old controls (n = 6 and 10) and young-in-old grafts versus old controls (n = 7 and 10). Genes were matched by upper-case symbol, keeping for duplicated symbols the entry with the largest absolute change. Concordance was the Spearman correlation between the signature effect size and each response across all shared genes (16,464-21,858 genes), with Benjamini-Hochberg adjustment across the 14 tests of the displayed matrix (Fig. 7A). For Fig. 7B, oocyte signatures were ranked by effect size and bilateral contrasts by the log2 ratio of group-mean CPM defined above, and analysed by pre-ranked GSEA with GSEApy (Fang *et al*, 2023) (1,000 permutations, seed 42, gene sets of 10-1,000 genes) against the 77 MitoCarta3.0 MitoPathways with at least 10 genes (Rath *et al*, 2021), with mouse symbols matched to MitoCarta symbols by case. The heatmap shows the 15 pathways with the largest summed |NES| across the bilateral aging and old-in-young contrasts among pathways with FDR < 0.25 in at least one of these two contrasts.

### Statistics and reproducibility

Tests are two-sided unless stated. For clock panels generated in TACO, transcriptomic age was modelled as a function of group across all samples in the analysis (metafor rma.uni with restricted maximum likelihood, using per-sample predictive standard errors where available), pairwise contrasts were derived with emmeans, and P values were Benjamini-Hochberg-adjusted across the comparisons drawn in each plot. For elastic net predictions the printed P values were reproduced exactly by an ordinary least-squares model of prediction on group fitted to all samples in the run, with Benjamini-Hochberg adjustment across the displayed comparisons. Where a panel combines comparisons against two reference groups, each set was adjusted separately. These P values describe differences in model predictions and do not account for the number of animals beyond the residual variance. Boxplots show the median and interquartile range with individual samples. Benjamini-Hochberg correction (Benjamini & Hochberg, 1995) was applied within the families stated in each section. For Fig. 7H, binary outcomes were pooled by recipient age across both donor-age classes, each recipient contributing one observation, and compared with two-sided Fisher’s exact tests on 2 × 2 tables. Sample odds ratios were 8.4 for copulatory plugs (P = 0.0375) and 12.0 for full-term pregnancy and live birth (P = 0.0138). Live birth after grafting old ovaries into young versus old recipients (5 of 7 versus 0 of 6) and after grafting young versus old ovaries into old recipients (4 of 6 versus 0 of 6) was compared by two-sided Fisher’s exact tests. These pooled comparisons assess recipient age and do not isolate donor age. Litter sizes were compared by Kruskal-Wallis test followed by pairwise Welch t tests with Benjamini-Hochberg correction (Fig. 7I). Analyses used R 4.3.3 with Bioconductor 3.18 and Python with SciPy (Virtanen *et al*, 2020), statsmodels and scikit-learn (Pedregosa *et al*, 2011). Several analyses were run in Edison Scientific notebooks.

## Data availability

The RNA sequencing and DNA methylation data generated in this study, the parabiosis metabolomics data and the MALDI imaging data will be made publicly available upon publication. UK Biobank data are available to approved researchers through the UK Biobank.

## Code availability

The analysis code, including the OvaAge model coefficients, will be made publicly available upon publication.

## Acknowledgements

This work was supported by grants from the National Institute on Aging to V.N.G. S.I. was supported by the Finnish Cultural Foundation through the Foundations’ Post Doc Pool, the Sigrid Jusélius Foundation, an NIH T32 training grant in Endocrinology at Brigham and Women’s Hospital and the Research Council of Finland.

## Author contributions

S.I. conceived and designed the study, performed the ovary transplantation, parabiosis and reproductive experiments, analysed the data and wrote the manuscript with input from all authors. A.T. and Ad.M. developed OvaAge and analysed transcriptomic data. B.Z. performed the parabiosis experiments. K.Y. performed the PathAge analyses. J.J.A.-L. processed the DNA methylation data. E.J. performed the cell-type deconvolution. H.S. derived the MNGC signature and analysed the human snRNA-seq data. L.G. performed the UK Biobank analyses. A.H. and N.A. performed the MALDI imaging. Y.B. performed sectioning and H&E analysis. F.C., S.Z., D.G., Al.M., J.R.P. and M.M. contributed to data analysis and interpretation. V.N.G. conceived and supervised the study, acquired funding and wrote and edited the manuscript. All authors reviewed and approved the final manuscript.

