## Supplemental Figures for "Systemic control and inter-organ communication of ovarian aging"

This file contains Extended Data Figs 1–25 followed by their legends. References cited in the legends are listed in the main manuscript.

#### Ext Fig. 1

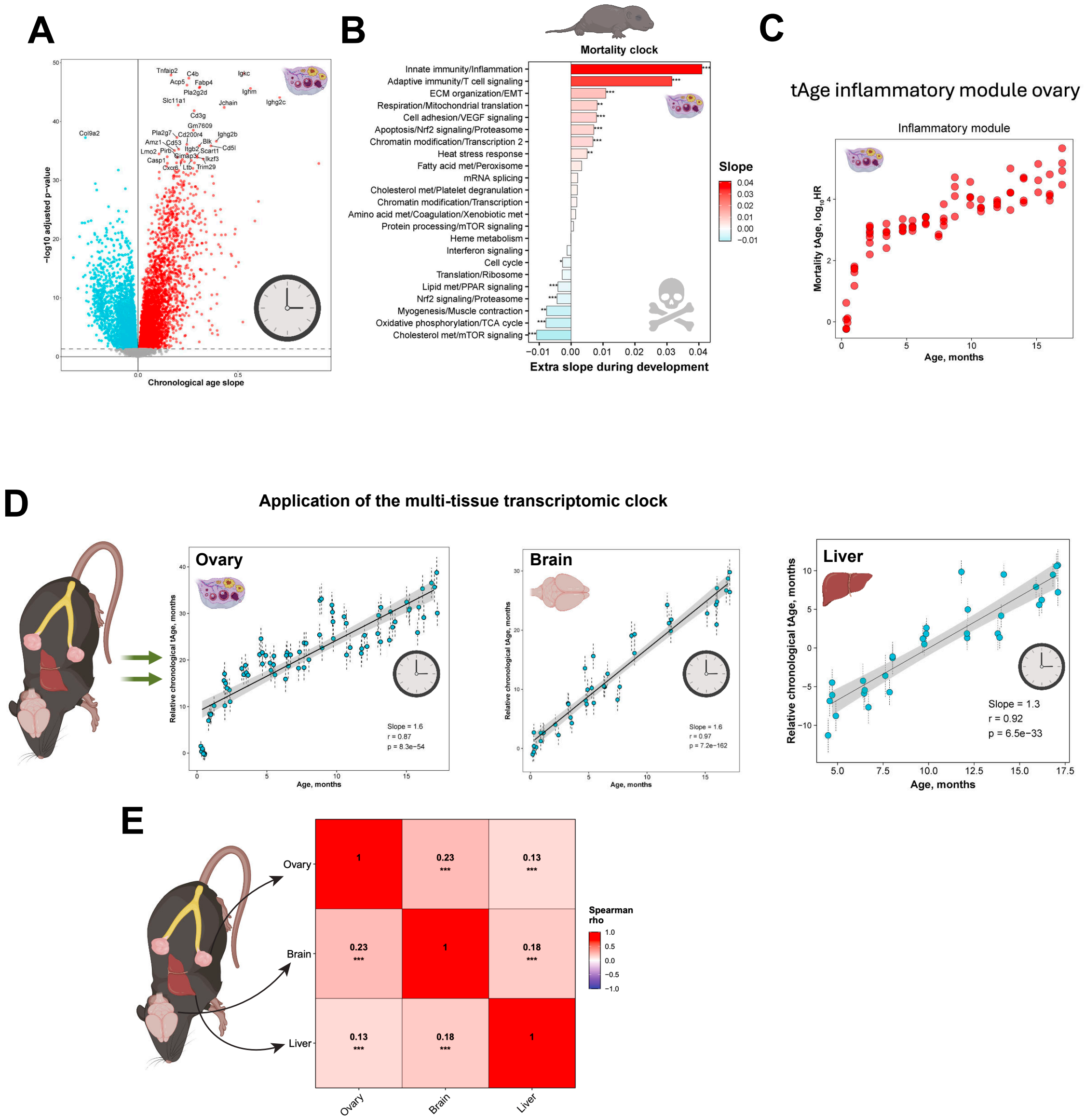

A
Chronological age slope
−log10 adjusted p-value
B
Mortality clock
Innate immunity/Inflammation
Adaptive immunity/T cell signaling
ECM organization/EMT
Respiration/Mitochondrial translation
Cell adhesion/VEGF signaling
Apoptosis/Nrf2 signaling/Proteasome
Chromatin modification/Transcription 2
Heat stress response
Fatty acid met/Peroxisome
mRNA splicing
Cholesterol met/Platelet degranulation
Chromatin modification/Transcription
Amino acid met/Coagulation/Xenobiotic met
Protein processing/mTOR signaling
Heme metabolism
Interferon signaling
Cell cycle
Translation/Ribosome
Lipid met/PPAR signaling
Nrf2 signaling/Proteasome
Myogenesis/Muscle contraction
Oxidative phosphorylation/TCA cycle
Cholesterol met/mTOR signaling
Slope
Extra slope during development
C
tAge inflammatory module ovary
Inflammatory module
Mortality tAge, log10HR
Age, months
D
Application of the multi-tissue transcriptomic clock
Ovary
Slope = 1.6
r = 0.87
p = 8.3e−54
Brain
Slope = 1.6
r = 0.97
p = 7.2e−162
Liver
Slope = 1.3
r = 0.92
p = 6.5e−33
Relative chronological tAge, months
Age, months
E
Ovary
Brain
Liver
Spearman rho

**Ext Fig. 2**

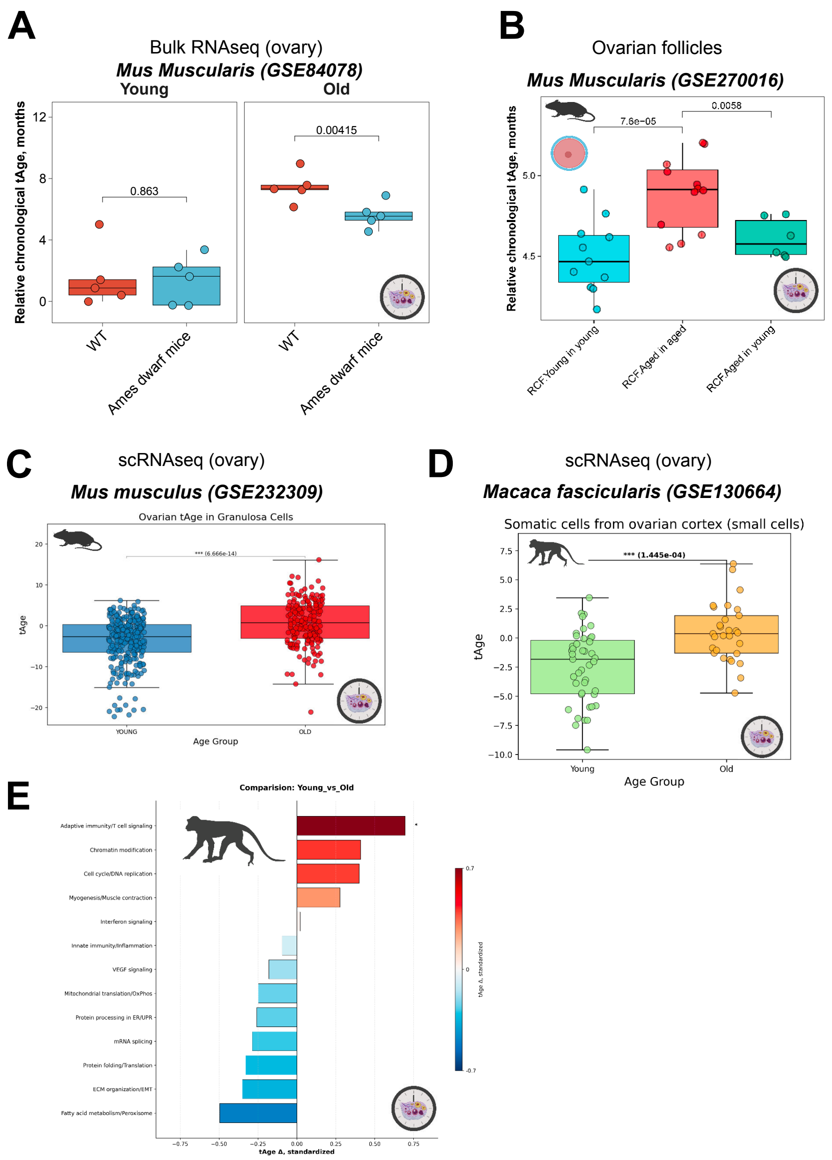
A
Bulk RNAseq (ovary)
Mus Muscularis (GSE84078)
Young
Old
Relative chronological tAge, months
0.863
0.00415
WT
Ames dwarf mice
B
Ovarian follicles
Mus Muscularis (GSE270016)
Relative chronological tAge, months
7.6e−05
0.0058
RCF:Young in young
RCF:Aged in aged
RCF:Aged in young
C
scRNAseq (ovary)
Mus musculus (GSE232309)
Ovarian tAge in Granulosa Cells
tAge
YOUNG
OLD
Age Group
D
scRNAseq (ovary)
Macaca fascicularis (GSE130664)
Somatic cells from ovarian cortex (small cells)
*** (1.445e-04)
tAge
Young
Old
Age Group
E
Comparision: Young_vs_Old
Adaptive immunity/T cell signaling
Chromatin modification
Cell cycle/DNA replication
Myogenesis/Muscle contraction
Interferon signaling
Innate immunity/Inflammation
VEGF signaling
Mitochondrial translation/OxPhos
Protein processing in ER/UPR
mRNA splicing
Protein folding/Translation
ECM organization/EMT
Fatty acid metabolism/Peroxisome
tAge Δ, standardized

#### Ext Fig. 3

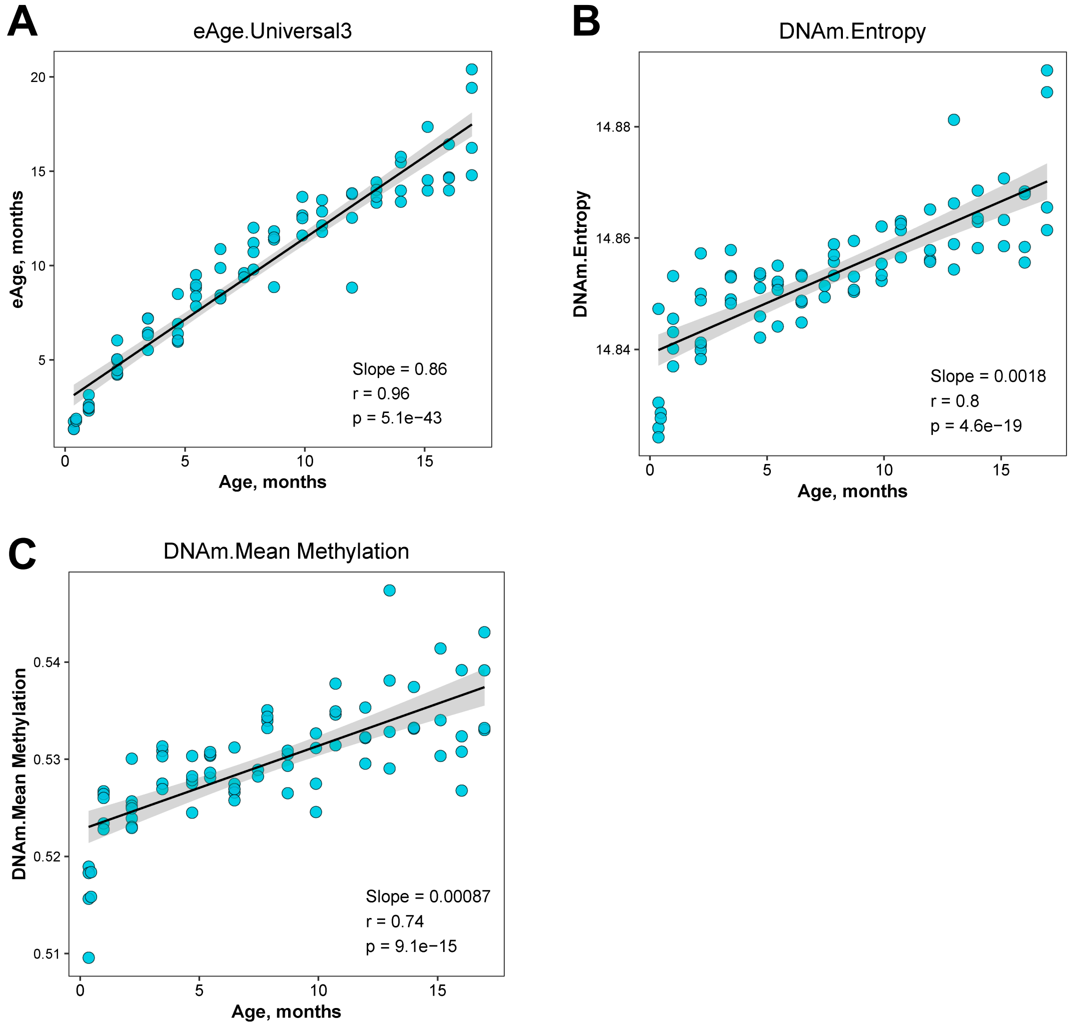
A
eAge.Universal3
eAge, months
Age, months
Slope = 0.86
r = 0.96
p = 5.1e−43
B
DNAm.Entropy
DNAm.Entropy
Age, months
Slope = 0.0018
r = 0.8
p = 4.6e−19
C
DNAm.Mean Methylation
DNAm.Mean Methylation
Age, months
Slope = 0.00087
r = 0.74
p = 9.1e−15

**Ext Fig. 4**

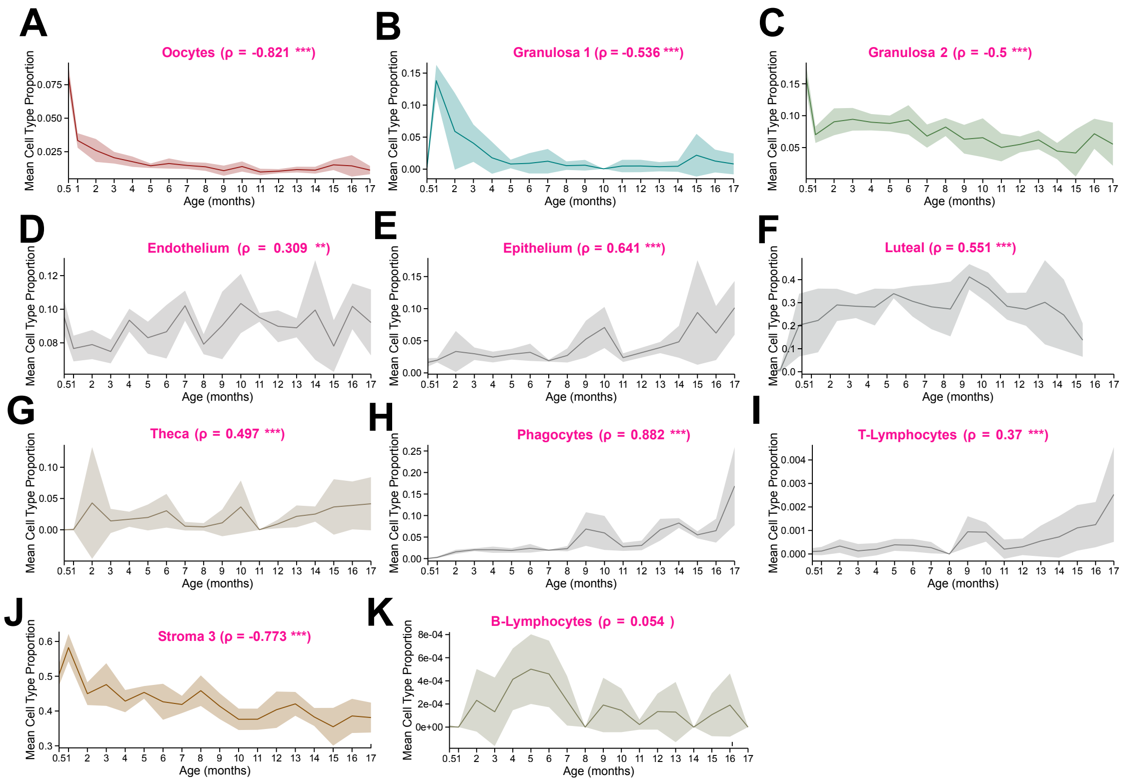
A
Oocytes (ρ = -0.821 ***)
B
Granulosa 1 (ρ = -0.536 ***)
C
Granulosa 2 (ρ = -0.5 ***)
D
Endothelium (ρ = 0.309 **)
E
Epithelium (ρ = 0.641 ***)
F
Luteal (ρ = 0.551 ***)
G
Theca (ρ = 0.497 ***)
H
Phagocytes (ρ = 0.882 ***)
I
T-Lymphocytes (ρ = 0.37 ***)
J
Stroma 3 (ρ = -0.773 ***)
K
B-Lymphocytes (ρ = 0.054 )
Mean Cell Type Proportion
Age (months)

**Ext. Fig. 5**

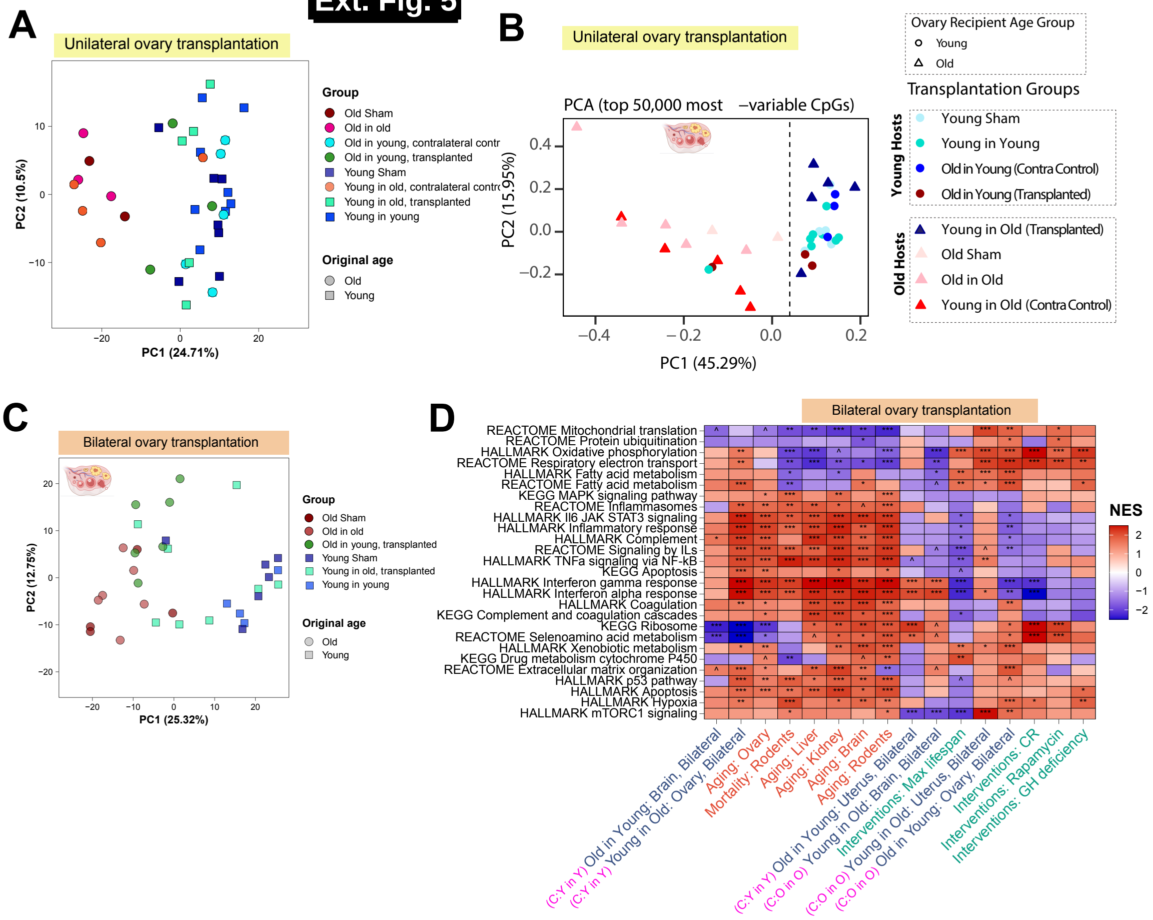

A
Unilateral ovary transplantation
PC1 (24.71%)
PC2 (10.5%)
Group
Old Sham
Old in old
Old in young, contralateral contr
Old in young, transplanted
Young Sham
Young in old, contralateral contr
Young in old, transplanted
Young in young
Original age
Old
Young
B
Unilateral ovary transplantation
PCA (top 50,000 most −variable CpGs)
PC1 (45.29%)
PC2 (15.95%)
Ovary Recipient Age Group
Young
Old
Transplantation Groups
Young Hosts
Young Sham
Young in Young
Old in Young (Contra Control)
Old in Young (Transplanted)
Old Hosts
Young in Old (Transplanted)
Old Sham
Old in Old
Young in Old (Contra Control)
C
Bilateral ovary transplantation
PC1 (25.32%)
PC2 (12.75%)
Group
Old Sham
Old in old
Old in young, transplanted
Young Sham
Young in old, transplanted
Young in young
Original age
Old
Young
D
Bilateral ovary transplantation
REACTOME Mitochondrial translation
REACTOME Protein ubiquitination
HALLMARK Oxidative phosphorylation
REACTOME Respiratory electron transport
HALLMARK Fatty acid metabolism
REACTOME Fatty acid metabolism
KEGG MAPK signaling pathway
REACTOME Inflammasomes
HALLMARK Il6 JAK STAT3 signaling
HALLMARK Inflammatory response
HALLMARK Complement
REACTOME Signaling by ILs
HALLMARK TNFa signaling via NF-kB
KEGG Apoptosis
HALLMARK Interferon gamma response
HALLMARK Interferon alpha response
HALLMARK Coagulation
KEGG Complement and coagulation cascades
KEGG Ribosome
REACTOME Selenoamino acid metabolism
HALLMARK Xenobiotic metabolism
KEGG Drug metabolism cytochrome P450
REACTOME Extracellular matrix organization
HALLMARK p53 pathway
HALLMARK Apoptosis
HALLMARK Hypoxia
HALLMARK mTORC1 signaling
NES
(C:Y in Y) Old in Young: Brain, Bilateral
(C:Y in Y) Young in Old: Ovary, Bilateral
Aging: Ovary
Mortality: Rodents
Aging: Liver
Aging: Kidney
Aging: Brain
Aging: Rodents
(C:Y in Y) Old in Young: Uterus, Bilateral
(C:O in O) Young in Old: Brain, Bilateral
Interventions: Max lifespan
(C:O in O) Young in Old: Uterus, Bilateral
(C:O in O) Old in Young: Ovary, Bilateral
Interventions: CR
Interventions: Rapamycin
Interventions: GH deficiency

**Extended Data Fig. 6 | Estimated cell-type proportions after unilateral transplantation**

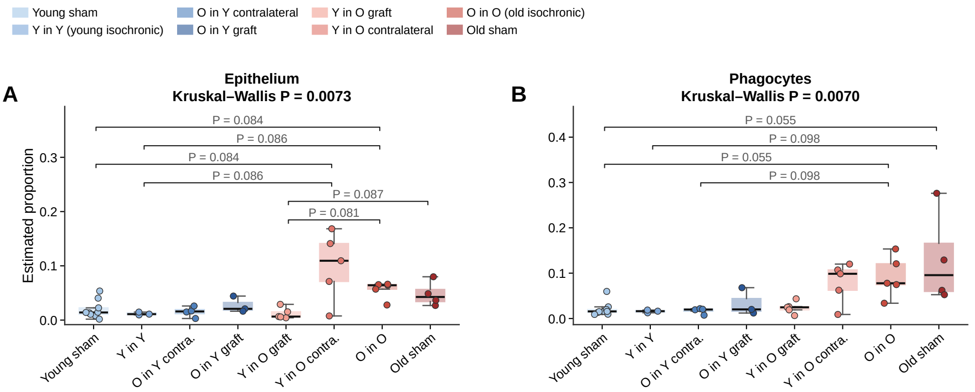


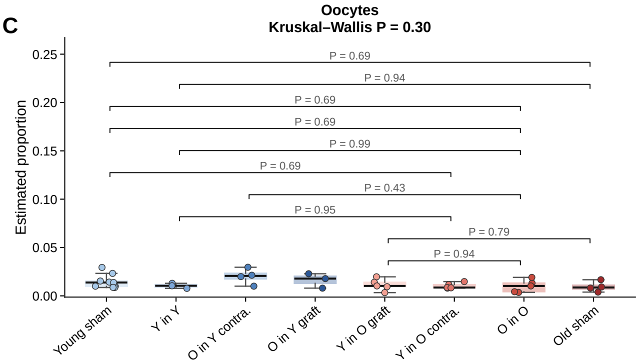


Brackets in A–C: the comparisons marked in the original analysis (A: 6 pairs, B: 4 pairs; C: both sets). P: two-sided Dunn test, BH-adjusted over all 28 pairs; bold, P < 0.05. The original plots used one-sided Dunn P values, which are half of these.

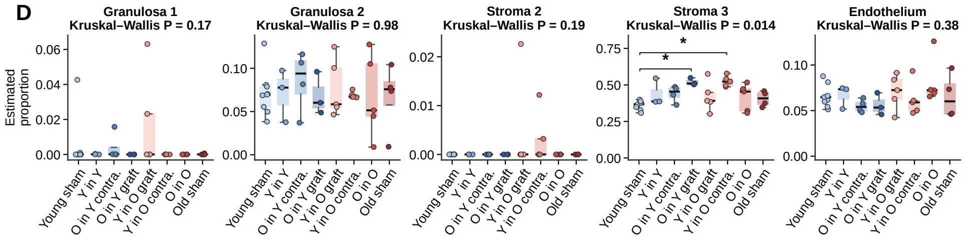


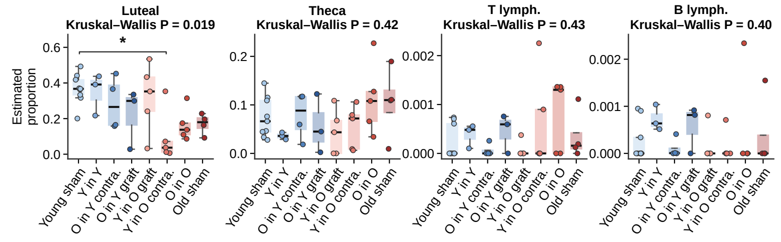


D: stars, two-sided Dunn BH P < 0.05 (* <0.05). Stroma 1 was 0 in all samples (not tested).

**Ext Fig. 7**

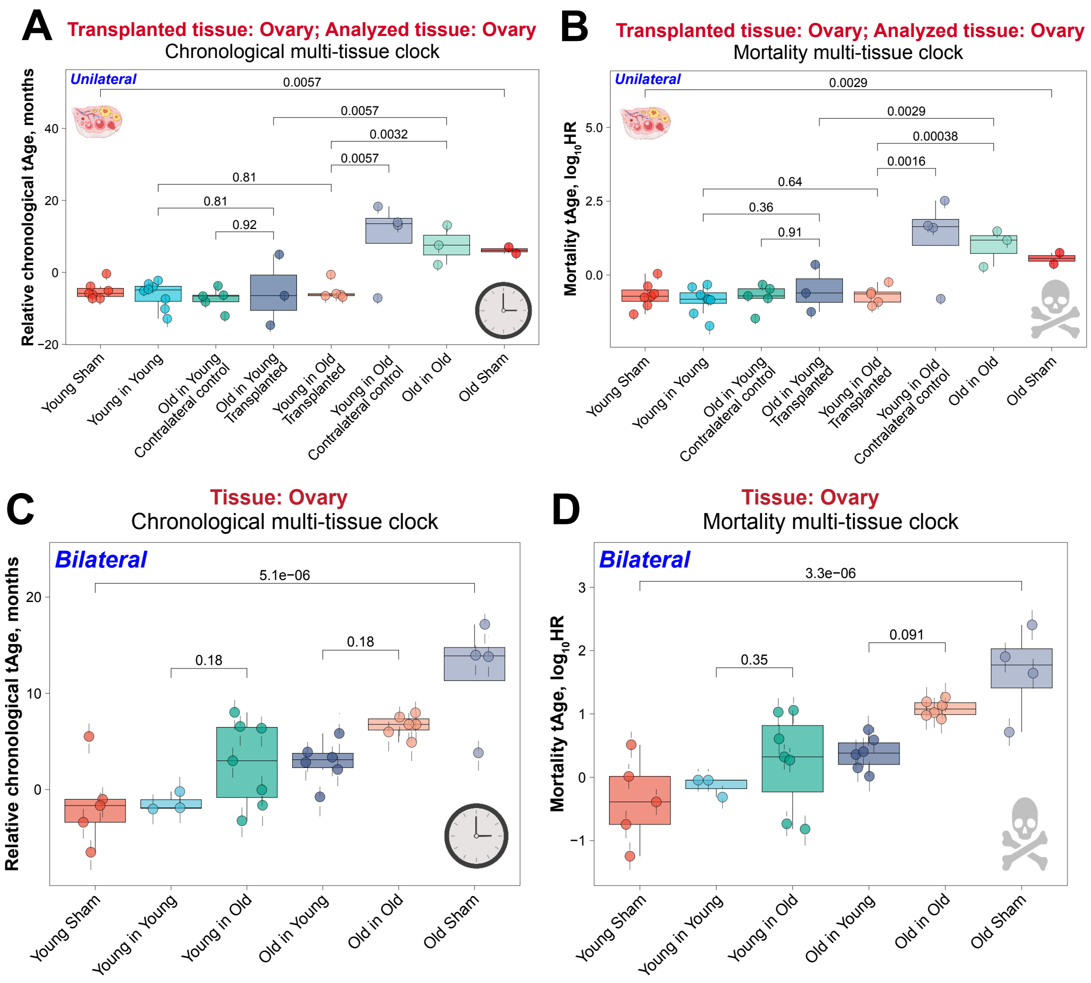
A
Transplanted tissue: Ovary; Analyzed tissue: Ovary
Chronological multi-tissue clock
Unilateral
Relative chronological tAge, months
0.0057
0.0057
0.0032
0.0057
0.81
0.81
0.92
Young Sham
Young in Young
Old in Young Contralateral control
Old in Young Transplanted
Young in Old Transplanted
Young in Old Contralateral control
Old in Old
Old Sham
B
Transplanted tissue: Ovary; Analyzed tissue: Ovary
Mortality multi-tissue clock
Unilateral
Mortality tAge, log10HR
0.0029
0.0029
0.00038
0.0016
0.64
0.36
0.91
C
Tissue: Ovary
Chronological multi-tissue clock
Bilateral
Relative chronological tAge, months
5.1e−06
0.18
0.18
Young Sham
Young in Young
Young in Old
Old in Young
Old in Old
Old Sham
D
Tissue: Ovary
Mortality multi-tissue clock
Bilateral
Mortality tAge, log10HR
3.3e−06
0.35
0.091

**Ext Fig. 8**

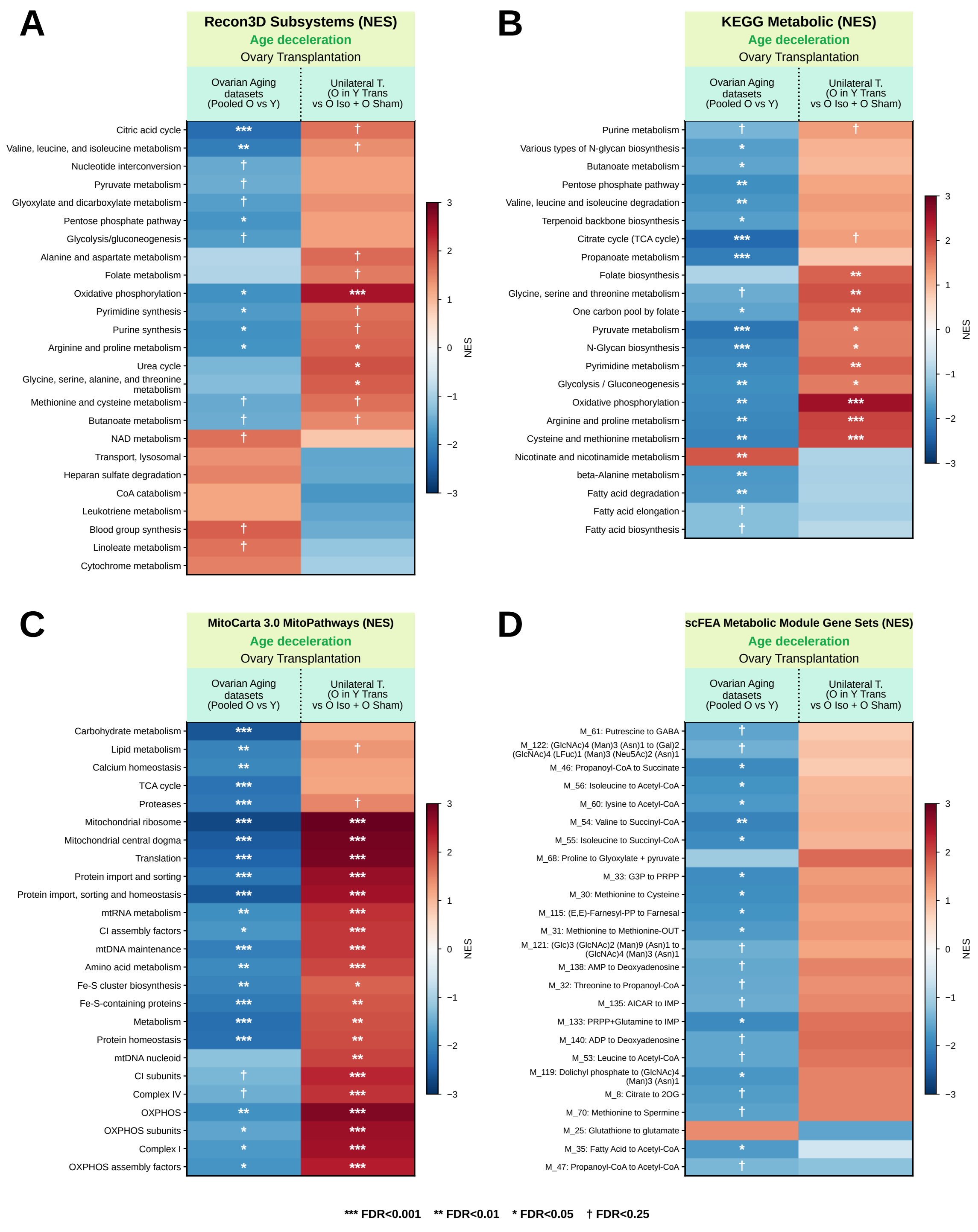
A
Recon3D Subsystems (NES)
Age deceleration
Ovary Transplantation
Ovarian Aging datasets (Pooled O vs Y)
Unilateral T. (O in Y Trans vs O Iso + O Sham)
Citric acid cycle
Valine, leucine, and isoleucine metabolism
Nucleotide interconversion
Pyruvate metabolism
Glyoxylate and dicarboxylate metabolism
Pentose phosphate pathway
Glycolysis/gluconeogenesis
Alanine and aspartate metabolism
Folate metabolism
Oxidative phosphorylation
Pyrimidine synthesis
Purine synthesis
Arginine and proline metabolism
Urea cycle
Glycine, serine, alanine, and threonine metabolism
Methionine and cysteine metabolism
Butanoate metabolism
NAD metabolism
Transport, lysosomal
Heparan sulfate degradation
CoA catabolism
Leukotriene metabolism
Blood group synthesis
Linoleate metabolism
Cytochrome metabolism
NES
B
KEGG Metabolic (NES)
Age deceleration
Ovary Transplantation
Ovarian Aging datasets (Pooled O vs Y)
Unilateral T. (O in Y Trans vs O Iso + O Sham)
Purine metabolism
Various types of N-glycan biosynthesis
Butanoate metabolism
Pentose phosphate pathway
Valine, leucine and isoleucine degradation
Terpenoid backbone biosynthesis
Citrate cycle (TCA cycle)
Propanoate metabolism
Folate biosynthesis
Glycine, serine and threonine metabolism
One carbon pool by folate
Pyruvate metabolism
N-Glycan biosynthesis
Pyrimidine metabolism
Glycolysis / Gluconeogenesis
Oxidative phosphorylation
Arginine and proline metabolism
Cysteine and methionine metabolism
Nicotinate and nicotinamide metabolism
beta-Alanine metabolism
Fatty acid degradation
Fatty acid elongation
Fatty acid biosynthesis
NES
C
MitoCarta 3.0 MitoPathways (NES)
Age deceleration
Ovary Transplantation
Ovarian Aging datasets (Pooled O vs Y)
Unilateral T. (O in Y Trans vs O Iso + O Sham)
Carbohydrate metabolism
Lipid metabolism
Calcium homeostasis
TCA cycle
Proteases
Mitochondrial ribosome
Mitochondrial central dogma
Translation
Protein import and sorting
Protein import, sorting and homeostasis
mtRNA metabolism
CI assembly factors
mtDNA maintenance
Amino acid metabolism
Fe-S cluster biosynthesis
Fe-S-containing proteins
Metabolism
Protein homeostasis
mtDNA nucleoid
CI subunits
Complex IV
OXPHOS
OXPHOS subunits
Complex I
OXPHOS assembly factors
NES
D
scFEA Metabolic Module Gene Sets (NES)
Age deceleration
Ovary Transplantation
Ovarian Aging datasets (Pooled O vs Y)
Unilateral T. (O in Y Trans vs O Iso + O Sham)
M_61: Putrescine to GABA
M_122: (GlcNAc)4 (Man)3 (Asn)1 to (Gal)2 (GlcNAc)4 (LFuc)1 (Man)3 (Neu5Ac)2 (Asn)1
M_46: Propanoyl-CoA to Succinate
M_56: Isoleucine to Acetyl-CoA
M_60: lysine to Acetyl-CoA
M_54: Valine to Succinyl-CoA
M_55: Isoleucine to Succinyl-CoA
M_68: Proline to Glyoxylate + pyruvate
M_33: G3P to PRPP
M_30: Methionine to Cysteine
M_115: (E,E)-Farnesyl-PP to Farnesal
M_31: Methionine to Methionine-OUT
M_121: (Glc)3 (GlcNAc)2 (Man)9 (Asn)1 to (GlcNAc)4 (Man)3 (Asn)1
M_138: AMP to Deoxyadenosine
M_32: Threonine to Propanoyl-CoA
M_135: AICAR to IMP
M_133: PRPP+Glutamine to IMP
M_140: ADP to Deoxyadenosine
M_53: Leucine to Acetyl-CoA
M_119: Dolichyl phosphate to (GlcNAc)4 (Man)3 (Asn)1
M_8: Citrate to 2OG
M_70: Methionine to Spermine
M_25: Glutathione to glutamate
M_35: Fatty Acid to Acetyl-CoA
M_47: Propanoyl-CoA to Acetyl-CoA
NES
*** FDR<0.001 ** FDR<0.01 * FDR<0.05 † FDR<0.25

Ext Fig. 9

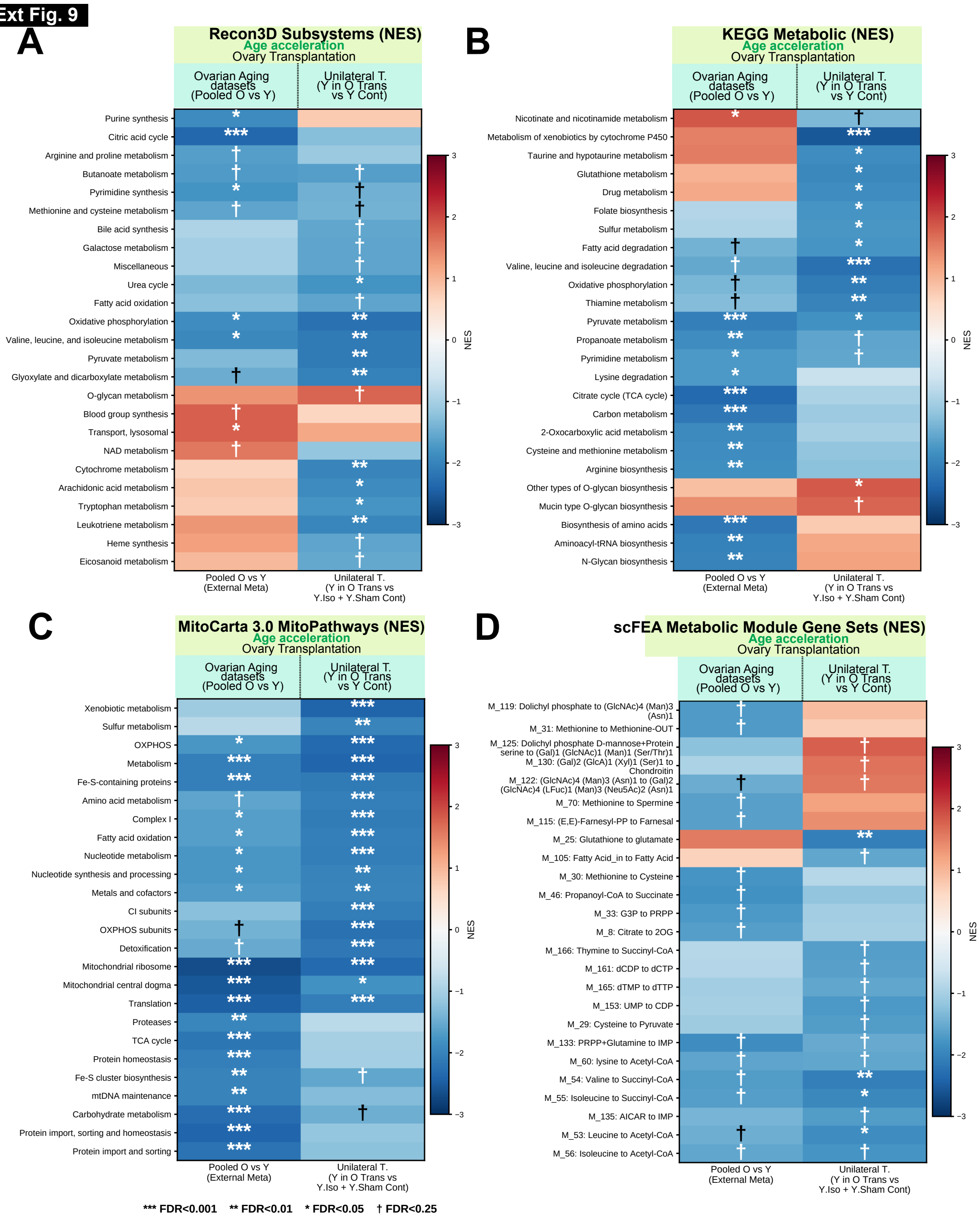
A
Recon3D Subsystems (NES)
Age acceleration
Ovary Transplantation
Ovarian Aging datasets (Pooled O vs Y)
Unilateral T. (Y in O Trans vs Y Cont)
Purine synthesis
Citric acid cycle
Arginine and proline metabolism
Butanoate metabolism
Pyrimidine synthesis
Methionine and cysteine metabolism
Bile acid synthesis
Galactose metabolism
Miscellaneous
Urea cycle
Fatty acid oxidation
Oxidative phosphorylation
Valine, leucine, and isoleucine metabolism
Pyruvate metabolism
Glyoxylate and dicarboxylate metabolism
O-glycan metabolism
Blood group synthesis
Transport, lysosomal
NAD metabolism
Cytochrome metabolism
Arachidonic acid metabolism
Tryptophan metabolism
Leukotriene metabolism
Heme synthesis
Eicosanoid metabolism
Pooled O vs Y (External Meta)
Unilateral T. (Y in O Trans vs Y.Iso + Y.Sham Cont)
NES
B
KEGG Metabolic (NES)
Age acceleration
Ovary Transplantation
Ovarian Aging datasets (Pooled O vs Y)
Unilateral T. (Y in O Trans vs Y Cont)
Nicotinate and nicotinamide metabolism
Metabolism of xenobiotics by cytochrome P450
Taurine and hypotaurine metabolism
Glutathione metabolism
Drug metabolism
Folate biosynthesis
Sulfur metabolism
Fatty acid degradation
Valine, leucine and isoleucine degradation
Oxidative phosphorylation
Thiamine metabolism
Pyruvate metabolism
Propanoate metabolism
Pyrimidine metabolism
Lysine degradation
Citrate cycle (TCA cycle)
Carbon metabolism
2-Oxocarboxylic acid metabolism
Cysteine and methionine metabolism
Arginine biosynthesis
Other types of O-glycan biosynthesis
Mucin type O-glycan biosynthesis
Biosynthesis of amino acids
Aminoacyl-tRNA biosynthesis
N-Glycan biosynthesis
Pooled O vs Y (External Meta)
Unilateral T. (Y in O Trans vs Y.Iso + Y.Sham Cont)
NES
C
MitoCarta 3.0 MitoPathways (NES)
Age acceleration
Ovary Transplantation
Ovarian Aging datasets (Pooled O vs Y)
Unilateral T. (Y in O Trans vs Y Cont)
Xenobiotic metabolism
Sulfur metabolism
OXPHOS
Metabolism
Fe-S-containing proteins
Amino acid metabolism
Complex I
Fatty acid oxidation
Nucleotide metabolism
Nucleotide synthesis and processing
Metals and cofactors
CI subunits
OXPHOS subunits
Detoxification
Mitochondrial ribosome
Mitochondrial central dogma
Translation
Proteases
TCA cycle
Protein homeostasis
Fe-S cluster biosynthesis
mtDNA maintenance
Carbohydrate metabolism
Protein import, sorting and homeostasis
Protein import and sorting
Pooled O vs Y (External Meta)
Unilateral T. (Y in O Trans vs Y.Iso + Y.Sham Cont)
NES
D
scFEA Metabolic Module Gene Sets (NES)
Age acceleration
Ovary Transplantation
Ovarian Aging datasets (Pooled O vs Y)
Unilateral T. (Y in O Trans vs Y Cont)
M_119: Dolichyl phosphate to (GlcNAc)4 (Man)3 (Asn)1
M_31: Methionine to Methionine-OUT
M_125: Dolichyl phosphate D-mannose+Protein serine to (Gal)1 (GlcNAc)1 (Man)1 (Ser/Thr)1
M_130: (Gal)2 (GlcA)1 (Xyl)1 (Ser)1 to Chondroitin
M_122: (GlcNAc)4 (Man)3 (Asn)1 to (Gal)2 (GlcNAc)4 (LFuc)1 (Man)3 (Neu5Ac)2 (Asn)1
M_70: Methionine to Spermine
M_115: (E,E)-Farnesyl-PP to Farnesal
M_25: Glutathione to glutamate
M_105: Fatty Acid_in to Fatty Acid
M_30: Methionine to Cysteine
M_46: Propanoyl-CoA to Succinate
M_33: G3P to PRPP
M_8: Citrate to 2OG
M_166: Thymine to Succinyl-CoA
M_161: dCDP to dCTP
M_165: dTMP to dTTP
M_153: UMP to CDP
M_29: Cysteine to Pyruvate
M_133: PRPP+Glutamine to IMP
M_60: lysine to Acetyl-CoA
M_54: Valine to Succinyl-CoA
M_55: Isoleucine to Succinyl-CoA
M_135: AICAR to IMP
M_53: Leucine to Acetyl-CoA
M_56: Isoleucine to Acetyl-CoA
Pooled O vs Y (External Meta)
Unilateral T. (Y in O Trans vs Y.Iso + Y.Sham Cont)
NES
*** FDR<0.001 ** FDR<0.01 * FDR<0.05 † FDR<0.25

**Ext Fig. 10**

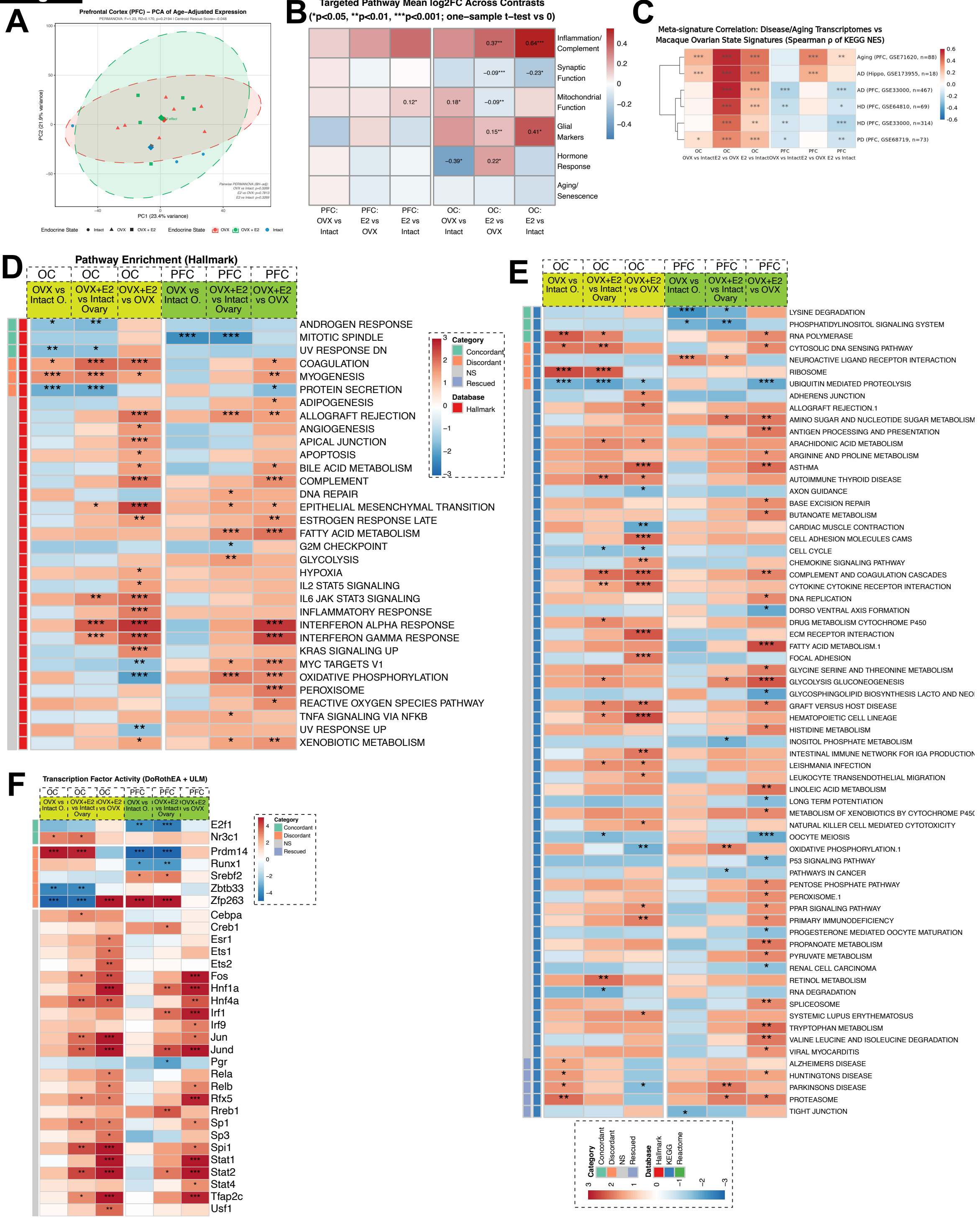
A
Prefrontal Cortex (PFC) – PCA of Age–Adjusted Expression
PC1 (23.4% variance)
PC2 (21.9% variance)
Endocrine State
Intact
OVX
OVX + E2
B
Targeted Pathway Mean log2FC Across Contrasts
(*p<0.05, **p<0.01, ***p<0.001; one–sample t–test vs 0)
Inflammation/ Complement
Synaptic Function
Mitochondrial Function
Glial Markers
Hormone Response
Aging/ Senescence
PFC: OVX vs Intact
PFC: E2 vs OVX
PFC: E2 vs Intact
OC: OVX vs Intact
OC: E2 vs OVX
OC: E2 vs Intact
0.37**
0.64***
−0.09***
−0.23*
0.12*
0.18*
−0.09**
0.15**
0.41*
−0.39*
0.22*
C
Meta-signature Correlation: Disease/Aging Transcriptomes vs Macaque Ovarian State Signatures (Spearman ρ of KEGG NES)
Aging (PFC, GSE71620, n=88)
AD (Hippo, GSE173955, n=18)
AD (PFC, GSE33000, n=467)
HD (PFC, GSE64810, n=69)
HD (PFC, GSE33000, n=314)
PD (PFC, GSE68719, n=73)
D
Pathway Enrichment (Hallmark)
OC
PFC
OVX vs Intact O.
OVX+E2 vs Intact Ovary
OVX+E2 vs OVX
Category
Concordant
Discordant
NS
Rescued
Database
Hallmark
ANDROGEN RESPONSE
MITOTIC SPINDLE
UV RESPONSE DN
COAGULATION
MYOGENESIS
PROTEIN SECRETION
ADIPOGENESIS
ALLOGRAFT REJECTION
ANGIOGENESIS
APICAL JUNCTION
APOPTOSIS
BILE ACID METABOLISM
COMPLEMENT
DNA REPAIR
EPITHELIAL MESENCHYMAL TRANSITION
ESTROGEN RESPONSE LATE
FATTY ACID METABOLISM
G2M CHECKPOINT
GLYCOLYSIS
HYPOXIA
IL2 STAT5 SIGNALING
IL6 JAK STAT3 SIGNALING
INFLAMMATORY RESPONSE
INTERFERON ALPHA RESPONSE
INTERFERON GAMMA RESPONSE
KRAS SIGNALING UP
MYC TARGETS V1
OXIDATIVE PHOSPHORYLATION
PEROXISOME
REACTIVE OXYGEN SPECIES PATHWAY
TNFA SIGNALING VIA NFKB
UV RESPONSE UP
XENOBIOTIC METABOLISM
E
LYSINE DEGRADATION
PHOSPHATIDYLINOSITOL SIGNALING SYSTEM
RNA POLYMERASE
CYTOSOLIC DNA SENSING PATHWAY
NEUROACTIVE LIGAND RECEPTOR INTERACTION
RIBOSOME
UBIQUITIN MEDIATED PROTEOLYSIS
ADHERENS JUNCTION
ALLOGRAFT REJECTION.1
AMINO SUGAR AND NUCLEOTIDE SUGAR METABOLISM
ANTIGEN PROCESSING AND PRESENTATION
ARACHIDONIC ACID METABOLISM
ARGININE AND PROLINE METABOLISM
ASTHMA
AUTOIMMUNE THYROID DISEASE
AXON GUIDANCE
BASE EXCISION REPAIR
BUTANOATE METABOLISM
CARDIAC MUSCLE CONTRACTION
CELL ADHESION MOLECULES CAMS
CELL CYCLE
CHEMOKINE SIGNALING PATHWAY
COMPLEMENT AND COAGULATION CASCADES
CYTOKINE CYTOKINE RECEPTOR INTERACTION
DNA REPLICATION
DORSO VENTRAL AXIS FORMATION
DRUG METABOLISM CYTOCHROME P450
ECM RECEPTOR INTERACTION
FATTY ACID METABOLISM.1
FOCAL ADHESION
GLYCINE SERINE AND THREONINE METABOLISM
GLYCOLYSIS GLUCONEOGENESIS
GLYCOSPHINGOLIPID BIOSYNTHESIS LACTO AND NEO
GRAFT VERSUS HOST DISEASE
HEMATOPOIETIC CELL LINEAGE
HISTIDINE METABOLISM
INOSITOL PHOSPHATE METABOLISM
INTESTINAL IMMUNE NETWORK FOR IGA PRODUCTION
LEISHMANIA INFECTION
LEUKOCYTE TRANSENDOTHELIAL MIGRATION
LINOLEIC ACID METABOLISM
LONG TERM POTENTIATION
METABOLISM OF XENOBIOTICS BY CYTOCHROME P450
NATURAL KILLER CELL MEDIATED CYTOTOXICITY
OOCYTE MEIOSIS
OXIDATIVE PHOSPHORYLATION.1
P53 SIGNALING PATHWAY
PATHWAYS IN CANCER
PENTOSE PHOSPHATE PATHWAY
PEROXISOME.1
PPAR SIGNALING PATHWAY
PRIMARY IMMUNODEFICIENCY
PROGESTERONE MEDIATED OOCYTE MATURATION
PROPANOATE METABOLISM
PYRUVATE METABOLISM
RENAL CELL CARCINOMA
RETINOL METABOLISM
RNA DEGRADATION
SPLICEOSOME
SYSTEMIC LUPUS ERYTHEMATOSUS
TRYPTOPHAN METABOLISM
VALINE LEUCINE AND ISOLEUCINE DEGRADATION
VIRAL MYOCARDITIS
ALZHEIMERS DISEASE
HUNTINGTONS DISEASE
PARKINSONS DISEASE
PROTEASOME
TIGHT JUNCTION
Category
Concordant
Discordant
NS
Rescued
Database
Hallmark
KEGG
Reactome
F
Transcription Factor Activity (DoRothEA + ULM)
E2f1
Nr3c1
Prdm14
Runx1
Srebf2
Zbtb33
Zfp263
Cebpa
Creb1
Esr1
Ets1
Ets2
Fos
Hnf1a
Hnf4a
Irf1
Irf9
Jun
Jund
Pgr
Rela
Relb
Rfx5
Rreb1
Sp1
Sp3
Spi1
Stat1
Stat2
Stat4
Tfap2c
Usf1

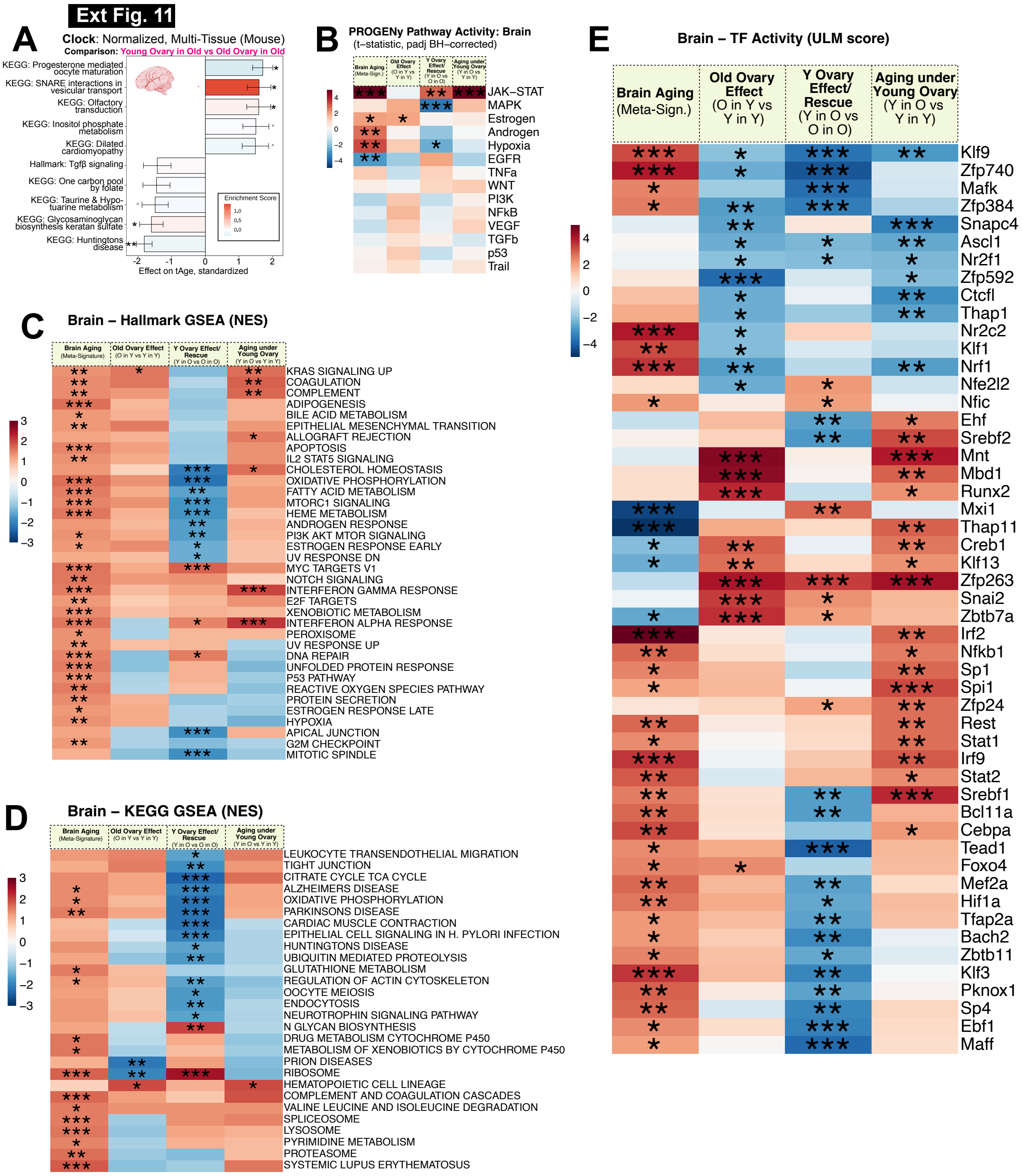
Ext Fig. 11
A
Clock: Normalized, Multi-Tissue (Mouse)
Comparison: Young Ovary in Old vs Old Ovary in Old
KEGG: Progesterone mediated oocyte maturation
KEGG: SNARE interactions in vesicular transport
KEGG: Olfactory transduction
KEGG: Inositol phosphate metabolism
KEGG: Dilated cardiomyopathy
Hallmark: Tgfβ signaling
KEGG: One carbon pool by folate
KEGG: Taurine & Hypotaurine metabolism
KEGG: Glycosaminoglycan biosynthesis keratan sulfate
KEGG: Huntingtons disease
Enrichment Score
Effect on tAge, standardized
B
PROGENy Pathway Activity: Brain
(t–statistic, padj BH–corrected)
Brain Aging (Meta-Sign.)
Old Ovary Effect (O in Y vs Y in Y)
Y Ovary Effect/ Rescue (Y in O vs O in O)
Aging under Young Ovary (Y in O vs Y in Y)
JAK–STAT
MAPK
Estrogen
Androgen
Hypoxia
EGFR
TNFa
WNT
PI3K
NFkB
VEGF
TGFb
p53
Trail
C
Brain – Hallmark GSEA (NES)
Brain Aging (Meta-Signature)
Old Ovary Effect (O in Y vs Y in Y)
Y Ovary Effect/ Rescue (Y in O vs O in O)
Aging under Young Ovary (Y in O vs Y in Y)
KRAS SIGNALING UP
COAGULATION
COMPLEMENT
ADIPOGENESIS
BILE ACID METABOLISM
EPITHELIAL MESENCHYMAL TRANSITION
ALLOGRAFT REJECTION
APOPTOSIS
IL2 STAT5 SIGNALING
CHOLESTEROL HOMEOSTASIS
OXIDATIVE PHOSPHORYLATION
FATTY ACID METABOLISM
MTORC1 SIGNALING
HEME METABOLISM
ANDROGEN RESPONSE
PI3K AKT MTOR SIGNALING
ESTROGEN RESPONSE EARLY
UV RESPONSE DN
MYC TARGETS V1
NOTCH SIGNALING
INTERFERON GAMMA RESPONSE
E2F TARGETS
XENOBIOTIC METABOLISM
INTERFERON ALPHA RESPONSE
PEROXISOME
UV RESPONSE UP
DNA REPAIR
UNFOLDED PROTEIN RESPONSE
P53 PATHWAY
REACTIVE OXYGEN SPECIES PATHWAY
PROTEIN SECRETION
ESTROGEN RESPONSE LATE
HYPOXIA
APICAL JUNCTION
G2M CHECKPOINT
MITOTIC SPINDLE
D
Brain – KEGG GSEA (NES)
LEUKOCYTE TRANSENDOTHELIAL MIGRATION
TIGHT JUNCTION
CITRATE CYCLE TCA CYCLE
ALZHEIMERS DISEASE
OXIDATIVE PHOSPHORYLATION
PARKINSONS DISEASE
CARDIAC MUSCLE CONTRACTION
EPITHELIAL CELL SIGNALING IN H. PYLORI INFECTION
HUNTINGTONS DISEASE
UBIQUITIN MEDIATED PROTEOLYSIS
GLUTATHIONE METABOLISM
REGULATION OF ACTIN CYTOSKELETON
OOCYTE MEIOSIS
ENDOCYTOSIS
NEUROTROPHIN SIGNALING PATHWAY
N GLYCAN BIOSYNTHESIS
DRUG METABOLISM CYTOCHROME P450
METABOLISM OF XENOBIOTICS BY CYTOCHROME P450
PRION DISEASES
RIBOSOME
HEMATOPOIETIC CELL LINEAGE
COMPLEMENT AND COAGULATION CASCADES
VALINE LEUCINE AND ISOLEUCINE DEGRADATION
SPLICEOSOME
LYSOSOME
PYRIMIDINE METABOLISM
PROTEASOME
SYSTEMIC LUPUS ERYTHEMATOSUS
E
Brain – TF Activity (ULM score)
Brain Aging (Meta-Sign.)
Old Ovary Effect (O in Y vs Y in Y)
Y Ovary Effect/ Rescue (Y in O vs O in O)
Aging under Young Ovary (Y in O vs Y in Y)
Klf9
Zfp740
Mafk
Zfp384
Snapc4
Ascl1
Nr2f1
Zfp592
Ctcfl
Thap1
Nr2c2
Klf1
Nrf1
Nfe2l2
Nfic
Ehf
Srebf2
Mnt
Mbd1
Runx2
Mxi1
Thap11
Creb1
Klf13
Zfp263
Snai2
Zbtb7a
Irf2
Nfkb1
Sp1
Spi1
Zfp24
Rest
Stat1
Irf9
Stat2
Srebf1
Bcl11a
Cebpa
Tead1
Foxo4
Mef2a
Hif1a
Tfap2a
Bach2
Zbtb11
Klf3
Pknox1
Sp4
Ebf1
Maff

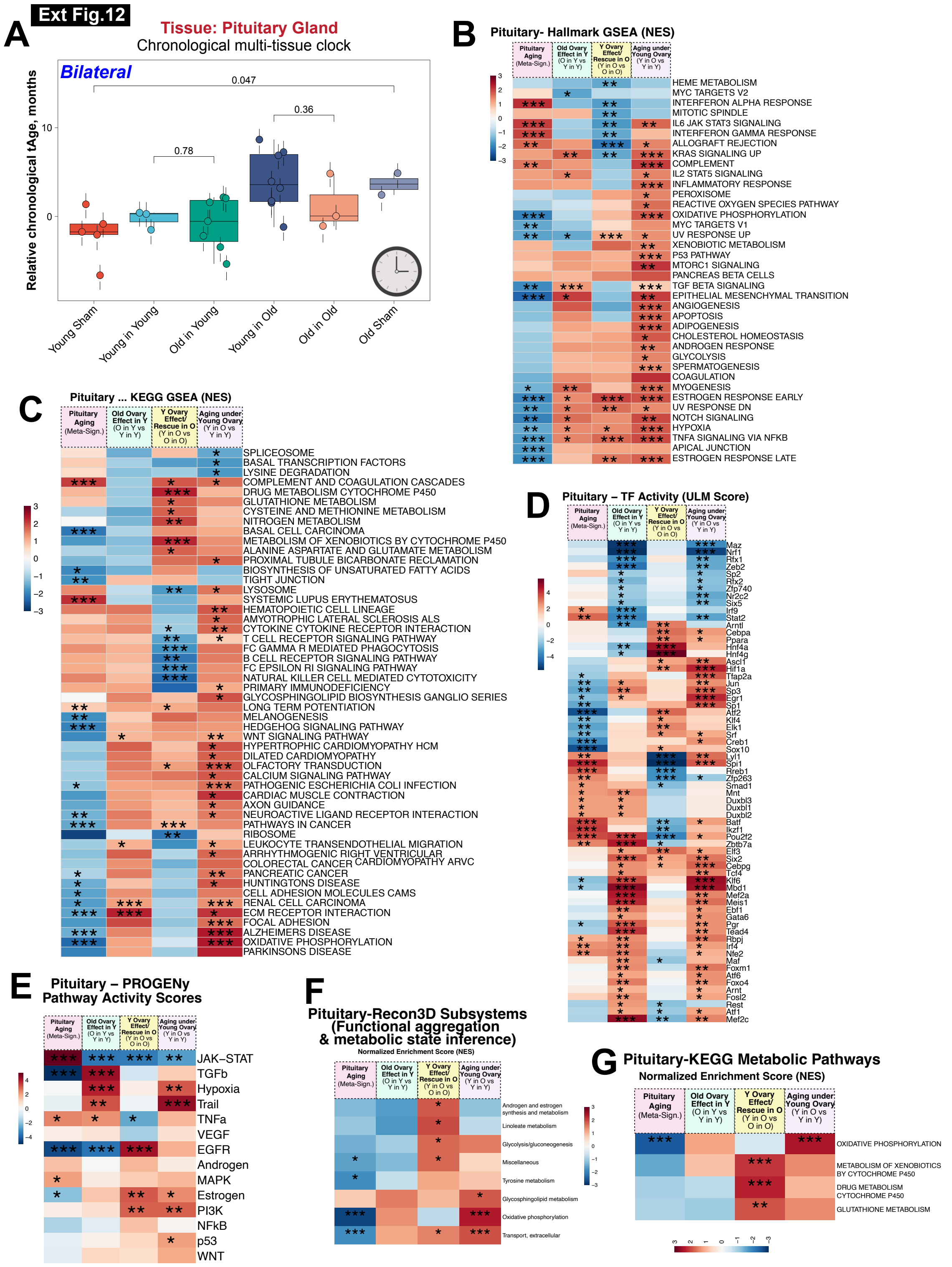

Ext Fig.12
A
Tissue: Pituitary Gland
Chronological multi-tissue clock
Bilateral
Relative chronological tAge, months
Young Sham
Young in Young
Old in Young
Young in Old
Old in Old
Old Sham
B
Pituitary- Hallmark GSEA (NES)
C
Pituitary ... KEGG GSEA (NES)
D
Pituitary – TF Activity (ULM Score)
E
Pituitary – PROGENy Pathway Activity Scores
F
Pituitary-Recon3D Subsystems (Functional aggregation & metabolic state inference)
Normalized Enrichment Score (NES)
G
Pituitary-KEGG Metabolic Pathways
Normalized Enrichment Score (NES)

### Ext Fig.13

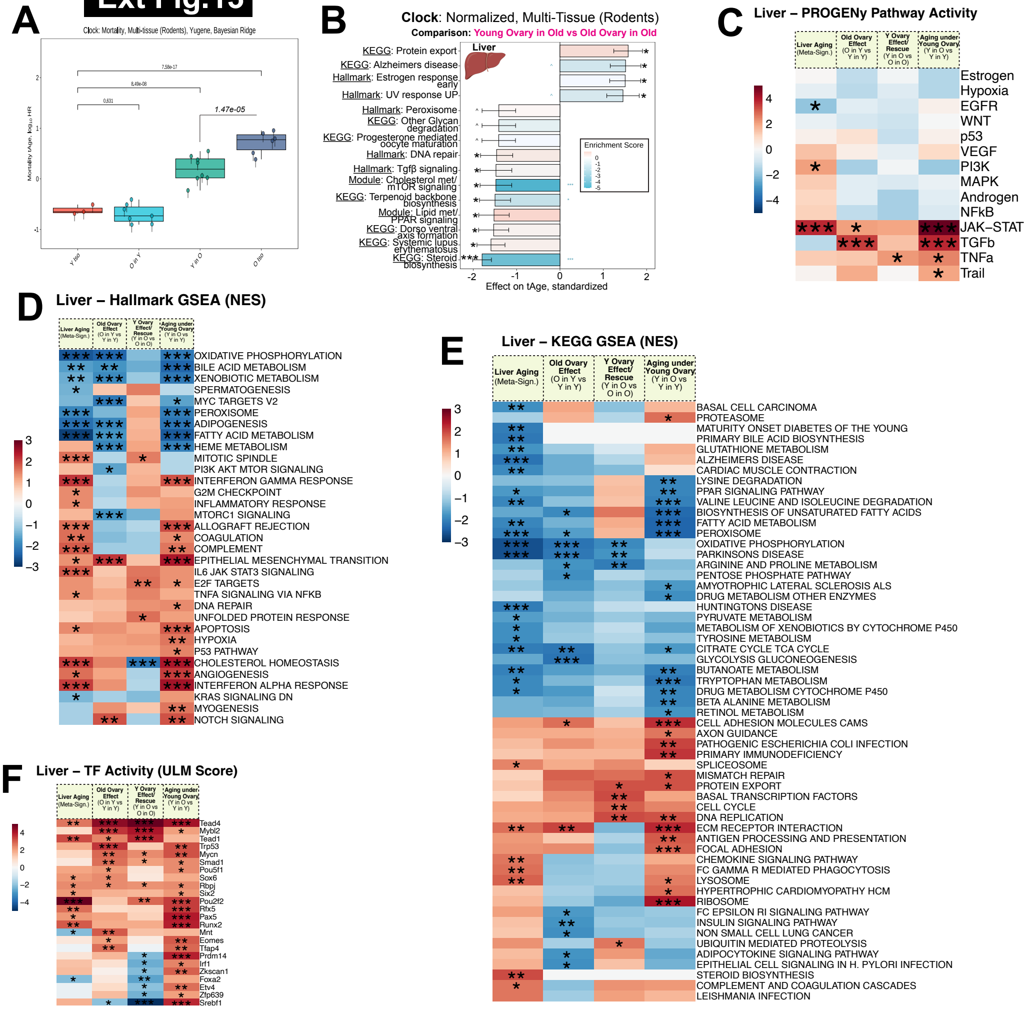
A
Clock: Mortality, Multi-tissue (Rodents), Yugene, Bayesian Ridge
7.58e-17
8.49e-08
0.831
1.47e-05
Mortality tAge, log10 HR
Y in Y
O in Y
Y in O
O in O
B
Clock: Normalized, Multi-Tissue (Rodents)
Comparison: Young Ovary in Old vs Old Ovary in Old
Liver
KEGG: Protein export
KEGG: Alzheimers disease
Hallmark: Estrogen response early
Hallmark: UV response UP
Hallmark: Peroxisome
KEGG: Other Glycan degradation
KEGG: Progesterone mediated oocyte maturation
Hallmark: DNA repair
Hallmark: Tgfβ signaling
Module: Cholesterol met/ mTOR signaling
KEGG: Terpenoid backbone biosynthesis
Module: Lipid met/ PPAR signaling
KEGG: Dorso ventral axis formation
KEGG: Systemic lupus erythematosus
KEGG: Steroid biosynthesis
Enrichment Score
Effect on tAge, standardized
C
Liver – PROGENy Pathway Activity
Liver Aging (Meta-Sign.)
Old Ovary Effect (O in Y vs Y in Y)
Y Ovary Effect/ Rescue (Y in O vs O in O)
Aging under Young Ovary (Y in O vs Y in Y)
Estrogen
Hypoxia
EGFR
WNT
p53
VEGF
PI3K
MAPK
Androgen
NFkB
JAK–STAT
TGFb
TNFa
Trail
D
Liver – Hallmark GSEA (NES)
OXIDATIVE PHOSPHORYLATION
BILE ACID METABOLISM
XENOBIOTIC METABOLISM
SPERMATOGENESIS
MYC TARGETS V2
PEROXISOME
ADIPOGENESIS
FATTY ACID METABOLISM
HEME METABOLISM
MITOTIC SPINDLE
PI3K AKT MTOR SIGNALING
INTERFERON GAMMA RESPONSE
G2M CHECKPOINT
INFLAMMATORY RESPONSE
MTORC1 SIGNALING
ALLOGRAFT REJECTION
COAGULATION
COMPLEMENT
EPITHELIAL MESENCHYMAL TRANSITION
IL6 JAK STAT3 SIGNALING
E2F TARGETS
TNFA SIGNALING VIA NFKB
DNA REPAIR
UNFOLDED PROTEIN RESPONSE
APOPTOSIS
HYPOXIA
P53 PATHWAY
CHOLESTEROL HOMEOSTASIS
ANGIOGENESIS
INTERFERON ALPHA RESPONSE
KRAS SIGNALING DN
MYOGENESIS
NOTCH SIGNALING
E
Liver – KEGG GSEA (NES)
BASAL CELL CARCINOMA
PROTEASOME
MATURITY ONSET DIABETES OF THE YOUNG
PRIMARY BILE ACID BIOSYNTHESIS
GLUTATHIONE METABOLISM
ALZHEIMERS DISEASE
CARDIAC MUSCLE CONTRACTION
LYSINE DEGRADATION
PPAR SIGNALING PATHWAY
VALINE LEUCINE AND ISOLEUCINE DEGRADATION
BIOSYNTHESIS OF UNSATURATED FATTY ACIDS
FATTY ACID METABOLISM
PEROXISOME
OXIDATIVE PHOSPHORYLATION
PARKINSONS DISEASE
ARGININE AND PROLINE METABOLISM
PENTOSE PHOSPHATE PATHWAY
AMYOTROPHIC LATERAL SCLEROSIS ALS
DRUG METABOLISM OTHER ENZYMES
HUNTINGTONS DISEASE
PYRUVATE METABOLISM
METABOLISM OF XENOBIOTICS BY CYTOCHROME P450
TYROSINE METABOLISM
CITRATE CYCLE TCA CYCLE
GLYCOLYSIS GLUCONEOGENESIS
BUTANOATE METABOLISM
TRYPTOPHAN METABOLISM
DRUG METABOLISM CYTOCHROME P450
BETA ALANINE METABOLISM
RETINOL METABOLISM
CELL ADHESION MOLECULES CAMS
AXON GUIDANCE
PATHOGENIC ESCHERICHIA COLI INFECTION
PRIMARY IMMUNODEFICIENCY
SPLICEOSOME
MISMATCH REPAIR
PROTEIN EXPORT
BASAL TRANSCRIPTION FACTORS
CELL CYCLE
DNA REPLICATION
ECM RECEPTOR INTERACTION
ANTIGEN PROCESSING AND PRESENTATION
FOCAL ADHESION
CHEMOKINE SIGNALING PATHWAY
FC GAMMA R MEDIATED PHAGOCYTOSIS
LYSOSOME
HYPERTROPHIC CARDIOMYOPATHY HCM
RIBOSOME
FC EPSILON RI SIGNALING PATHWAY
INSULIN SIGNALING PATHWAY
NON SMALL CELL LUNG CANCER
UBIQUITIN MEDIATED PROTEOLYSIS
ADIPOCYTOKINE SIGNALING PATHWAY
EPITHELIAL CELL SIGNALING IN H. PYLORI INFECTION
STEROID BIOSYNTHESIS
COMPLEMENT AND COAGULATION CASCADES
LEISHMANIA INFECTION
F
Liver – TF Activity (ULM Score)
Tead4
Mybl2
Tead1
Trp53
Mycn
Smad1
Pou5f1
Sox6
Rbpj
Six2
Pou2f2
Rfx5
Pax5
Runx2
Mnt
Eomes
Tfap4
Prdm14
Irf1
Zkscan1
Foxa2
Etv4
Zfp639
Srebf1

### Ext Fig. 14

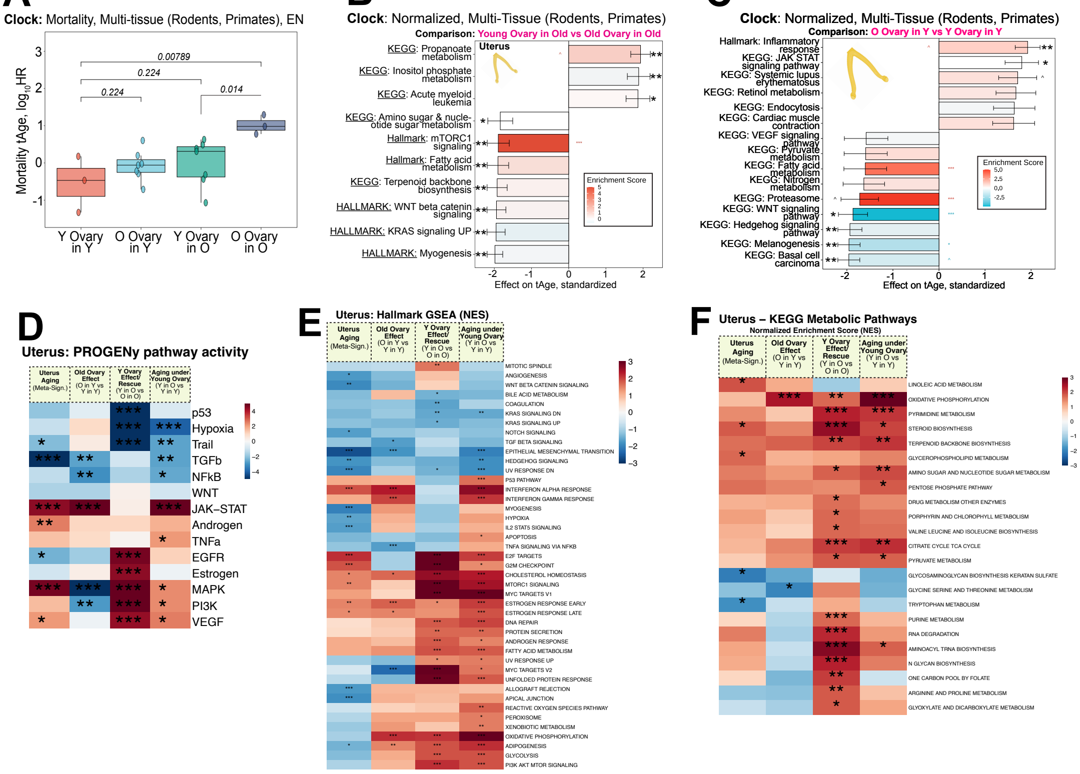
A
Clock: Mortality, Multi-tissue (Rodents, Primates), EN
Mortality tAge, log10HR
0.00789
0.224
0.224
0.014
Y Ovary in Y
O Ovary in Y
Y Ovary in O
O Ovary in O
B
Clock: Normalized, Multi-Tissue (Rodents, Primates)
Comparison: Young Ovary in Old vs Old Ovary in Old
Uterus
KEGG: Propanoate metabolism
KEGG: Inositol phosphate metabolism
KEGG: Acute myeloid leukemia
KEGG: Amino sugar & nucleotide sugar metabolism
Hallmark: mTORC1 signaling
Hallmark: Fatty acid metabolism
KEGG: Terpenoid backbone biosynthesis
HALLMARK: WNT beta catenin signaling
HALLMARK: KRAS signaling UP
HALLMARK: Myogenesis
Enrichment Score
Effect on tAge, standardized
C
Clock: Normalized, Multi-Tissue (Rodents, Primates)
Comparison: O Ovary in Y vs Y Ovary in Y
Hallmark: Inflammatory response
KEGG: JAK STAT signaling pathway
KEGG: Systemic lupus erythematosus
KEGG: Retinol metabolism
KEGG: Endocytosis
KEGG: Cardiac muscle contraction
KEGG: VEGF signaling pathway
KEGG: Pyruvate metabolism
KEGG: Fatty acid metabolism
KEGG: Nitrogen metabolism
KEGG: Proteasome
KEGG: WNT signaling pathway
KEGG: Hedgehog signaling pathway
KEGG: Melanogenesis
KEGG: Basal cell carcinoma
Enrichment Score
Effect on tAge, standardized
D
Uterus: PROGENy pathway activity
Uterus Aging (Meta-Sign.)
Old Ovary Effect (O in Y vs Y in Y)
Y Ovary Effect/Rescue (Y in O vs O in O)
Aging under Young Ovary (Y in O vs Y in Y)
p53
Hypoxia
Trail
TGFb
NFkB
WNT
JAK–STAT
Androgen
TNFa
EGFR
Estrogen
MAPK
PI3K
VEGF
E
Uterus: Hallmark GSEA (NES)
MITOTIC SPINDLE
ANGIOGENESIS
WNT BETA CATENIN SIGNALING
BILE ACID METABOLISM
COAGULATION
KRAS SIGNALING DN
KRAS SIGNALING UP
NOTCH SIGNALING
TGF BETA SIGNALING
EPITHELIAL MESENCHYMAL TRANSITION
HEDGEHOG SIGNALING
UV RESPONSE DN
P53 PATHWAY
INTERFERON ALPHA RESPONSE
INTERFERON GAMMA RESPONSE
MYOGENESIS
HYPOXIA
IL2 STAT5 SIGNALING
APOPTOSIS
TNFA SIGNALING VIA NFKB
E2F TARGETS
G2M CHECKPOINT
CHOLESTEROL HOMEOSTASIS
MTORC1 SIGNALING
MYC TARGETS V1
ESTROGEN RESPONSE EARLY
ESTROGEN RESPONSE LATE
DNA REPAIR
PROTEIN SECRETION
ANDROGEN RESPONSE
FATTY ACID METABOLISM
UV RESPONSE UP
MYC TARGETS V2
UNFOLDED PROTEIN RESPONSE
ALLOGRAFT REJECTION
APICAL JUNCTION
REACTIVE OXYGEN SPECIES PATHWAY
PEROXISOME
XENOBIOTIC METABOLISM
OXIDATIVE PHOSPHORYLATION
ADIPOGENESIS
GLYCOLYSIS
PI3K AKT MTOR SIGNALING
F
Uterus – KEGG Metabolic Pathways
Normalized Enrichment Score (NES)
LINOLEIC ACID METABOLISM
OXIDATIVE PHOSPHORYLATION
PYRIMIDINE METABOLISM
STEROID BIOSYNTHESIS
TERPENOID BACKBONE BIOSYNTHESIS
GLYCEROPHOSPHOLIPID METABOLISM
AMINO SUGAR AND NUCLEOTIDE SUGAR METABOLISM
PENTOSE PHOSPHATE PATHWAY
DRUG METABOLISM OTHER ENZYMES
PORPHYRIN AND CHLOROPHYLL METABOLISM
VALINE LEUCINE AND ISOLEUCINE BIOSYNTHESIS
CITRATE CYCLE TCA CYCLE
PYRUVATE METABOLISM
GLYCOSAMINOGLYCAN BIOSYNTHESIS KERATAN SULFATE
GLYCINE SERINE AND THREONINE METABOLISM
TRYPTOPHAN METABOLISM
PURINE METABOLISM
RNA DEGRADATION
AMINOACYL TRNA BIOSYNTHESIS
N GLYCAN BIOSYNTHESIS
ONE CARBON POOL BY FOLATE
ARGININE AND PROLINE METABOLISM
GLYOXYLATE AND DICARBOXYLATE METABOLISM

**Ext. Fig. 15**

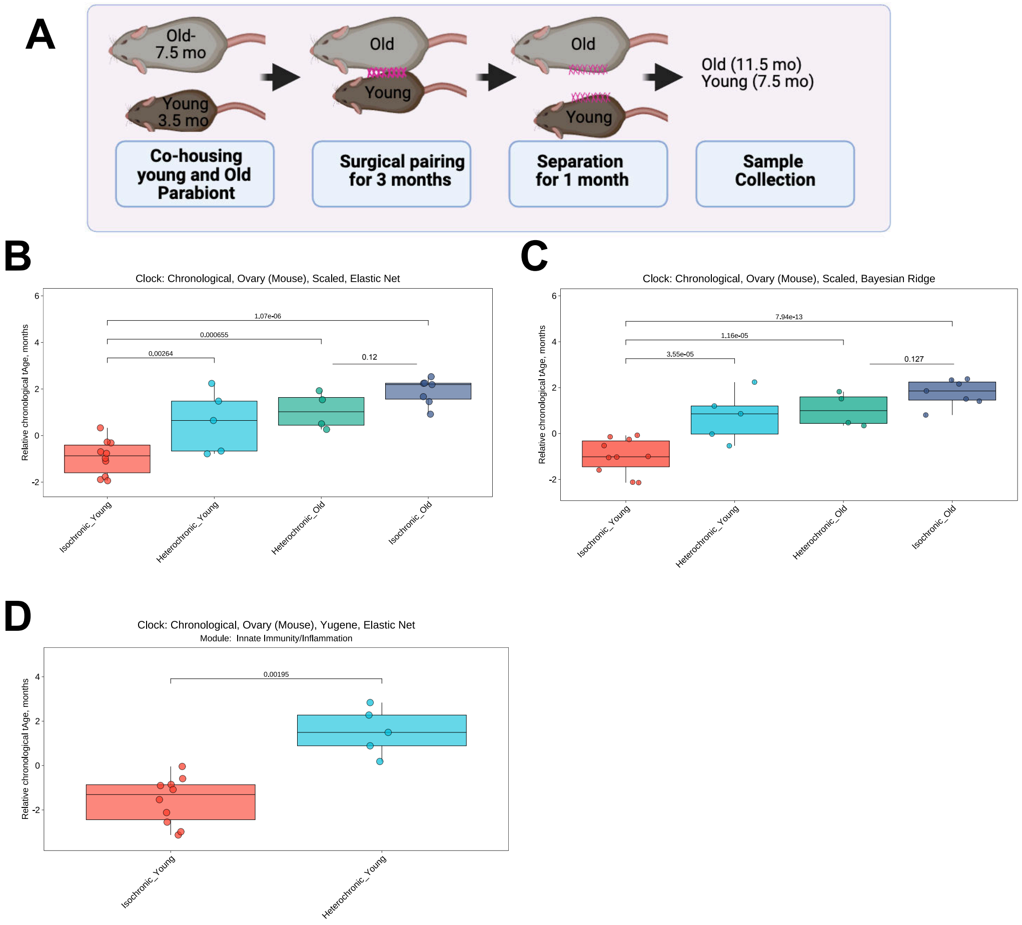
A
Old-
7.5 mo
Young
3.5 mo
Co-housing young and Old Parabiont
Old
Young
Surgical pairing for 3 months
Old
Young
Separation for 1 month
Old (11.5 mo)
Young (7.5 mo)
Sample Collection
B
Clock: Chronological, Ovary (Mouse), Scaled, Elastic Net
Relative chronological tAge, months
1.07e-06
0.000655
0.00264
0.12
Isochronic_Young
Heterochronic_Young
Heterochronic_Old
Isochronic_Old
C
Clock: Chronological, Ovary (Mouse), Scaled, Bayesian Ridge
Relative chronological tAge, months
7.94e-13
1.16e-05
3.55e-05
0.127
Isochronic_Young
Heterochronic_Young
Heterochronic_Old
Isochronic_Old
D
Clock: Chronological, Ovary (Mouse), Yugene, Elastic Net
Module: Innate Immunity/Inflammation
Relative chronological tAge, months
0.00195
Isochronic_Young
Heterochronic_Young

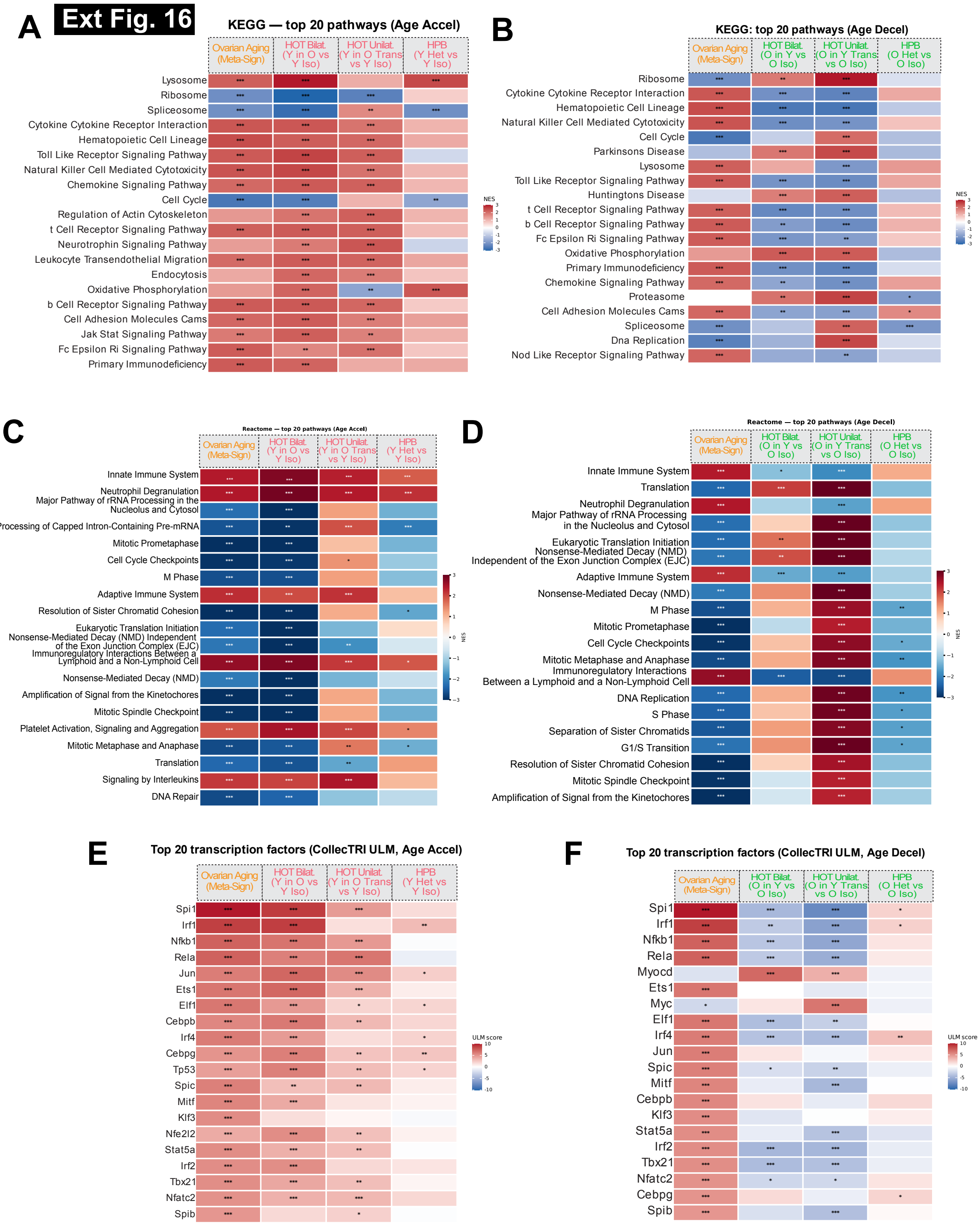

Ext Fig. 16
A
KEGG — top 20 pathways (Age Accel)
Ovarian Aging (Meta-Sign)
HOT Bilat. (Y in O vs Y Iso)
HOT Unilat. (Y in O Trans vs Y Iso)
HPB (Y Het vs Y Iso)
Lysosome
Ribosome
Spliceosome
Cytokine Cytokine Receptor Interaction
Hematopoietic Cell Lineage
Toll Like Receptor Signaling Pathway
Natural Killer Cell Mediated Cytotoxicity
Chemokine Signaling Pathway
Cell Cycle
Regulation of Actin Cytoskeleton
t Cell Receptor Signaling Pathway
Neurotrophin Signaling Pathway
Leukocyte Transendothelial Migration
Endocytosis
Oxidative Phosphorylation
b Cell Receptor Signaling Pathway
Cell Adhesion Molecules Cams
Jak Stat Signaling Pathway
Fc Epsilon Ri Signaling Pathway
Primary Immunodeficiency
NES
B
KEGG: top 20 pathways (Age Decel)
Ovarian Aging (Meta-Sign)
HOT Bilat. (O in Y vs O Iso)
HOT Unilat. (O in Y Trans vs O Iso)
HPB (O Het vs O Iso)
Ribosome
Cytokine Cytokine Receptor Interaction
Hematopoietic Cell Lineage
Natural Killer Cell Mediated Cytotoxicity
Cell Cycle
Parkinsons Disease
Lysosome
Toll Like Receptor Signaling Pathway
Huntingtons Disease
t Cell Receptor Signaling Pathway
b Cell Receptor Signaling Pathway
Fc Epsilon Ri Signaling Pathway
Oxidative Phosphorylation
Primary Immunodeficiency
Chemokine Signaling Pathway
Proteasome
Cell Adhesion Molecules Cams
Spliceosome
Dna Replication
Nod Like Receptor Signaling Pathway
NES
C
Reactome — top 20 pathways (Age Accel)
Ovarian Aging (Meta-Sign)
HOT Bilat. (Y in O vs Y Iso)
HOT Unilat. (Y in O Trans vs Y Iso)
HPB (Y Het vs Y Iso)
Innate Immune System
Neutrophil Degranulation
Major Pathway of rRNA Processing in the Nucleolus and Cytosol
Processing of Capped Intron-Containing Pre-mRNA
Mitotic Prometaphase
Cell Cycle Checkpoints
M Phase
Adaptive Immune System
Resolution of Sister Chromatid Cohesion
Eukaryotic Translation Initiation
Nonsense-Mediated Decay (NMD) Independent of the Exon Junction Complex (EJC)
Immunoregulatory Interactions Between a Lymphoid and a Non-Lymphoid Cell
Nonsense-Mediated Decay (NMD)
Amplification of Signal from the Kinetochores
Mitotic Spindle Checkpoint
Platelet Activation, Signaling and Aggregation
Mitotic Metaphase and Anaphase
Translation
Signaling by Interleukins
DNA Repair
NES
D
Reactome — top 20 pathways (Age Decel)
Ovarian Aging (Meta-Sign)
HOT Bilat. (O in Y vs O Iso)
HOT Unilat. (O in Y Trans vs O Iso)
HPB (O Het vs O Iso)
Innate Immune System
Translation
Neutrophil Degranulation
Major Pathway of rRNA Processing in the Nucleolus and Cytosol
Eukaryotic Translation Initiation
Nonsense-Mediated Decay (NMD) Independent of the Exon Junction Complex (EJC)
Adaptive Immune System
Nonsense-Mediated Decay (NMD)
M Phase
Mitotic Prometaphase
Cell Cycle Checkpoints
Mitotic Metaphase and Anaphase
Immunoregulatory Interactions Between a Lymphoid and a Non-Lymphoid Cell
DNA Replication
S Phase
Separation of Sister Chromatids
G1/S Transition
Resolution of Sister Chromatid Cohesion
Mitotic Spindle Checkpoint
Amplification of Signal from the Kinetochores
NES
E
Top 20 transcription factors (CollecTRI ULM, Age Accel)
Ovarian Aging (Meta-Sign)
HOT Bilat. (Y in O vs Y Iso)
HOT Unilat. (Y in O Trans vs Y Iso)
HPB (Y Het vs Y Iso)
Spi1
Irf1
Nfkb1
Rela
Jun
Ets1
Elf1
Cebpb
Irf4
Cebpg
Tp53
Spic
Mitf
Klf3
Nfe2l2
Stat5a
Irf2
Tbx21
Nfatc2
Spib
ULM score
F
Top 20 transcription factors (CollecTRI ULM, Age Decel)
Ovarian Aging (Meta-Sign)
HOT Bilat. (O in Y vs O Iso)
HOT Unilat. (O in Y Trans vs O Iso)
HPB (O Het vs O Iso)
Spi1
Irf1
Nfkb1
Rela
Myocd
Ets1
Myc
Elf1
Irf4
Jun
Spic
Mitf
Cebpb
Klf3
Stat5a
Irf2
Tbx21
Nfatc2
Cebpg
Spib
ULM score

Ext Fig. 17

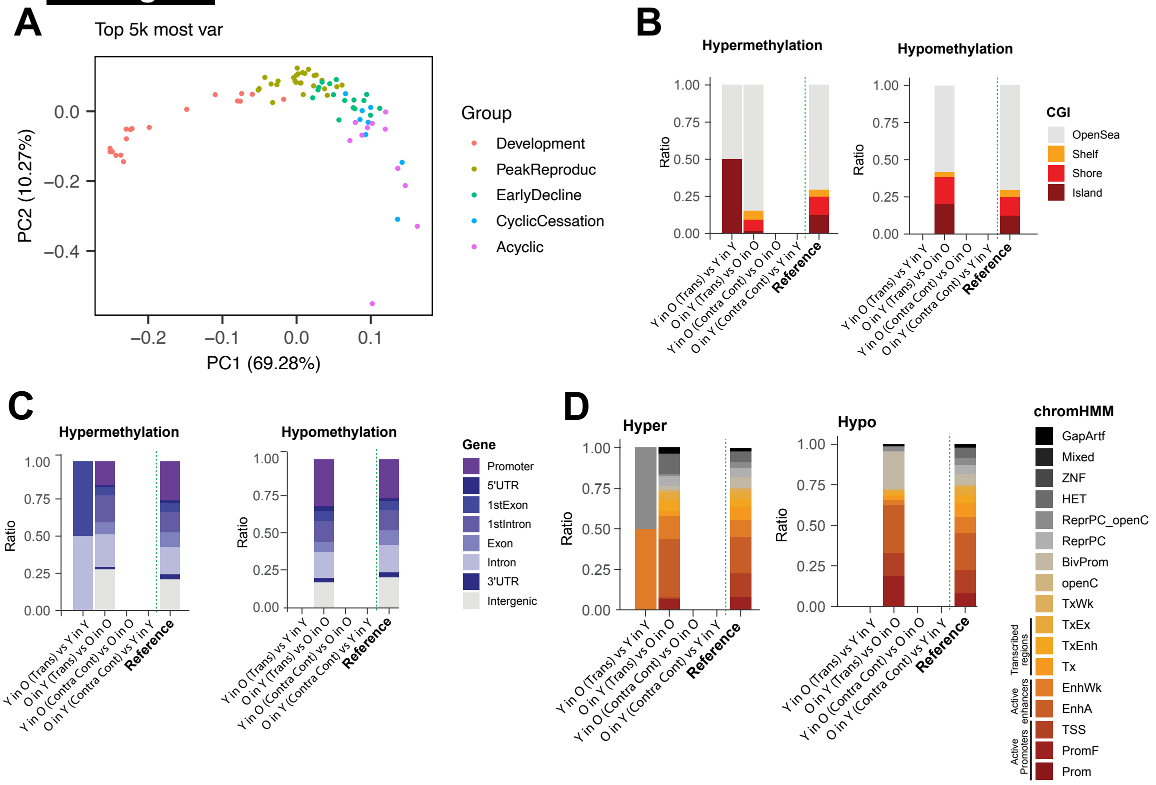
A
Top 5k most var
PC2 (10.27%)
PC1 (69.28%)
0.0
−0.2
−0.4
−0.2
−0.1
0.0
0.1
Group
Development
PeakReproduc
EarlyDecline
CyclicCessation
Acyclic
B
Hypermethylation
Hypomethylation
Ratio
1.00
0.75
0.50
0.25
0.00
Y in O (Trans) vs Y in Y
O in Y (Trans) vs O in O
Y in O (Contra Cont) vs O in O
O in Y (Contra Cont) vs Y in Y
Reference
CGI
OpenSea
Shelf
Shore
Island
C
Hypermethylation
Hypomethylation
Gene
Promoter
5'UTR
1stExon
1stIntron
Exon
Intron
3'UTR
Intergenic
D
Hyper
Hypo
chromHMM
GapArtf
Mixed
ZNF
HET
ReprPC_openC
ReprPC
BivProm
openC
TxWk
TxEx
TxEnh
Tx
EnhWk
EnhA
TSS
PromF
Prom
Transcribed regions
Active enhancers
Active Promoters

### Ext Fig.18

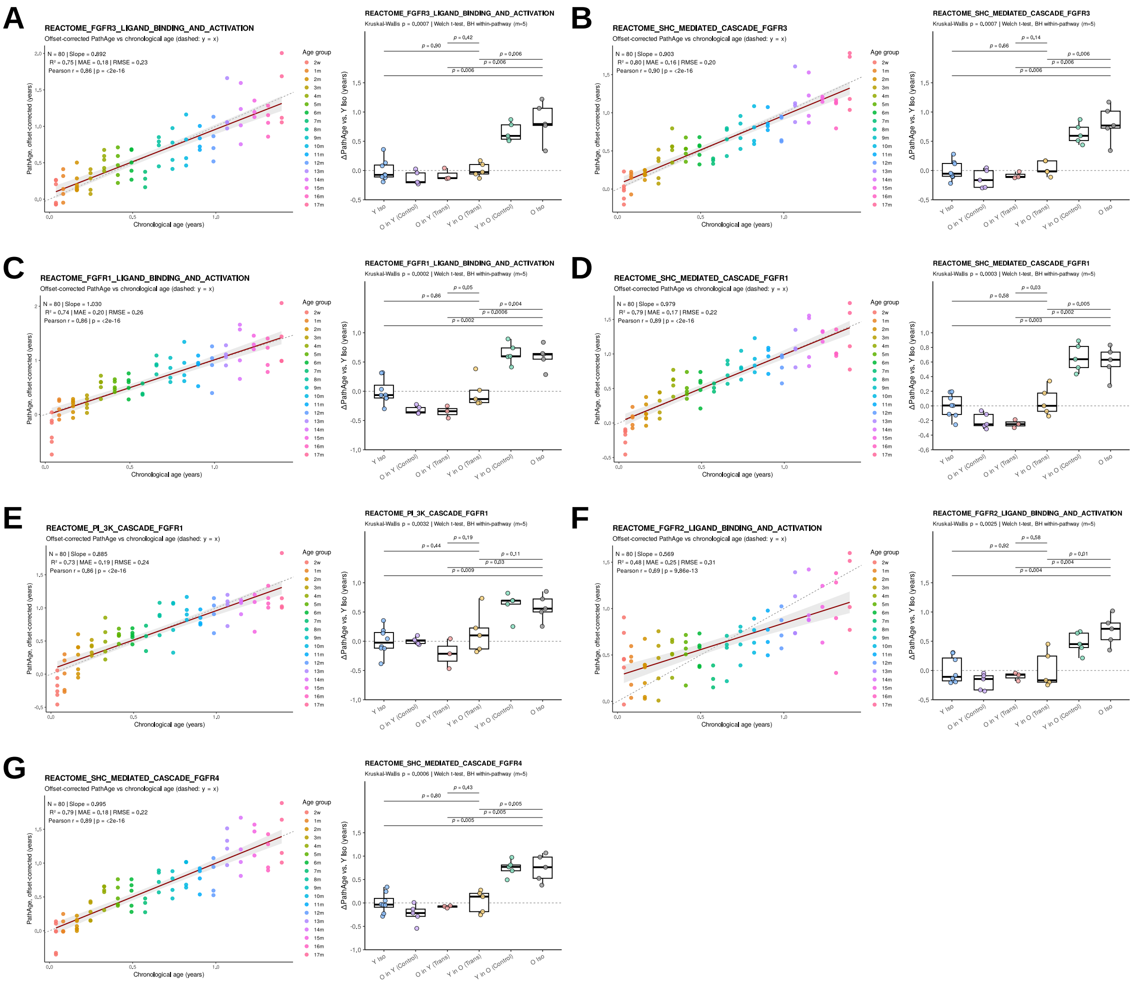
A
REACTOME_FGFR3_LIGAND_BINDING_AND_ACTIVATION
Offset-corrected PathAge vs chronological age (dashed: y = x)
B
REACTOME_SHC_MEDIATED_CASCADE_FGFR3
C
REACTOME_FGFR1_LIGAND_BINDING_AND_ACTIVATION
D
REACTOME_SHC_MEDIATED_CASCADE_FGFR1
E
REACTOME_PI_3K_CASCADE_FGFR1
F
REACTOME_FGFR2_LIGAND_BINDING_AND_ACTIVATION
G
REACTOME_SHC_MEDIATED_CASCADE_FGFR4
Chronological age (years)
PathAge, offset-corrected (years)
ΔPathAge vs. Y Iso (years)
Age group
Y Iso
O in Y (Control)
O in Y (Trans)
Y in O (Trans)
Y in O (Control)
O Iso

### Ext Fig.19

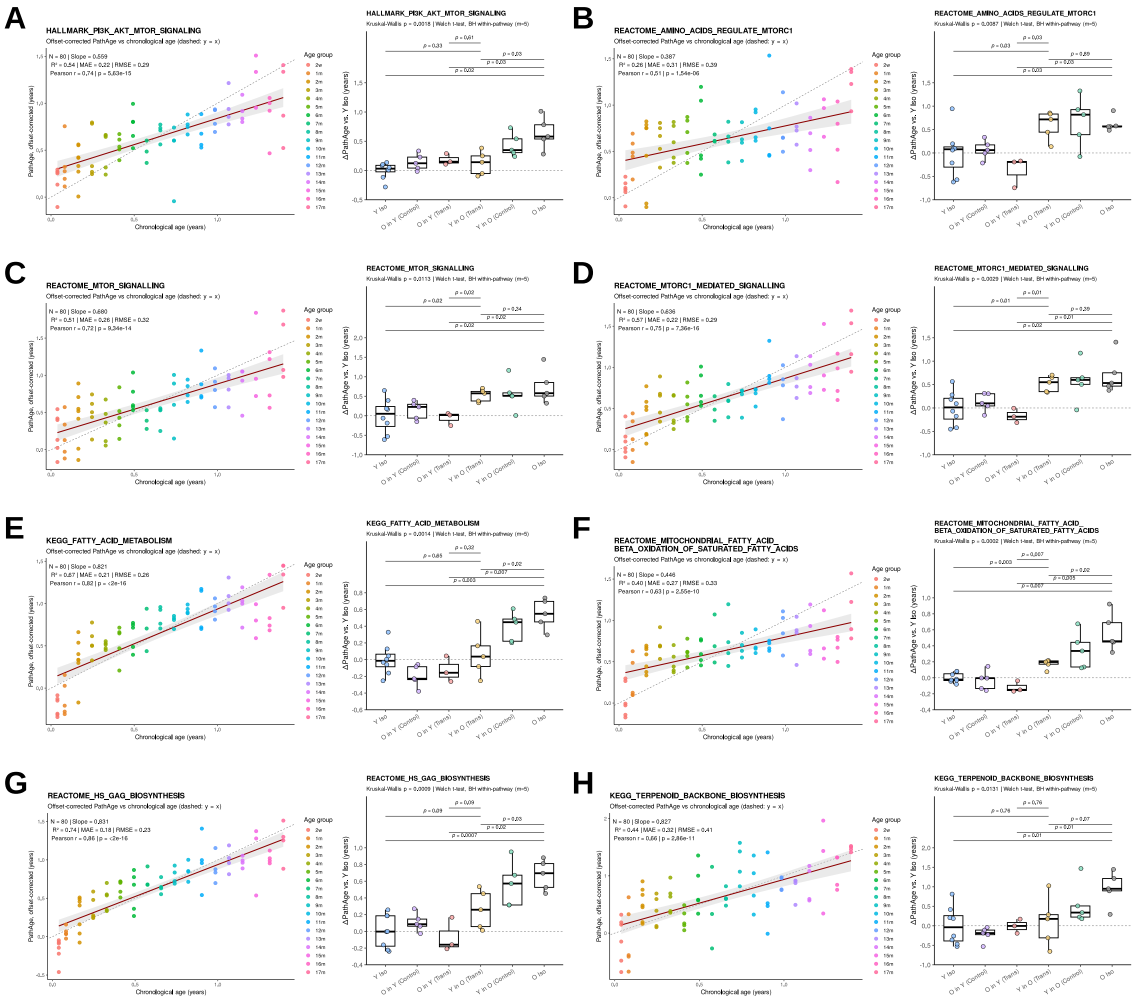
A
HALLMARK_PI3K_AKT_MTOR_SIGNALING
Offset-corrected PathAge vs chronological age (dashed: y = x)
B
REACTOME_AMINO_ACIDS_REGULATE_MTORC1
C
REACTOME_MTOR_SIGNALLING
D
REACTOME_MTORC1_MEDIATED_SIGNALLING
E
KEGG_FATTY_ACID_METABOLISM
F
REACTOME_MITOCHONDRIAL_FATTY_ACID_BETA_OXIDATION_OF_SATURATED_FATTY_ACIDS
G
REACTOME_HS_GAG_BIOSYNTHESIS
H
KEGG_TERPENOID_BACKBONE_BIOSYNTHESIS
Chronological age (years)
PathAge, offset-corrected (years)
ΔPathAge vs. Y Iso (years)
Age group
Y Iso
O in Y (Control)
O in Y (Trans)
Y in O (Trans)
Y in O (Control)
O Iso

### Ext Fig.20

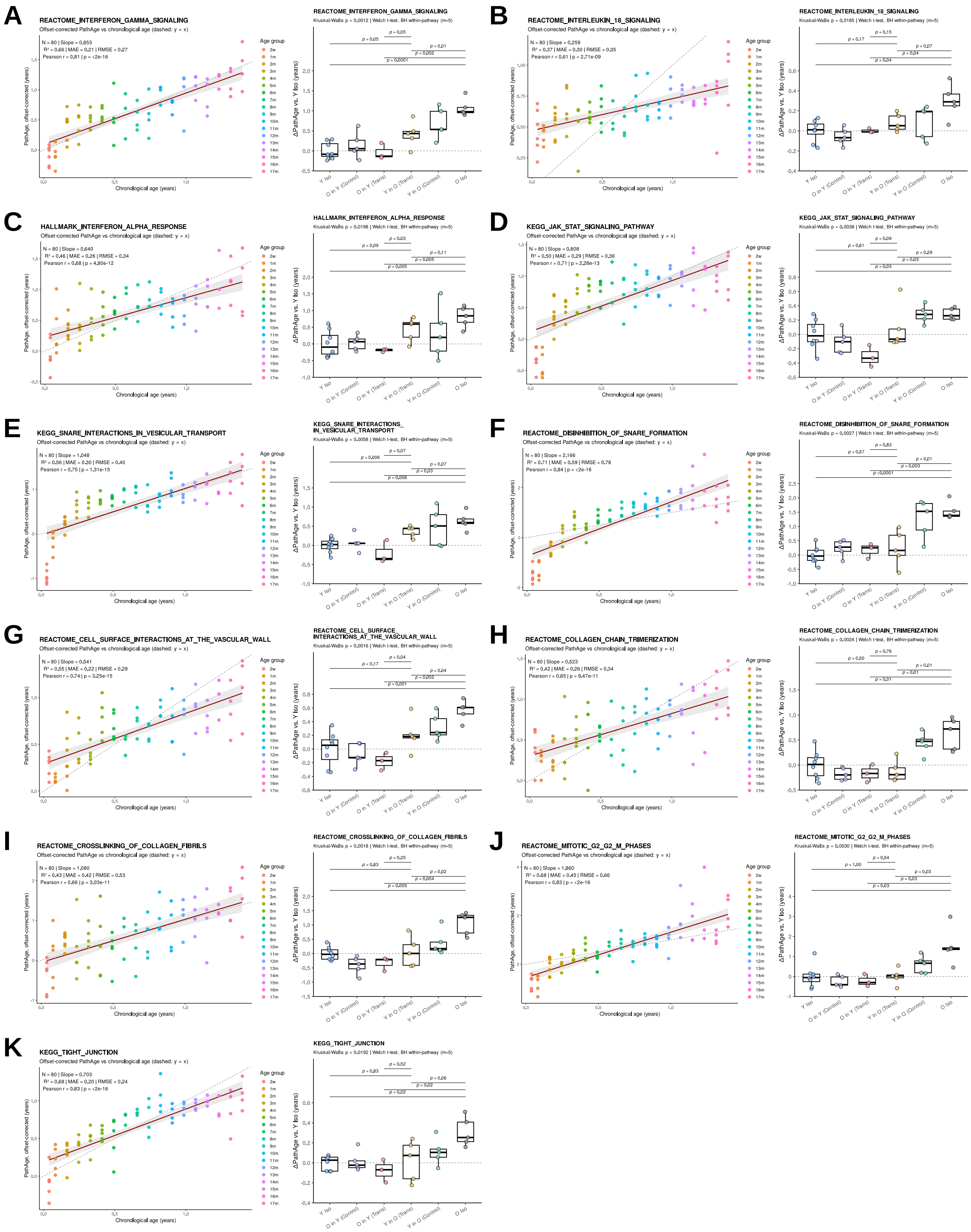

A
REACTOME_INTERFERON_GAMMA_SIGNALING
Offset-corrected PathAge vs chronological age (dashed: y = x)
Kruskal-Wallis p = 0,0012 | Welch t-test, BH within-pathway (m=5)
B
REACTOME_INTERLEUKIN_18_SIGNALING
Kruskal-Wallis p = 0,0185 | Welch t-test, BH within-pathway (m=5)
C
HALLMARK_INTERFERON_ALPHA_RESPONSE
Kruskal-Wallis p = 0,0198 | Welch t-test, BH within-pathway (m=5)
D
KEGG_JAK_STAT_SIGNALING_PATHWAY
Kruskal-Wallis p = 0,0038 | Welch t-test, BH within-pathway (m=5)
E
KEGG_SNARE_INTERACTIONS_IN_VESICULAR_TRANSPORT
Kruskal-Wallis p = 0,0058 | Welch t-test, BH within-pathway (m=5)
F
REACTOME_DISINHIBITION_OF_SNARE_FORMATION
Kruskal-Wallis p = 0,0027 | Welch t-test, BH within-pathway (m=5)
G
REACTOME_CELL_SURFACE_INTERACTIONS_AT_THE_VASCULAR_WALL
Kruskal-Wallis p = 0,0016 | Welch t-test, BH within-pathway (m=5)
H
REACTOME_COLLAGEN_CHAIN_TRIMERIZATION
Kruskal-Wallis p = 0,0024 | Welch t-test, BH within-pathway (m=5)
I
REACTOME_CROSSLINKING_OF_COLLAGEN_FIBRILS
Kruskal-Wallis p = 0,0018 | Welch t-test, BH within-pathway (m=5)
J
REACTOME_MITOTIC_G2_G2_M_PHASES
Kruskal-Wallis p = 0,0030 | Welch t-test, BH within-pathway (m=5)
K
KEGG_TIGHT_JUNCTION
Kruskal-Wallis p = 0,0192 | Welch t-test, BH within-pathway (m=5)
Chronological age (years)
PathAge, offset-corrected (years)
ΔPathAge vs. Y Iso (years)
Age group
Y Iso
O in Y (Control)
O in Y (Trans)
Y in O (Trans)
Y in O (Control)
O Iso

Ext. Fig. 21

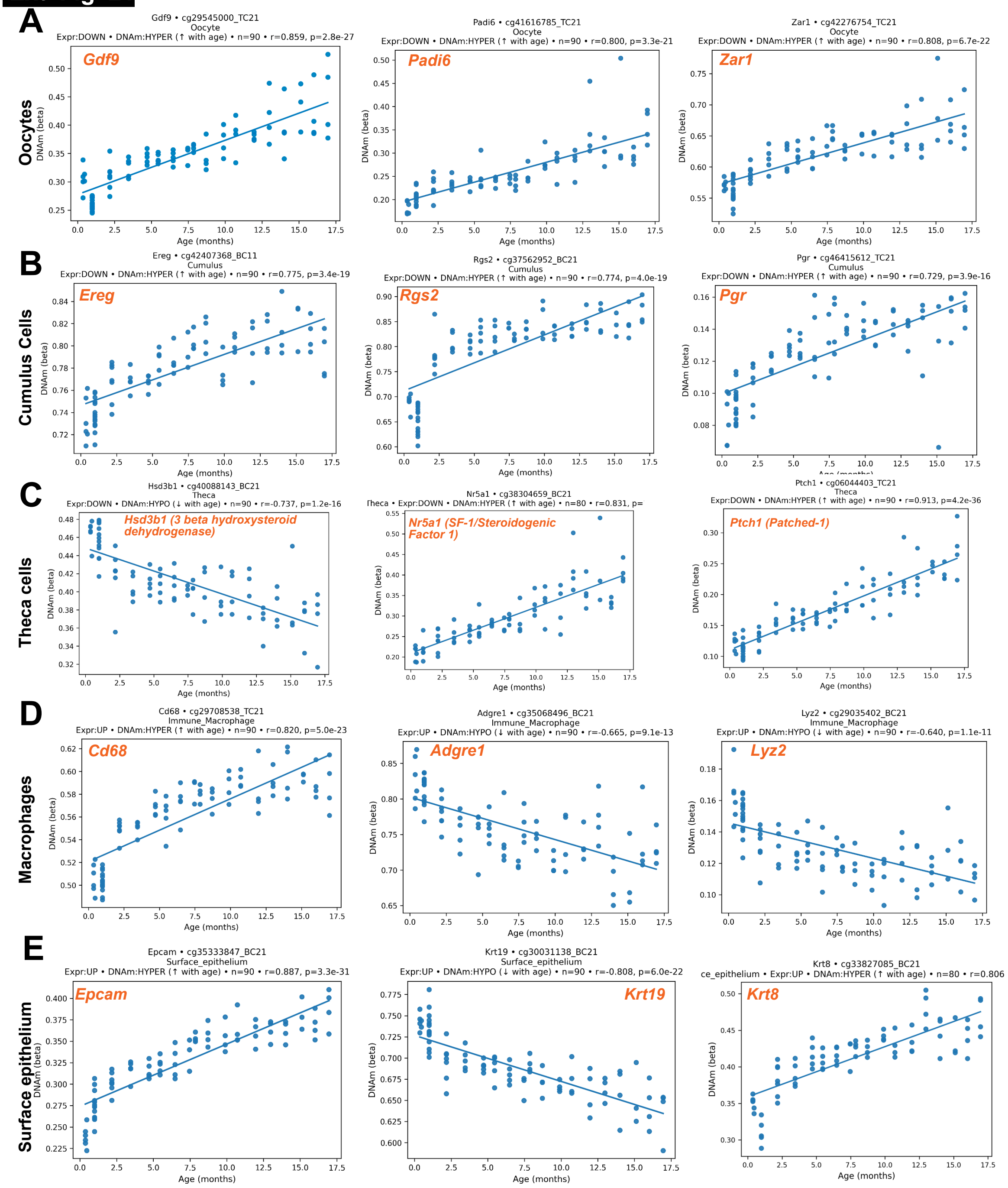

A
Oocytes
Gdf9 • cg29545000_TC21
Oocyte
Expr:DOWN • DNAm:HYPER (↑ with age) • n=90 • r=0.859, p=2.8e-27
Gdf9
Padi6 • cg41616785_TC21
Oocyte
Expr:DOWN • DNAm:HYPER (↑ with age) • n=90 • r=0.800, p=3.3e-21
Padi6
Zar1 • cg42276754_TC21
Oocyte
Expr:DOWN • DNAm:HYPER (↑ with age) • n=90 • r=0.808, p=6.7e-22
Zar1
B
Cumulus Cells
Ereg • cg42407368_BC11
Cumulus
Expr:DOWN • DNAm:HYPER (↑ with age) • n=90 • r=0.775, p=3.4e-19
Ereg
Rgs2 • cg37562952_BC21
Cumulus
Expr:DOWN • DNAm:HYPER (↑ with age) • n=90 • r=0.774, p=4.0e-19
Rgs2
Pgr • cg46415612_TC21
Cumulus
Expr:DOWN • DNAm:HYPER (↑ with age) • n=90 • r=0.729, p=3.9e-16
Pgr
C
Theca cells
Hsd3b1 • cg40088143_BC21
Theca
Expr:DOWN • DNAm:HYPO (↓ with age) • n=90 • r=-0.737, p=1.2e-16
Hsd3b1 (3 beta hydroxysteroid dehydrogenase)
Nr5a1 • cg38304659_BC21
Theca • Expr:DOWN • DNAm:HYPER (↑ with age) • n=80 • r=0.831, p=
Nr5a1 (SF-1/Steroidogenic Factor 1)
Ptch1 • cg06044403_TC21
Theca
Expr:DOWN • DNAm:HYPER (↑ with age) • n=90 • r=0.913, p=4.2e-36
Ptch1 (Patched-1)
D
Macrophages
Cd68 • cg29708538_TC21
Immune_Macrophage
Expr:UP • DNAm:HYPER (↑ with age) • n=90 • r=0.820, p=5.0e-23
Cd68
Adgre1 • cg35068496_BC21
Immune_Macrophage
Expr:UP • DNAm:HYPO (↓ with age) • n=90 • r=-0.665, p=9.1e-13
Adgre1
Lyz2 • cg29035402_BC21
Immune_Macrophage
Expr:UP • DNAm:HYPO (↓ with age) • n=90 • r=-0.640, p=1.1e-11
Lyz2
E
Surface epithelium
Epcam • cg35333847_BC21
Surface_epithelium
Expr:UP • DNAm:HYPER (↑ with age) • n=90 • r=0.887, p=3.3e-31
Epcam
Krt19 • cg30031138_BC21
Surface_epithelium
Expr:UP • DNAm:HYPO (↓ with age) • n=90 • r=-0.808, p=6.0e-22
Krt19
Krt8 • cg33827085_BC21
ce_epithelium • Expr:UP • DNAm:HYPER (↑ with age) • n=80 • r=0.806
Krt8
DNAm (beta)
Age (months)

**Ext Fig. 22**

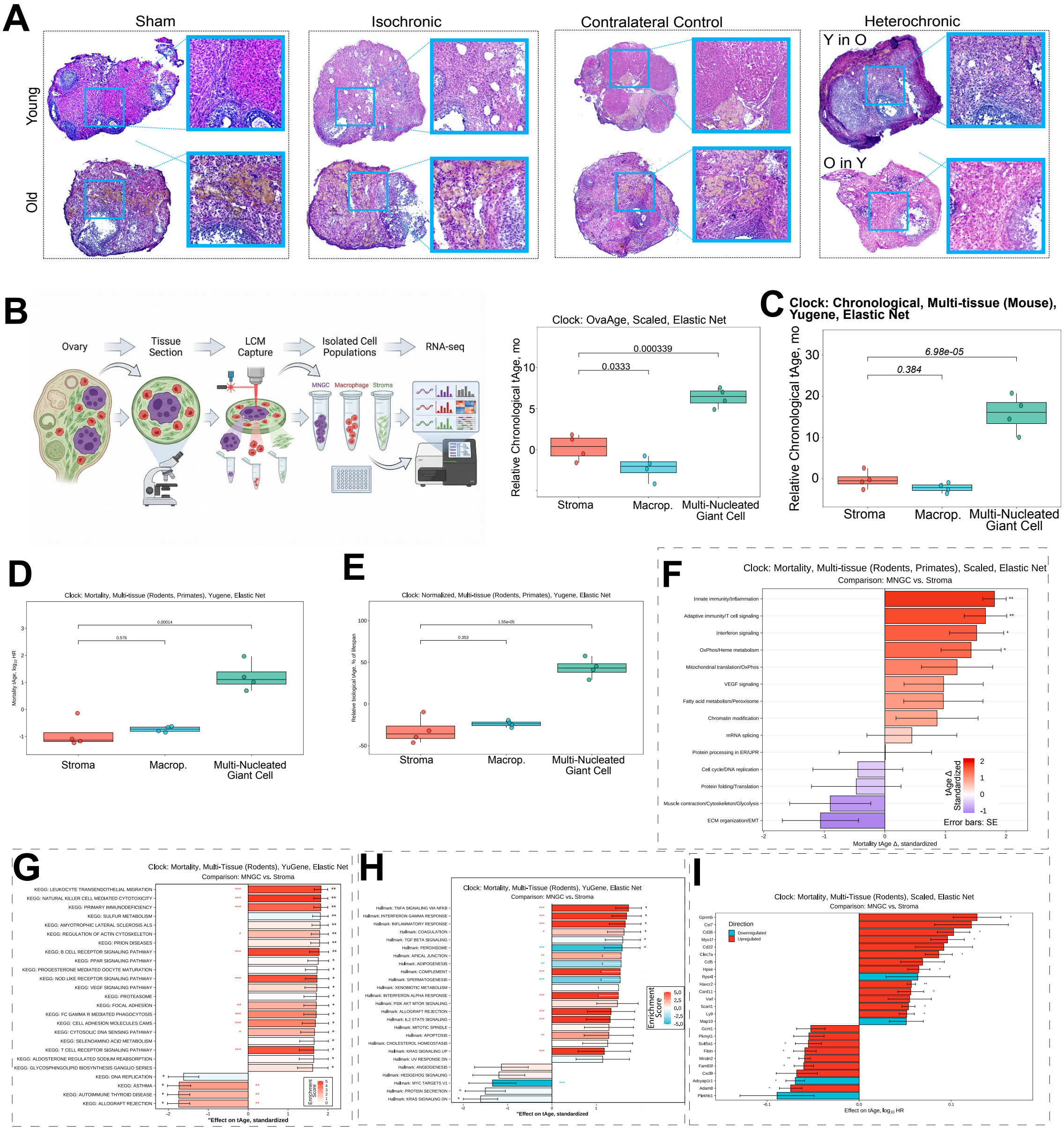

A
Sham
Isochronic
Contralateral Control
Heterochronic
Young
Old
Y in O
O in Y
B
Ovary
Tissue Section
LCM Capture
Isolated Cell Populations
RNA-seq
MNGC
Macrophage
Stroma
Clock: OvaAge, Scaled, Elastic Net
Relative Chronological tAge, mo
0.000339
0.0333
Stroma
Macrop.
Multi-Nucleated Giant Cell
C
Clock: Chronological, Multi-tissue (Mouse), Yugene, Elastic Net
Relative Chronological tAge, mo
6.98e-05
0.384
Stroma
Macrop.
Multi-Nucleated Giant Cell
D
Clock: Mortality, Multi-tissue (Rodents, Primates), Yugene, Elastic Net
Stroma
Macrop.
Multi-Nucleated Giant Cell
E
Clock: Normalized, Multi-tissue (Rodents, Primates), Yugene, Elastic Net
Stroma
Macrop.
Multi-Nucleated Giant Cell
F
Clock: Mortality, Multi-tissue (Rodents, Primates), Scaled, Elastic Net
Comparison: MNGC vs. Stroma
Error bars: SE
G
Clock: Mortality, Multi-Tissue (Rodents), YuGene, Elastic Net
Comparison: MNGC vs. Stroma
H
Clock: Mortality, Multi-Tissue (Rodents), YuGene, Elastic Net
Comparison: MNGC vs. Stroma
Enrichment Score
I
Clock: Mortality, Multi-Tissue (Rodents), Scaled, Elastic Net
Comparison: MNGC vs. Stroma

**Ext Fig. 23**

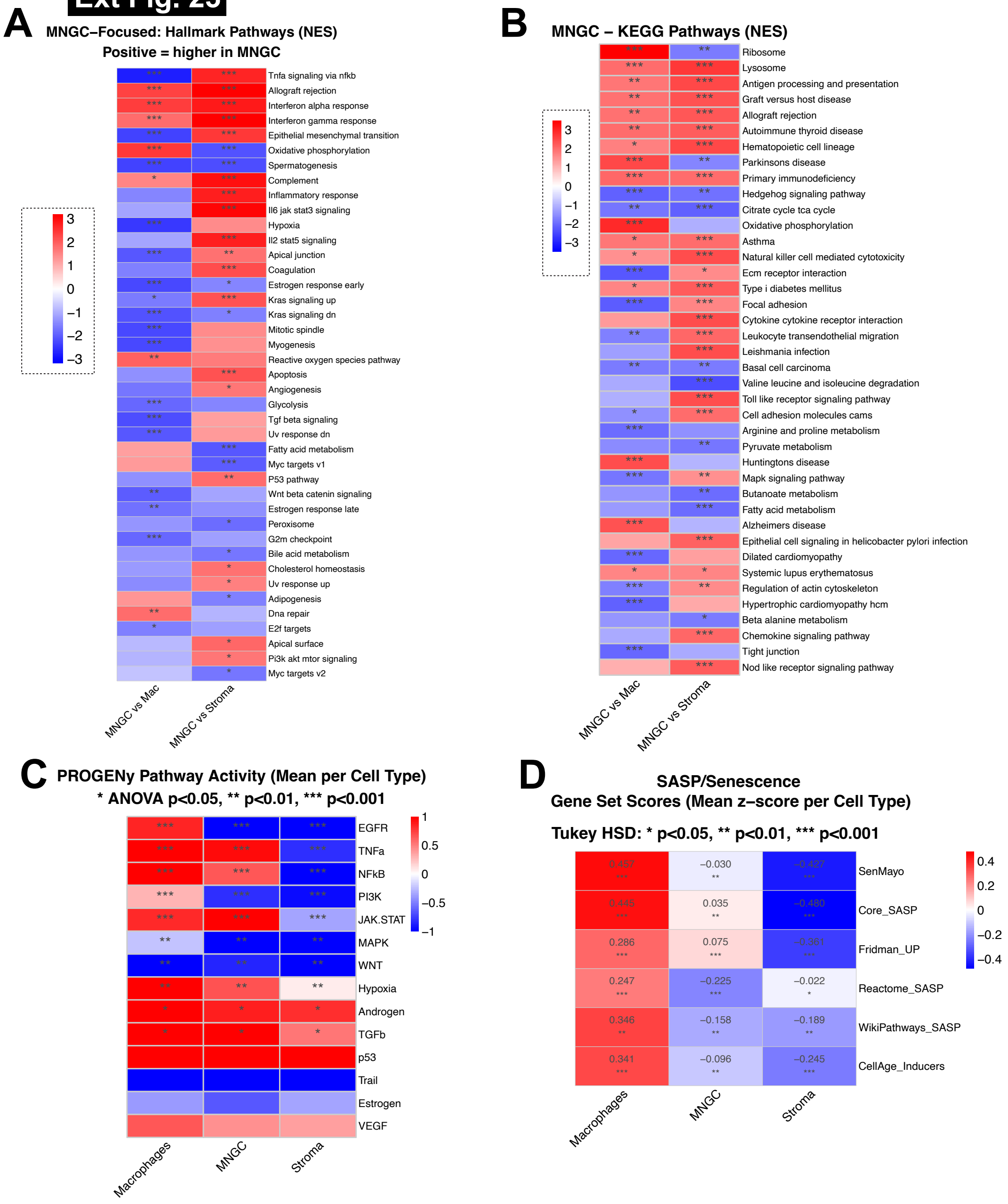
A
MNGC–Focused: Hallmark Pathways (NES)
Positive = higher in MNGC
Tnfa signaling via nfkb
Allograft rejection
Interferon alpha response
Interferon gamma response
Epithelial mesenchymal transition
Oxidative phosphorylation
Spermatogenesis
Complement
Inflammatory response
Il6 jak stat3 signaling
Hypoxia
Il2 stat5 signaling
Apical junction
Coagulation
Estrogen response early
Kras signaling up
Kras signaling dn
Mitotic spindle
Myogenesis
Reactive oxygen species pathway
Apoptosis
Angiogenesis
Glycolysis
Tgf beta signaling
Uv response dn
Fatty acid metabolism
Myc targets v1
P53 pathway
Wnt beta catenin signaling
Estrogen response late
Peroxisome
G2m checkpoint
Bile acid metabolism
Cholesterol homeostasis
Uv response up
Adipogenesis
Dna repair
E2f targets
Apical surface
Pi3k akt mtor signaling
Myc targets v2
MNGC vs Mac
MNGC vs Stroma
B
MNGC – KEGG Pathways (NES)
Ribosome
Lysosome
Antigen processing and presentation
Graft versus host disease
Allograft rejection
Autoimmune thyroid disease
Hematopoietic cell lineage
Parkinsons disease
Primary immunodeficiency
Hedgehog signaling pathway
Citrate cycle tca cycle
Oxidative phosphorylation
Asthma
Natural killer cell mediated cytotoxicity
Ecm receptor interaction
Type i diabetes mellitus
Focal adhesion
Cytokine cytokine receptor interaction
Leukocyte transendothelial migration
Leishmania infection
Basal cell carcinoma
Valine leucine and isoleucine degradation
Toll like receptor signaling pathway
Cell adhesion molecules cams
Arginine and proline metabolism
Pyruvate metabolism
Huntingtons disease
Mapk signaling pathway
Butanoate metabolism
Fatty acid metabolism
Alzheimers disease
Epithelial cell signaling in helicobacter pylori infection
Dilated cardiomyopathy
Systemic lupus erythematosus
Regulation of actin cytoskeleton
Hypertrophic cardiomyopathy hcm
Beta alanine metabolism
Chemokine signaling pathway
Tight junction
Nod like receptor signaling pathway
MNGC vs Mac
MNGC vs Stroma
C
PROGENy Pathway Activity (Mean per Cell Type)
* ANOVA p<0.05, ** p<0.01, *** p<0.001
EGFR
TNFa
NFkB
PI3K
JAK.STAT
MAPK
WNT
Hypoxia
Androgen
TGFb
p53
Trail
Estrogen
VEGF
Macrophages
MNGC
Stroma
D
SASP/Senescence
Gene Set Scores (Mean z–score per Cell Type)
Tukey HSD: * p<0.05, ** p<0.01, *** p<0.001
SenMayo
Core_SASP
Fridman_UP
Reactome_SASP
WikiPathways_SASP
CellAge_Inducers
−0.030
0.035
0.286
0.075
0.247
−0.225
−0.022
0.346
−0.158
−0.189
0.341
−0.096
−0.245
Macrophages
MNGC
Stroma

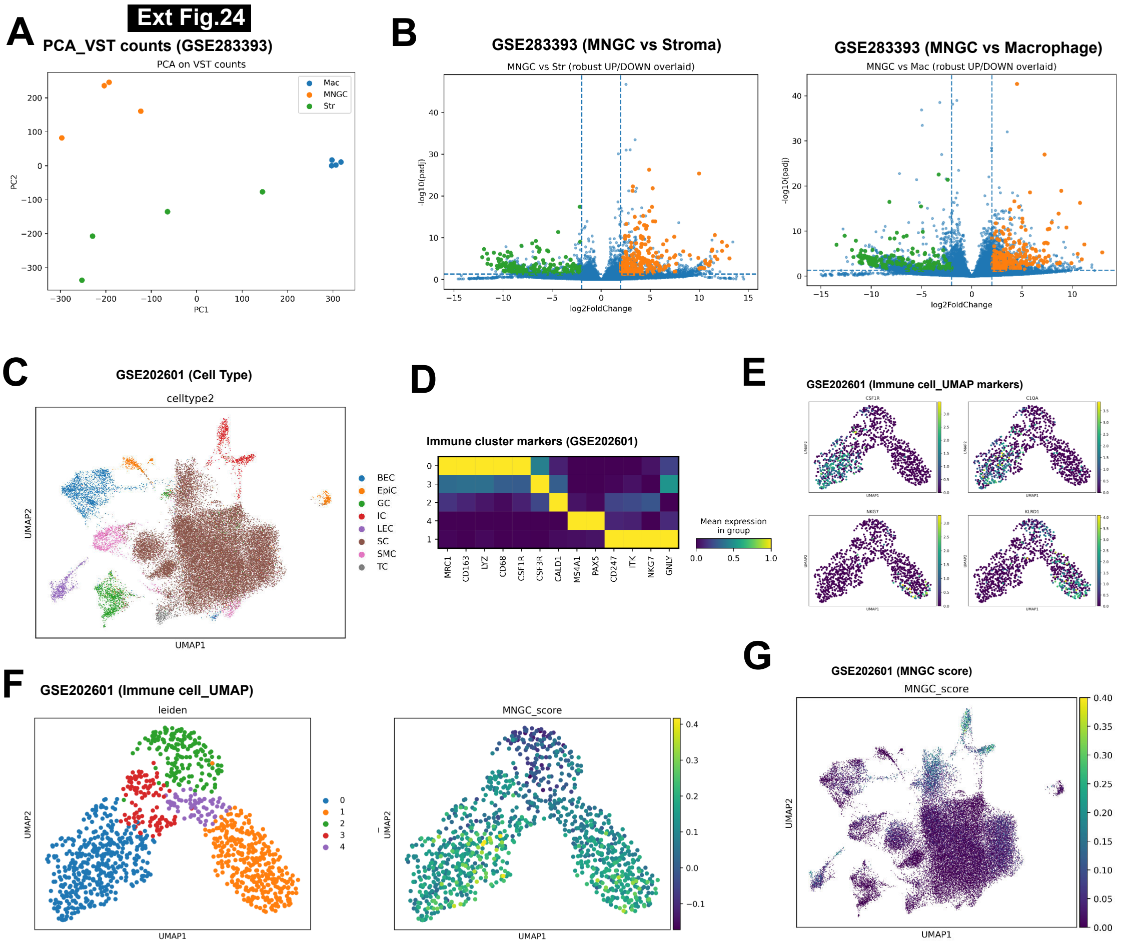

Ext Fig.24
A
PCA_VST counts (GSE283393)
PCA on VST counts
Mac
MNGC
Str
PC1
PC2
B
GSE283393 (MNGC vs Stroma)
MNGC vs Str (robust UP/DOWN overlaid)
GSE283393 (MNGC vs Macrophage)
MNGC vs Mac (robust UP/DOWN overlaid)
log2FoldChange
-log10(padj)
C
GSE202601 (Cell Type)
celltype2
BEC
EpiC
GC
IC
LEC
SC
SMC
TC
UMAP1
UMAP2
D
Immune cluster markers (GSE202601)
MRC1
CD163
LYZ
CD68
CSF1R
CSF3R
CALD1
MS4A1
PAX5
CD247
ITK
NKG7
GNLY
Mean expression in group
E
GSE202601 (Immune cell_UMAP markers)
F
GSE202601 (Immune cell_UMAP)
leiden
MNGC_score
G
GSE202601 (MNGC score)
MNGC_score

Ext Fig.25

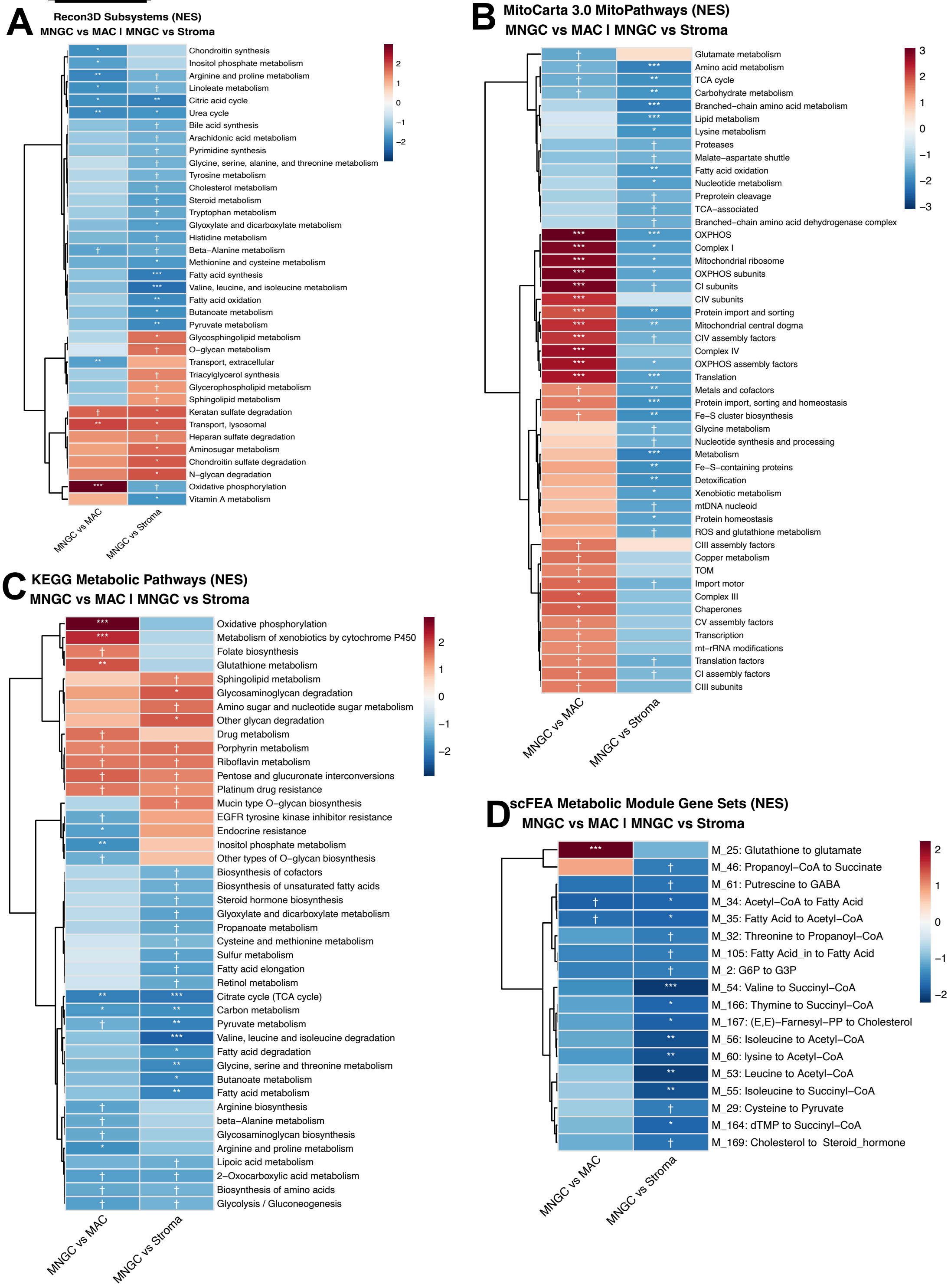
A
Recon3D Subsystems (NES)
MNGC vs MAC I MNGC vs Stroma
Chondroitin synthesis
Inositol phosphate metabolism
Arginine and proline metabolism
Linoleate metabolism
Citric acid cycle
Urea cycle
Bile acid synthesis
Arachidonic acid metabolism
Pyrimidine synthesis
Glycine, serine, alanine, and threonine metabolism
Tyrosine metabolism
Cholesterol metabolism
Steroid metabolism
Tryptophan metabolism
Glyoxylate and dicarboxylate metabolism
Histidine metabolism
Beta-Alanine metabolism
Methionine and cysteine metabolism
Fatty acid synthesis
Valine, leucine, and isoleucine metabolism
Fatty acid oxidation
Butanoate metabolism
Pyruvate metabolism
Glycosphingolipid metabolism
O−glycan metabolism
Transport, extracellular
Triacylglycerol synthesis
Glycerophospholipid metabolism
Sphingolipid metabolism
Keratan sulfate degradation
Transport, lysosomal
Heparan sulfate degradation
Aminosugar metabolism
Chondroitin sulfate degradation
N−glycan degradation
Oxidative phosphorylation
Vitamin A metabolism
MNGC vs MAC
MNGC vs Stroma
2
1
0
−1
−2
B
MitoCarta 3.0 MitoPathways (NES)
MNGC vs MAC I MNGC vs Stroma
Glutamate metabolism
Amino acid metabolism
TCA cycle
Carbohydrate metabolism
Branched−chain amino acid metabolism
Lipid metabolism
Lysine metabolism
Proteases
Malate−aspartate shuttle
Fatty acid oxidation
Nucleotide metabolism
Preprotein cleavage
TCA−associated
Branched−chain amino acid dehydrogenase complex
OXPHOS
Complex I
Mitochondrial ribosome
OXPHOS subunits
CI subunits
CIV subunits
Protein import and sorting
Mitochondrial central dogma
CIV assembly factors
Complex IV
OXPHOS assembly factors
Translation
Metals and cofactors
Protein import, sorting and homeostasis
Fe−S cluster biosynthesis
Glycine metabolism
Nucleotide synthesis and processing
Metabolism
Fe−S−containing proteins
Detoxification
Xenobiotic metabolism
mtDNA nucleoid
Protein homeostasis
ROS and glutathione metabolism
CIII assembly factors
Copper metabolism
TOM
Import motor
Complex III
Chaperones
CV assembly factors
Transcription
mt−rRNA modifications
Translation factors
CI assembly factors
CIII subunits
MNGC vs MAC
MNGC vs Stroma
3
2
1
0
−1
−2
−3
C
KEGG Metabolic Pathways (NES)
MNGC vs MAC I MNGC vs Stroma
Oxidative phosphorylation
Metabolism of xenobiotics by cytochrome P450
Folate biosynthesis
Glutathione metabolism
Sphingolipid metabolism
Glycosaminoglycan degradation
Amino sugar and nucleotide sugar metabolism
Other glycan degradation
Drug metabolism
Porphyrin metabolism
Riboflavin metabolism
Pentose and glucuronate interconversions
Platinum drug resistance
Mucin type O−glycan biosynthesis
EGFR tyrosine kinase inhibitor resistance
Endocrine resistance
Inositol phosphate metabolism
Other types of O−glycan biosynthesis
Biosynthesis of cofactors
Biosynthesis of unsaturated fatty acids
Steroid hormone biosynthesis
Glyoxylate and dicarboxylate metabolism
Propanoate metabolism
Cysteine and methionine metabolism
Sulfur metabolism
Fatty acid elongation
Retinol metabolism
Citrate cycle (TCA cycle)
Carbon metabolism
Pyruvate metabolism
Valine, leucine and isoleucine degradation
Fatty acid degradation
Glycine, serine and threonine metabolism
Butanoate metabolism
Fatty acid metabolism
Arginine biosynthesis
beta−Alanine metabolism
Glycosaminoglycan biosynthesis
Arginine and proline metabolism
Lipoic acid metabolism
2−Oxocarboxylic acid metabolism
Biosynthesis of amino acids
Glycolysis / Gluconeogenesis
MNGC vs MAC
MNGC vs Stroma
2
1
0
−1
−2
D
scFEA Metabolic Module Gene Sets (NES)
MNGC vs MAC I MNGC vs Stroma
M_25: Glutathione to glutamate
M_46: Propanoyl−CoA to Succinate
M_61: Putrescine to GABA
M_34: Acetyl−CoA to Fatty Acid
M_35: Fatty Acid to Acetyl−CoA
M_32: Threonine to Propanoyl−CoA
M_105: Fatty Acid_in to Fatty Acid
M_2: G6P to G3P
M_54: Valine to Succinyl−CoA
M_166: Thymine to Succinyl−CoA
M_167: (E,E)−Farnesyl−PP to Cholesterol
M_56: Isoleucine to Acetyl−CoA
M_60: lysine to Acetyl−CoA
M_53: Leucine to Acetyl−CoA
M_55: Isoleucine to Succinyl−CoA
M_29: Cysteine to Pyruvate
M_164: dTMP to Succinyl−CoA
M_169: Cholesterol to Steroid_hormone
MNGC vs MAC
MNGC vs Stroma
2
1
0
−1
−2

**Ext Fig.26**

**Key Meta-Signature Correlations: Scatter Plots**
**(Green = significant, Pink = not significant)**

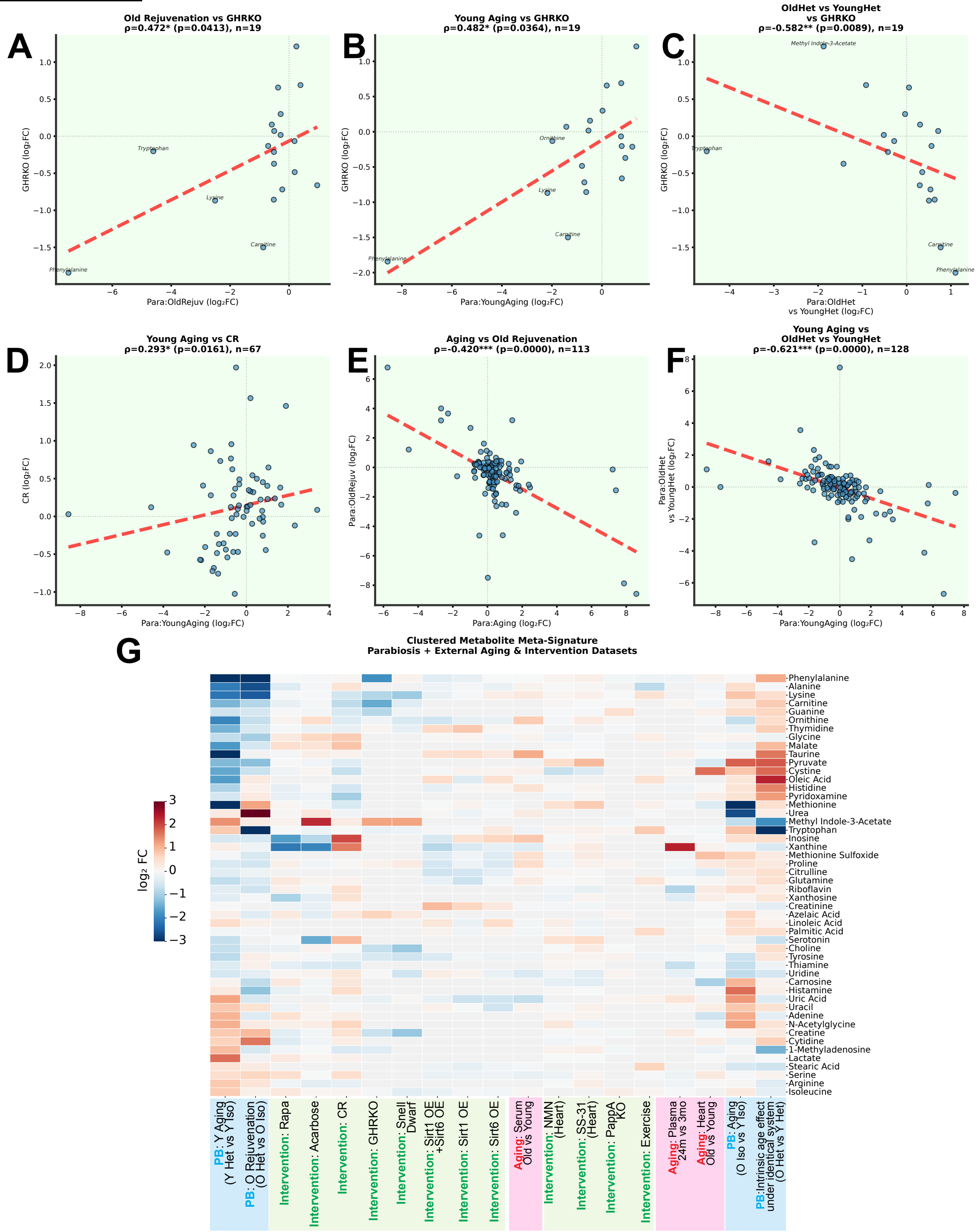

#### Extended Data figure legends

**Extended Data Figure 1 | Gene- and module-level components of ovarian transcriptomic aging and multi-tissue clock performance**

(A) Gene-level chronological-age slopes in the ovarian lifespan series against −log10 BH-adjusted P (negative binomial generalized linear model with a likelihood-ratio test in edgeR; Robinson et al, 2010). Red, positive slopes; blue, negative slopes; selected genes are labeled.

(B) Additional contribution of each co-regulated module (Tyshkovskiy *et al*, 2026) to the ovarian mortality-clock tAge slope during development (0.5–2 months) relative to the adult interval ("extra slope during development"). Bar color shows this slope; asterisks show BH-adjusted significance.

(C) Mortality tAge (log10 hazard ratio) from the innate immunity/inflammation module-specific clock against age in ovary.

(D) Rodent multi-tissue chronological clock (Tyshkovskiy *et al*, 2026) applied to ovary, brain and liver from the same mice. Slope = 1.6, r = 0.87, $P = 8.3 \times 10^{-54}$ (ovary); slope = 1.6, r = 0.97 (brain); slope = 1.3, r = 0.92, $P = 6.5 \times 10^{-33}$ (liver).

(E) Spearman correlations between age-adjusted transcriptomic age estimates (residuals) of ovary, brain and liver from the same animals.

Points are individual samples; lines show linear fits with 95% confidence bands. *P < 0.05, **P < 0.01, ***P < 0.001.

**Extended Data Figure 2 | OvaAge in independent ovarian datasets**

(A) OvaAge in ovaries from young and old wild-type (WT) and Ames dwarf mice (GSE84078; Schneider *et al*, 2017); old WT versus old Ames dwarf P = 0.0042.

(B) Reconstructed follicles (RCF) containing young or aged oocytes in young or aged follicular environments (GSE270016; Wang *et al*, 2024).

(C) Mouse granulosa cells from the single-cell ovarian aging atlas (GSE232309; Isola *et al*, 2024), young versus old.

(D) Cynomolgus macaque ovarian somatic cells (GSE130664; Wang *et al*, 2020), young versus old; $P = 1.4 \times 10^{-4}$.

(E) Standardized differences in module-specific tAge (tAge Δ) between young and old macaque ovarian somatic cells.

Box plots show the median and interquartile range; points are individual samples. Pairwise P values are shown above brackets. Predictions for other species and cell compartments are relative comparisons made with a mouse whole-ovary model.

**Extended Data Figure 3 | Ovarian DNA methylation measures across age**

(A) Universal3 eAge against chronological age (Lu *et al*, 2023); slope = 0.86, Pearson r = 0.96.

(B) Methylation entropy against chronological age; r = 0.80.

(C) Mean methylation against chronological age; r = 0.74.

Points are individual ovaries; lines show linear fits with 95% confidence bands. Slope, r and P are shown on each plot.

**Extended Data Figure 4 | Estimated ovarian cell-type proportions across age**

(A–K) Deconvolution estimates of oocytes (A), granulosa 1 (B), granulosa 2 (C), endothelium (D), epithelium (E), luteal cells (F), theca (G), phagocytes (H), T lymphocytes (I), stroma 3 (J) and B lymphocytes (K), using the mouse ovarian single-cell reference (Isola *et al*, 2024). Lines show the mean estimated proportion at each age and shaded bands ± 1 s.d. Panel titles give the Spearman correlation with age and BH-adjusted significance (*P < 0.05, **P < 0.01, ***P < 0.001).

**Extended Data Figure 5 | Ovarian expression and methylation after unilateral and bilateral transplantation**

(A) PCA of unilateral ovarian transcriptomes, colored by group and shaped by original ovary age. Contralateral control denotes the endogenous ovary retained in a transplanted recipient; in O in Y recipients it is young, and in Y in O recipients it is old.

(B) PCA of the 50,000 most variable CpGs in unilateral transplantation samples, colored by group and shaped by recipient age.

(C) PCA of bilateral ovarian transcriptomes, colored by group and shaped by original ovary age.

(D) GSEA (Hallmark, KEGG and Reactome) of bilateral transplantation signatures from ovary, brain and uterus, alongside reference aging and longevity-intervention signatures (Tyshkovskiy *et al*, 2026). Pink labels give the reference group for each contrast; color shows NES.

Group names give ovary (donor) age before recipient age: Y in O, young ovary in an old recipient; O in Y, old ovary in a young recipient; Y in Y and O in O, isochronic grafts (Y Iso, O Iso). PCA axes give the percentage of variance explained. In D, *P < 0.05, **P < 0.01, ***P < 0.001 (BH-adjusted); ^P < 0.1.

**Extended Data Figure 6 | Estimated cell-type proportions after unilateral transplantation**

(A–C) Deconvolution-estimated proportions of epithelial cells (A), phagocytes (B) and oocytes (C) in the unilateral transplantation ovaries: young sham, Y in Y, the contralateral (endogenous young) ovary and the graft of O-in-Y recipients, the graft and the contralateral (endogenous old) ovary of Y-in-O recipients, O in O and old sham (n = 9, 3, 4, 3, 5, 5, 5 and 4; 38 ovaries). Brackets mark the pairwise comparisons tested in the original analysis (A, six pairs; B, four pairs; C, both sets). P values are from two-sided Dunn tests, BH-adjusted over all 28 pairs; values below 0.05 are in bold. Kruskal–Wallis P = 0.0073 (A), 0.0070 (B) and 0.30 (C). (D) The remaining estimated cell types (granulosa 1 and 2, stroma 2 and 3, endothelium, luteal, theca, T and B lymphocytes). Asterisks mark two-sided Dunn BH-adjusted P < 0.05: stroma 3, young sham versus O-in-Y graft (P = 0.046) and versus Y-in-O contralateral (P = 0.012); luteal, young sham versus Y-in-O contralateral (P = 0.037). Stroma 1 was estimated as zero in every sample.

Group names give ovary (donor) age before recipient age: Y in O, young ovary in an old recipient; O in Y, old ovary in a young recipient; Y in Y and O in O, isochronic grafts (Y Iso, O Iso). Box plots show the median and interquartile range; points are individual samples.

**Extended Data Figure 7 | Multi-tissue transcriptomic clocks applied to transplanted ovaries**

Bayesian ridge multi-tissue clocks (Tyshkovskiy *et al*, 2026) were applied to ovarian data without retraining.

(A, B) Unilateral transplantation: chronological tAge (A, months) and mortality tAge (B, log10 hazard ratio). Grafts and contralateral ovaries are shown separately.

(C, D) Bilateral transplantation: chronological (C) and mortality (D) tAge.

Group names give ovary (donor) age before recipient age: Y in O, young ovary in an old recipient; O in Y, old ovary in a young recipient; Y in Y and O in O, isochronic grafts (Y Iso, O Iso). Box plots show the median and interquartile range; points are individual samples. BH-adjusted pairwise P values are shown above brackets.

**Extended Data Figure 8 | Metabolic gene-set enrichment in old ovaries transplanted into young recipients**

(A–D) GSEA of Recon3D subsystems (A) (Brunk *et al*, 2018), KEGG metabolic pathways (B) (Kanehisa *et al*, 2021), MitoCarta 3.0 pathways (C) (Rath *et al*, 2021) and scFEA-defined metabolic modules (D) (Alghamdi *et al*, 2021). Left column, the ovarian aging meta-signature (old vs young, pooled across ovarian aging datasets). Right column, unilateral O in Y grafts (n = 3) versus old unilateral controls (O Iso and O Sham, n = 5), with genes ranked by the DESeq2 Wald statistic.

Color shows NES; positive values indicate enrichment in the first group of each contrast. ***FDR < 0.001, **FDR < 0.01, *FDR < 0.05, †FDR < 0.25. Panel D reports enrichment of module-associated genes.

**Extended Data Figure 9 | Metabolic gene-set enrichment in young ovaries transplanted into old recipients**

(A–D) GSEA of Recon3D subsystems (A), KEGG metabolic pathways (B), MitoCarta 3.0 pathways (C) and scFEA-defined metabolic modules (D). Left column, the ovarian aging meta-signature (old vs young, pooled). Right column, unilateral Y in O grafts versus young controls (Y Iso and Y Sham), as labeled on the lower axis.

Color shows NES. ***FDR < 0.001, **FDR < 0.01, *FDR < 0.05, †FDR < 0.25. Panel D reports enrichment of module-associated genes.

**Extended Data Figure 10 | Cortical expression after ovariectomy and estradiol treatment in rhesus macaques**

(A) PCA of prefrontal cortex (PFC) expression in intact, OVX and OVX+E2 macaques (Cervera-Juanes *et al*, 2024); ellipses show 95% confidence regions. PERMANOVA P = 0.22.

(B) Mean log2 fold changes of curated gene groups (inflammation/complement, synaptic, mitochondrial, glial, hormone response, aging/senescence) in PFC and occipital cortex (OC). Asterisks show one-sample t-tests against 0.

(C) Spearman correlations between macaque KEGG NES profiles and human aging and neurodegenerative-disease signatures (Alzheimer's disease (AD), Huntington's disease (HD) and Parkinson's disease (PD); GEO accessions and sample sizes are shown on the plot).

(D, E) Hallmark (D) and KEGG (E) GSEA across regions and contrasts. Side bars classify pathways as concordant, discordant, rescued or not significant (NS).

(F) Transcription-factor activity (DoRothEA regulons, ULM) (Garcia-Alonso *et al*, 2019; Badia-i-Mompel *et al*, 2022).

Contrasts: OVX versus intact, OVX+E2 versus intact, and OVX+E2 versus OVX. *P < 0.05, **P < 0.01, ***P < 0.001.

**Extended Data Figure 11 | Brain responses to ovarian donor and recipient age**

(A) Standardized effects of KEGG and Hallmark gene sets on brain tAge (normalized mouse multi-tissue clock), Y in O versus O in O; bar color shows the enrichment score (key), error bars show s.e.

(B) PROGENy pathway activity (t statistic) (Schubert *et al*, 2018).

(C, D) Hallmark (C) and KEGG (D) GSEA; color shows NES.

(E) Transcription-factor activity (DoRothEA mouse regulons, ULM score) (Garcia-Alonso *et al*, 2019; Badia-i-Mompel *et al*, 2022).

Heatmap columns: brain aging meta-signature (public mouse brain aging datasets; Methods), old ovary effect (O in Y vs Y in Y), young ovary effect (Y in O vs O in O) and aging under a young ovary (Y in O vs Y in Y). Group names give ovary (donor) age before recipient age: Y in O, young ovary in an old recipient; O in Y, old ovary in a young recipient; Y in Y and O in O, isochronic grafts (Y Iso, O Iso). ^P < 0.1, *P < 0.05, **P < 0.01, ***P < 0.001 (BH-adjusted).

**Extended Data Figure 12 | Pituitary responses to ovary transplantation**

(A) Chronological tAge from the rodent multi-tissue clock (Tyshkovskiy *et al*, 2026) in pituitary after bilateral transplantation. O in Y versus Y in Y P = 0.78; Y in O versus O in O P = 0.36; Young Sham versus Old Sham P = 0.047.

(B, C) Hallmark (B) and KEGG (C) GSEA.

(D) Transcription-factor activity (DoRothEA, ULM score).

(E) PROGENy pathway activity.

(F, G) GSEA of Recon3D subsystems (F) and KEGG metabolic pathways (G).

Heatmap columns: pituitary aging meta-signature, O in Y versus Y in Y, Y in O versus O in O, and Y in O versus Y in Y. Group names give ovary (donor) age before recipient age: Y in O, young ovary in an old recipient; O in Y, old ovary in a young recipient; Y in Y and O in O, isochronic grafts (Y Iso, O Iso). Box plots show the median and interquartile range; points are individual samples. *P < 0.05, **P < 0.01, ***P < 0.001.

**Extended Data Figure 13 | Liver responses to ovary transplantation**

(A) Mortality tAge (log10 hazard ratio) from the YuGene-normalized rodent multi-tissue Bayesian ridge clock (Tyshkovskiy *et al*, 2026) in liver. Y in O versus O in O $P = 1.47 \times 10^{-5}$.

(B) Standardized effects of KEGG, Hallmark and module gene sets on liver tAge (normalized rodent multi-tissue clock), Y in O versus O in O; bar color shows the enrichment score (key), error bars show s.e.

(C) PROGENy pathway activity.

(D, E) Hallmark (D) and KEGG (E) GSEA.

(F) Transcription-factor activity (DoRothEA, ULM score).

Heatmap columns: liver aging meta-signature, old ovary effect (O in Y vs Y in Y), young ovary effect (Y in O vs O in O) and aging under a young ovary (Y in O vs Y in Y). Group names give ovary (donor) age before recipient age: Y in O, young ovary in an old recipient; O in Y, old ovary in a young recipient; Y in Y and O in O, isochronic grafts (Y Iso, O Iso). Box plots show the median and interquartile range; points are individual samples. ^P < 0.1, *P < 0.05, **P < 0.01, ***P < 0.001.

**Extended Data Figure 14 | Uterine responses to ovary transplantation**

(A) Mortality tAge (log10 hazard ratio) from the multi-tissue elastic net clock trained on rodents and primates (Tyshkovskiy *et al*, 2026) in uterus after bilateral transplantation. Y in O versus O in O P = 0.014.

(B, C) Standardized effects of KEGG and Hallmark gene sets on uterine tAge (normalized multi-tissue clock), for Y in O versus O in O (B) and O in Y versus Y in Y (C). Bar color shows the enrichment score; error bars show s.e.

(D) PROGENy pathway activity.

(E) Hallmark GSEA.

(F) GSEA of KEGG metabolic pathways.

Heatmap columns: uterine aging meta-signature, O in Y versus Y in Y, Y in O versus O in O, and Y in O versus Y in Y. Group names give ovary (donor) age before recipient age: Y in O, young ovary in an old recipient; O in Y, old ovary in a young recipient; Y in Y and O in O, isochronic grafts (Y Iso, O Iso). Box plots show the median and interquartile range; points are individual samples. ^P < 0.1, *P < 0.05, **P < 0.01, ***P < 0.001.

**Extended Data Figure 15 | Ovarian transcriptomic age after heterochronic parabiosis**

(A) Design: mice aged 3.5 and 7.5 months were co-housed, joined for 3 months and separated for 1 month before collection (7.5 and 11.5 months).

(B, C) OvaAge from the scaled elastic net (B) and Bayesian ridge (C) models. Y Het versus Y Iso P = 0.0026 (B) and $3.6 \times 10^{-5}$ (C); O Het versus O Iso P = 0.12 (B) and 0.127 (C).

(D) Innate immunity/inflammation module-specific clock (YuGene-normalized elastic net) in young isochronic and young heterochronic partners; P = 0.0020.

Iso, isochronic pair; Het, heterochronic pair. Box plots show the median and interquartile range; points are individual samples. FDR-adjusted pairwise P values are shown above brackets.

**Extended Data Figure 16 | KEGG, Reactome and transcription-factor responses shared across ovarian interventions**

(A, B) KEGG GSEA for the age-acceleration (A: Y in O vs Y Iso; Y in O Trans vs Y Iso; Y Het vs Y Iso) and age-deceleration (B: O in Y vs O Iso; O in Y Trans vs O Iso; O Het vs O Iso) contrasts, alongside the ovarian aging reference signature.

(C, D) Reactome GSEA for the same contrasts.

(E, F) CollecTRI-based transcription-factor activity (ULM score) for the same contrasts (Müller-Dott *et al*, 2023).

Rows are the 20 top-ranked pathways or transcription factors. HOT, heterochronic ovary transplantation (bilateral or unilateral); HPB, heterochronic parabiosis. Color shows NES (A–D) or ULM score (E, F). *P < 0.05, **P < 0.01, ***P < 0.001 (BH-adjusted).

**Extended Data Figure 17 | Ovarian methylome across age and genomic context of transplantation-associated methylation changes**

(A) PCA of the 5,000 most variable CpGs across the ovarian lifespan, colored by reproductive stage (development, peak reproductive, early decline, cyclic cessation, acyclic). PC1 and PC2 explain 69.28% and 10.27% of the variance.

(B–D) Hypermethylated and hypomethylated differentially methylated positions (DMPs) in the unilateral transplantation contrasts, annotated by CpG-island context (B), gene feature (C) and ChromHMM chromatin state (D). The contrasts are Y in O (Trans) vs Y in Y, O in Y (Trans) vs O in O, Y in O (Contra Cont) vs O in O, and O in Y (Contra Cont) vs Y in Y. Reference bars show the array background. The bars give annotation fractions.

Group names give ovary (donor) age before recipient age: Y in O, young ovary in an old recipient; O in Y, old ovary in a young recipient; Y in Y and O in O, isochronic grafts (Y Iso, O Iso). Trans, graft; Contra Cont, contralateral endogenous ovary.

**Extended Data Figure 18 | FGFR-related PathAge clocks in the lifespan series and after unilateral transplantation**

(A) REACTOME_FGFR3_LIGAND_BINDING_AND_ACTIVATION (n = 80; slope = 0.89, $R^2$ = 0.75, MAE = 0.18 years, r = 0.86, $P < 2 \times 10^{-16}$). Mean ΔPathAge of O in Y grafts versus O Iso is −0.91 years; Kruskal–Wallis P = 0.0007.

(B–D) REACTOME_SHC_MEDIATED_CASCADE_FGFR3 (B; r = 0.90), REACTOME_FGFR1_LIGAND_BINDING_AND_ACTIVATION (C; r = 0.86) and REACTOME_SHC_MEDIATED_CASCADE_FGFR1 (D; r = 0.89); all $P < 2 \times 10^{-16}$. O in Y versus O Iso: −0.85 to −0.94 years; Kruskal–Wallis P ≤ 0.0007.

(E) REACTOME_PI_3K_CASCADE_FGFR1 (r = 0.86, $R^2$ = 0.73). O in Y versus O Iso: −0.79 years; Kruskal–Wallis P = 0.0032. Y in O grafts did not differ from Y Iso.

(F, G) REACTOME_FGFR2_LIGAND_BINDING_AND_ACTIVATION (F; r = 0.69, $R^2$ = 0.48, P = $9.86 \times 10^{-13}$) and REACTOME_SHC_MEDIATED_CASCADE_FGFR4 (G; r = 0.89, $R^2$ = 0.79). O in Y versus O Iso: −0.78 to −0.82 years; Kruskal–Wallis P ≤ 0.0025.

PathAge was computed from DNAm at CpGs in each pathway's gene regulatory domains. Left, offset-corrected PathAge against chronological age in the lifespan series (2 weeks to 17 months), colored by age; dashed line, y = x; MAE and RMSE are computed against y = x. Right, ΔPathAge relative to the mean of Y Iso (dashed line at 0) in unilateral transplantation groups. Box plots show the median and interquartile range; points are individual samples. Pairwise comparisons use Welch's t-tests with BH correction across five planned contrasts per pathway. Offset correction and Y Iso referencing are constant shifts within each pathway.

**Extended Data Figure 19 | PI3K–mTOR, nutrient-sensing and lipid-metabolism PathAge clocks**

(A) HALLMARK_PI3K_AKT_MTOR_SIGNALING (n = 80; slope = 0.56, $R^2$ = 0.54, MAE = 0.22 years, r = 0.74, P = $5.63 \times 10^{-15}$). O in Y versus O Iso: −0.46 years; Kruskal–Wallis P = 0.0018.

(B–D) REACTOME_AMINO_ACIDS_REGULATE_MTORC1 (B), REACTOME_MTOR_SIGNALLING (C) and REACTOME_MTORC1_MEDIATED_SIGNALLING (D); r = 0.51–0.75, P ≤ $1.54 \times 10^{-6}$. O in Y versus O Iso: −0.80 to −0.99 years; Kruskal–Wallis P ≤ 0.0113.

(E, F) KEGG_FATTY_ACID_METABOLISM (E; r = 0.82, $P < 2 \times 10^{-16}$) and REACTOME_MITOCHONDRIAL_FATTY_ACID_BETA_OXIDATION_OF_SATURATED_FATTY_ACIDS (F; r = 0.63, P = $2.55 \times 10^{-10}$). O in Y versus O Iso: −0.67 to −0.69 years; Kruskal–Wallis P ≤ 0.0014.

(G, H) REACTOME_HS_GAG_BIOSYNTHESIS (G) and KEGG_TERPENOID_BACKBONE_BIOSYNTHESIS (H); r = 0.66–0.86, P ≤ $2.86 \times 10^{-11}$. O in Y versus O Iso: −0.74 to −0.97 years; Kruskal–Wallis P ≤ 0.0131.

Plots, reference lines and tests are as in Extended Data Fig. 18.

**Extended Data Figure 20 | Inflammatory, vesicular, extracellular-matrix and cell-cycle PathAge clocks**

(A–D) REACTOME_INTERFERON_GAMMA_SIGNALING (A), REACTOME_INTERLEUKIN_18_SIGNALING (B), HALLMARK_INTERFERON_ALPHA_RESPONSE (C) and KEGG_JAK_STAT_SIGNALING_PATHWAY (D); r = 0.61–0.81, $P \leq 2.71 \times 10^{-9}$. O in Y versus O Iso: −0.30 to −1.11 years; Kruskal–Wallis P ≤ 0.0198.

(E, F) KEGG_SNARE_INTERACTIONS_IN_VESICULAR_TRANSPORT (E) and REACTOME_DISINHIBITION_OF_SNARE_FORMATION (F); r = 0.75–0.84, $P \leq 1.31 \times 10^{-15}$. O in Y versus O Iso: −0.84 to −1.37 years; Kruskal–Wallis P ≤ 0.0058.

(G–I) REACTOME_CELL_SURFACE_INTERACTIONS_AT_THE_VASCULAR_WALL (G), REACTOME_COLLAGEN_CHAIN_TRIMERIZATION (H) and REACTOME_CROSSLINKING_OF_COLLAGEN_FIBRILS (I); r = 0.65–0.74, $P \leq 9.47 \times 10^{-11}$. O in Y versus O Iso: −0.74 to −1.40 years; Kruskal–Wallis P ≤ 0.0024.

(J, K) REACTOME_MITOTIC_G2_G2_M_PHASES (J) and KEGG_TIGHT_JUNCTION (K); r = 0.83, $P < 2 \times 10^{-16}$. O in Y versus O Iso: −0.39 to −1.73 years; Kruskal–Wallis P ≤ 0.0192.

Plots, reference lines and tests are as in Extended Data Fig. 18.

**Extended Data Figure 21 | DNA methylation trajectories at ovarian cell-type marker loci**

(A–E) DNAm (β value) against age at the most age-correlated CpG annotated to each marker gene, for marker genes of oocytes (A; Gdf9, Padi6, Zar1), cumulus cells (B; Ereg, Rgs2, Pgr), theca cells (C; Hsd3b1, Nr5a1, Ptch1), macrophages (D; Cd68, Adgre1, Lyz2) and surface epithelium (E; Epcam, Krt19, Krt8).

Panel titles give the gene and probe, the directions of age-associated expression and methylation change, the number of samples (n = 80–90), and Pearson r and P for methylation against age. Points are ovarian samples; lines show linear fits. Cell-type labels annotate bulk-tissue measurements.

**Extended Data Figure 22 | Histology and transcriptomic age of multinucleated giant cell-rich regions**

(A) Representative H&E-stained sections of young and old ovaries from sham, isochronic, contralateral control and heterochronic (Y in O, O in Y) groups; insets enlarge the boxed regions.

(B) Left, schematic of the laser-capture microdissection and RNA-seq workflow used to generate the profiles of MNGC regions, macrophages and stroma (GSE283393; Converse *et al*, 2025). Right, OvaAge (scaled elastic net) for these profiles.

(C–E) Multi-tissue clocks (Tyshkovskiy et al, 2026): chronological (C; mouse, YuGene, elastic net), mortality (D; rodents and primates, log10 hazard ratio) and normalized tAge (E; % of lifespan).

(F) Standardized differences in module-specific mortality tAge (tAge Δ), MNGC versus stroma.

(G, H) Standardized effects of KEGG (G) and Hallmark (H) gene sets on mortality tAge, MNGC versus stroma; bar color shows the enrichment score.

(I) Gene-level contributions to the mortality-tAge difference (log10 hazard ratio), MNGC versus stroma; color shows the direction of expression change.

Box plots show the median and interquartile range; points are individual samples. Pairwise P values are shown above brackets. Error bars show s.e. ^P < 0.1, *P < 0.05, **P < 0.01, ***P < 0.001.

**Extended Data Figure 23 | Pathway and senescence-associated expression in MNGC, macrophage and stromal captures**

(A, B) Hallmark (A) and KEGG (B) GSEA of MNGC captures versus macrophages and versus stroma (GSE283393; Converse *et al*, 2025). Positive NES indicates enrichment in MNGCs; asterisks show BH-adjusted significance.

(C) Mean PROGENy activity scores per capture type; asterisks show one-way ANOVA significance.

(D) Mean z-scores for six senescence and senescence-associated secretory phenotype (SASP) gene sets (SenMayo, Core SASP, Fridman UP, Reactome SASP, WikiPathways SASP and CellAge inducers); asterisks show Tukey HSD comparisons.

*$P < 0.05$, **$P < 0.01$, ***$P < 0.001$.

**Extended Data Figure 24 | Projection of a mouse MNGC expression signature into human ovarian nuclei**

(A) PCA of variance-stabilized mouse laser-capture profiles of MNGC regions, macrophages (Mac) and stroma (Str) (GSE283393; Converse *et al*, 2025).

(B) Differential expression of MNGCs versus stroma (left) and versus macrophages (right), used to define the MNGC signature.

(C) UMAP of human ovarian single-nucleus RNA-seq (GSE202601; Jin *et al*, 2025), colored by annotated cell type.

(D) Expression of selected immune markers across human immune clusters.

(E) UMAP feature plots of selected immune-cell markers.

(F) Re-clustered immune nuclei (Leiden clusters, left) and per-nucleus MNGC signature scores (right).

(G) MNGC signature scores across the full human dataset.

**Extended Data Figure 25 | Metabolic gene-set enrichment in MNGC captures**

(A–D) GSEA of Recon3D subsystems (A), MitoCarta 3.0 pathways (B), KEGG metabolic pathways (C) and scFEA-defined metabolic modules (D) for MNGC captures versus macrophages and versus stroma (GSE283393; Converse *et al*, 2025).

Positive NES indicates enrichment in MNGCs. ***FDR < 0.001, **FDR < 0.01, *FDR < 0.05, +FDR < 0.25. Panel D shows enrichment of module-associated genes.

**Extended Data Figure 26 | Plasma metabolite signatures after heterochronic parabiosis compared with external datasets**

(A–F) Metabolite log2 fold changes compared between pairs of signatures: O Het vs O Iso against GHRKO (A), Y Het vs Y Iso against GHRKO (B), O Het vs Y Het against GHRKO (C), Y Het vs Y Iso against caloric restriction (D), aging (O Iso vs Y Iso) against O Het vs O Iso (E), and Y Het vs Y Iso against O Het vs Y Het (F). Spearman $\rho = 0.472, 0.482, -0.582, 0.293, -0.420$ and $-0.621$, with 19, 19, 19, 67, 113 and 128 matched metabolites. Red dashed lines are linear fits; green backgrounds mark $P < 0.05$.

(G) Clustered log2 fold changes of metabolites shared between the parabiosis (PB) comparisons and external aging and intervention datasets; colors are clipped at ±3. Missing measurements are blank.

GHRKO, growth hormone receptor knockout; Het, heterochronic; Iso, isochronic. Some comparisons share samples or control groups, which can contribute to correlations between them.
